# Inhibition of the macrophage demethylase LSD1 reverses *Leishmania amazonensis*-induced transcriptomic changes and causes a decrease in parasite load

**DOI:** 10.1101/2023.09.29.560133

**Authors:** Maria Gutiérrez-Sanchez, Adina A. Baniahmad, Charfeddine Gharsallah, Sheng Zhang, Suzanne Lamotte, Karin Schmidtkunz, Hugo Varet, Rachel Legendre, Nicolás Tráves, Florent Dingli, Damarys Loew, Dante Rotili, Sergio Valente, Antonello Mai, Philippe M. Loiseau, Sébastien Pomel, Manfred Jung, Hervé Lecoeur, Eric Prina, Gerald F. Späth

## Abstract

Intracellular pathogens exploit host cell functions to favor their own survival. In recent years, the subversion of epigenetic regulation has emerged as a key microbial strategy to modify host cell gene expression and evade antimicrobial immune responses. Using the protozoan parasite *Leishmania* as a model system, we have recently demonstrated that infection causes histone H3 hypomethylation, which is associated with the establishment of an anti-inflammatory phenotype, suggesting that host cell demethylases may play a role in the intracellular survival of these parasites. In this study, we combined pharmacological inhibition with RNA sequencing and quantitative immune-precipitation analysis to investigate the role of the macrophage lysine demethylase LSD1 (KDM1a) in *Leishmania* intracellular infection *in vitro*. Treatment of infected macrophages with validated, LSD1-specific inhibitors resulted in a significant reduction in parasite burden. We confirmed the impact of these inhibitors on LSD1 activity within macrophage nuclear extracts using an *in vitro* demethylase assay and established their LSD1 target engagement *in situ* by cellular thermal shift assay. RNA-seq analysis of infected and inhibitor-treated macrophages linked parasite killing to a partial reversion of infection-dependent expression changes, restoring the macrophage anti-microbial response and limiting cholesterol biosynthesis. While we ruled out any impact of *Leishmania* on LSD1 expression or localization, we uncovered significant alterations in LSD1 complex formation within infected macrophages, involving unique interactions with host cell regulatory proteins such as Rcor-1. Our study sheds important new light on the epigenetic mechanisms of macrophage immuno-metabolic subversion by intracellular *Leishmania* and identifies LSD1 as a potential candidate for host-directed, anti-leishmanial therapy.

## INTRODUCTION

Many viral, bacterial and eukaryotic pathogens infect mammalian cells and have co-evolved strategies to modulate the gene expression profiles of their host cells and promote their intracellular survival (Benoit et al., 2008; Herbein and Varin, 2010; Herbert et al., 2004; Kamhawi and Serafim, 2020; Noel et al., 2004; Pearce and MacDonald, 2002; Raes et al., 2007). This co-evolution is well illustrated for protozoan parasites of the genus *Leishmania* that proliferate inside sand fly insect vectors as motile promastigotes, and as non-motile amastigotes inside fully acidified phagolysosomes of mammalian macrophages. *Leishmania* subverts the immunological and metabolic functions of these important and highly toxic immune cells by interfering with cell signaling (Gregory and Olivier, 2005), metabolism (Rabhi et al., 2012) or epigenetic control (Afrin et al., 2019; Kamhawi and Serafim, 2020; Lecoeur et al., 2020). The host consequences of *Leishmania* immune subversion are often devastating causing severe immune pathologies termed ‘leishmaniases’ that range from self-healing cutaneous to lethal visceral forms (Conceicao-Silva and Morgado, 2019; McGwire and Satoskar, 2014).

Despite the severe consequences to human health, the mechanisms underlying the subversion of macrophage functions by intracellular *Leishmania* remain poorly understood. Epigenetic control of host immuno-metabolic processes in infected macrophages and dendritic cells has recently emerged as a new parasite strategy for host cell subversion (Kamhawi and Serafim, 2020; Lecoeur et al., 2022a; Lecoeur et al., 2020). First, *Leishmania* has been shown to manipulate host cell miRNAs to limit pro-inflammatory gene expression (Ganguly et al., 2022; Lemaire et al., 2013; Mukherjee et al., 2015; Muxel et al., 2018; Nandan et al., 2021; Tiwari et al., 2017). Second, intracellular *Leishmania* infection has been correlated to changes in the macrophage DNA methylation pattern, influencing host cell signalling pathways, oxidative phosphorylation, and cell adhesion (Marr et al., 2014). Finally, we and others have shown that *Leishmania* infections alters host cell histone modification, such as acetylation or methylation, whose levels are regulated by writer enzymes (e.g. acetyltransferases, methyltransferases) and eraser enzymes (i.e. deacetylases, demethylases). Our recent work has established a first link between a macrophage epigenetic eraser activity and *Leishmania* infection: applying a ChIPqPCR approach we provided first evidence that *L. amazonensis* infection causes hypo-methylation at pro-inflammatory gene promoters at activating histone H3K4 and repressing H3K9 marks, which correlated with a corresponding decrease in transcript output (Kamhawi and Serafim, 2020; Lecoeur et al., 2020). These H3 modifications are both removed by the Lysine-Specific Demethylase 1 (LSD1, a.k.a. KDM1a) (Shi et al., 2004), which in accordance to a given lysine mark can increase or decrease gene expression.

LSD1 is composed of three domains: the catalytic AO domain, the Swi3/Rcs8/Moira (SWIRM) domain and the Tower domain (Burg et al., 2015). Aside its enzymatic function, it carries important scaffolding functions via the Tower domain that serves as binding site for the CoREST transcription repressor complex (essential for LSD1 to associate with the nucleosome and exert its demethylase activity) and the SWIRM domain that functions as a platform for protein-protein interactions (Burg et al., 2015; Laurent and Shi, 2016). LSD1 is involved in diverse biological processes including differentiation (Whyte et al., 2012), autophagy (Ambrosio et al., 2019) or metabolic processes (Sakamoto et al., 2015). Conceivably, deregulation of LSD1 functions causes severe human diseases, including cancer (Majello et al., 2019) and, as a consequence, represents one of the prime therapeutic targets, with various compounds undergoing clinical testing for treating small lung cancer cells (SCLC) or acute myeloid leukaemia (Fang et al., 2019). Surprisingly, despite the availability of LSD1-specific inhibitors and the known role of LSD1 to control macrophage polarization, its potential as a host-directed target for anti-microbial intervention has not been explored yet, except for viral infections (Mazzarella et al., 2021; Zwergel et al., 2018). We previously proposed an experimental framework for the discovery of host-directed inhibitors targeting macrophage histone-modifying enzymes (Lamotte et al., 2019; Lamotte et al., 2017) for which large compound libraries are available given current efforts to cure non-communicable human diseases such as cancer, obesity, auto-immunity by restoring a normal epigenetic profile (Hogg et al., 2020; Nebbioso et al., 2018).

Here, we established a first proof-of-principle for host-directed, anti-leishmanial therapy by exploring the possibility to target LSD1 for drug discovery. Screening a diverse epigenetic inhibitor library against intracellular and extracellular *L. amazonensis* parasites identified a series of anti-parasitic compounds designed to target mammalian LSD1 (https://cordis.europa.eu/project/id/602080/fr). We demonstrate that LSD1 targeting compounds partially revert infection-induced expression changes that correlate with reduced parasite survival and growth, and provide first evidence that *L. amazonensis* subverts the LSD1 scaffolding functions to establish a macrophage phenotype permissive for intracellular infection.

## MATERIAL AND METHODS

### Ethics statement

Six-week-old female C57BL/6 and Swiss *nu/nu* mice were purchased from Janvier (Saint Germain-sur-l’Arbresle, France). All animals were housed in A3 animal facilities according to the guidelines of Institut Pasteur and the “Comité d’Ethique pour l’Expérimentation Animale” (CEEA) and protocols were approved by the “Ministère de l’Enseignement Supérieur, Direction Générale pour la Recherche et l’Innovation” under number 2013-0047. EP is authorized to perform experiments on vertebrate animals (license 75–1265) issued by the ‘‘Direction Départementale de la Protection des Populations de Paris’’ and is responsible for all the experiments conducted personally or under his supervision as governed by the laws and regulations relating to the protection of animals.

### Bone marrow-derived macrophages and cell line cultures

Bone marrow cell suspensions were recovered from tibias and femurs of female C57Bl/6JRj mice in Dulbecco’s phosphate buffered solution (PBS), and cultured in complete medium containing 4 g/L glucose, 1 mM pyruvate and 3.97 mM L-Alanyl-L-Glutamine and supplemented with 10% heat-inactivated fetal calf serum (FCS), streptomycin (50 µg/mL), and penicillin (50 IU/mL) (de La Llave et al., 2011). Briefly, one million cells per ml of complete medium supplemented with 50 ng/ml of mouse recombinant colony-stimulating factor 1 (rm-CSF1, ImmunoTools) were incubated in bacteriological Petri dishes (Greiner bio-one 664161) at 37°C in a 7.5% CO_2_ atmosphere. After 6 days of culture, the medium was removed and adherent cells were incubated with pre-warmed PBS pH 7.4 containing 25 mM EDTA for 30 min at 37°C. Bone marrow-derived macrophages (BMDMs) were detached by gentle flushing, collected, and resuspended in complete medium supplemented with 30 ng/ml of rmCSF-1. BMDMs were seeded in 100 mm tissue culture dishes (Petri BD Falcon, 353003) for nuclear or total protein extraction, and RNA isolation, and in 384-well tissue culture-treated flat-optically clear microplates (CellCarrier™ plate, PerkinElmer) for High Content Assay (HCA).

### Parasite isolation and promastigote differentiation

mCherry*-*transgenic, tissue-derived amastigotes of *Leishmania amazonensis* strain LV79 (WHO reference number MPRO/BR/72/M1841) and *Leishmania donovani* strain 1S2D (WHO reference number MHOM/SD/62/1S-CL2D) were isolated from infected footpads of Swiss nude mice (Lecoeur et al., 2020) and infected spleen of golden Syrian hamsters, respectively. Promastigotes were differentiated from animal-derived amastigotes and maintained at 27°C in M199 medium supplemented with 10% FCS, 20 mM HEPES pH 6.9, 4 mM NaHCO3, 2 mM glutamine, 20 μM folic acid, 100 μM adenine, 13.7 µM hemin, 8 μM 6-biopterin, 100 U ml^-1^ of penicillin, and 100 μg ml^-1^ of streptomycin. Promastigotes were maintained in culture by dilution in fresh medium upon reaching stationary phase and were used in the Log growth phase for the Dye Reduction Assay (DRA).

### Chemical compounds

All small chemical compounds used in screening assays were part of the EU FP7 A-ParaDDisE project (https://cordis.europa.eu/project/id/602080/fr) and represent potential or validated epigenetic inhibitors. Inhibitors of lysine specific demethylase 1, including tranylcypromine (TCP), a 4-phenylbenzamide TCP derivative MC2652, and a heterobenzoylamino TCP derivative MC4030 were synthesized following the procedure described by Fioravanti and colleagues (Fioravanti et al., 2022). The synthesis of hydrophobic Tags is described in supplementary information.

### Cell infection and treatment

After seeding at day 6 of differentiation, BMDMs were left for 5 hours at 37°C to attach to their new culture support. BMDMs were infected with lesion-derived *L. amazonensis* amastigotes at a multiplicity of infection (MOI) of 4 and then cultured at 34°C. LSD1*i* were added to uninfected and *Leishmania-* Infected Macrophages (LIMs) 24 hours after infection, at a final concentration of 10 µM (HCA, epifluorescence microscopy) or 15 µM (WB, CETSA).

### High Content Assay

The High Content Assay (HCA) was used to assess the anti-leishmanial activity against intramacrophagic amastigotes of *L*. *amazonensis* (Lamotte et al., 2019). 425 compounds (A-ParaDDisE libraryhttps://cordis.europa.eu/project/id/602080/fr) were added one day after macrophage infection at 10 μM final concentration (1% DMSO final concentration). Controls included a vehicle control (DMSO), a leishmanicidal control (amphotericin B), and a macrophage cytotoxic control (cycloheximide). Each compound was tested in quadruplicates. Briefly, after 3 days of co-incubation with compounds, BMDM nuclei and parasitophorous vacuoles (PVs) were stained with Hoechst 33342 (Invitrogen Molecular Probes, 10 µM) and LysoTracker Green (Life Technologies, DND-26, 1 µM), respectively. Acquisition of images was performed on live cell cultures using the OPERA QEHS device with a 10x air objective (NA 0.4) using the green (488 nm), the blue (405 nm) and the red (561 nm) channels for the detection of PVs, nuclei and amastigotes, respectively (Lamotte et al., 2019). Images were transferred to the Columbus Conductor™ Database (Perkin Elmer Technologies) and analysed using the integrated Image analysis building blocks (Lamotte et al., 2019). Quantitative values obtained for host cell and parasite numbers were exported to Excel and SigmaPlot for further analysis and graphical representations.

### Nuclear and cytoplasmic protein extraction

For western blotting and LSD1 Activity Assay, nuclear and cytoplasmic extracts were obtained after 2 days of infection. The medium was removed from each dish, and cells were gently washed with PBS at room temperature (RT). PBS was then replaced by 1 ml of Lysis Buffer (LBI, Table S2).

After 10 min of incubation on ice, cells were detached with a cell scraper and collected in 2 ml Eppendorf tubes. The cell homogenate was passed 5 times through a 27 G needle to isolate nuclei. The quality of isolated nuclei was assessed by the EVOS Cell Imaging System and by Western blotting (Figure S6). Samples were centrifuged for 10 min at 4°C at 250 g for activity assay, and Western blots. Supernatants containing cytoplasmic proteins were removed and kept for further analysis, whereas 150 µl of Extraction Buffer (EBI, Table S2) were added to pellets. Tubes were vortexed and incubated for 10 min at 4°C. Sonication was performed using the Diagenode’s Bioruptor® Standard for 5 min at high power (30” on/30” off sonication cycles). Extracts were cleared by centrifugation at 15,000g for 10 min at 4°C.

### Analysis of LSD1 enzymatic activity

LSD activity assay way performed with the fluorometric KDM1/LSD1 activity quantitation kit (ABCAM) according to the manufacturer’s instructions. The activity was measured with 1 mg nuclear protein extracts from uninfected BMDMs and dimethylated lysine 4 of the Histone H3 (H3K4me2) as substrate.

### Cellular Thermal Shift Assay (CETSA)

Uninfected and infected BMDMs were treated or not at 1 day after infection with 15 µM LSD1*i* (MC2652, MC4030, and TCP) or 0.1% DMSO as control. After one day of treatment, cells were washed with the pre-warmed medium before adding 1.5 ml of cold Extraction Buffer (1x PBS – 1x PIC – 1 mM PMSF). Then, cells were collected with a cell scraper in two 1.5 ml Eppendorf tubes and subjected to three cycles of freezing in liquid nitrogen and thawing in a thermomixer at 25°C with a 400 rpm agitation step. Lysates were submitted to 3 sonication cycles (30” on/30” off) at high power, then pooled in one Eppendorf tube per condition and centrifuged at 20,000 g for 20 min at 4°C.

Supernatants containing soluble proteins were transferred to 1.5 ml tubes, then distributed into twelve 0.2 ml PCR tubes and heated to different temperatures (from 37°C to 70°C, ΔT°=+3°C) in a Thermocycler (Applied ProFlex ™ 96 Well PCR System) for 3 min followed by cooling at 4°C. Afterwards, protein solutions were transferred to 1.5 ml Eppendorf tubes and cleared at 20,000 g for 20 min at 4°C. Carefully, 75 µL of supernatant containing the soluble protein fraction was collected in a new LoBind® Eppendorf tube.

The detection of the remaining soluble proteins was performed by Western Blot, where 11 µL of each cleared sample were mixed with 2 µL of NuPAGE® Reducing Agent and 5 µL of NuPAGE® LDS Sample Buffer. The samples for each CETSA melt curve were loaded on the same gel. The quantification was performed by measuring the fluorescence intensity emitted by the ECL Plex fluorescent secondary antibodies with the Amersham™ ImageQuant™ 800. Normalization was carried out with the Amersham ImageQuant TL analysis software considering the band intensity of the sample heated to 37°C as 100%. Melting curves were generated with GraphPad Prism software package.

### Western blotting

Experiments were performed on 5-10 μg proteins. Proteins were resolved by SDS–PAGE (4–12% Bis-Tris NuPAGE gels) in MOPS buffer and electroblotted onto polyvinylidene difluoride (PVDF) membranes, that were blocked with 2% Amersham ECL Prime blocking agent in 1x PBS containing 0.25% Tween 20. Membranes were then probed overnight at 4°C with the following primary antibodies: rabbit anti-LSD1 (C69G12, Cell Signaling Technology), rabbit anti-MAO-A (10539-1-AP, Proteintech), mouse anti-GAPDH (60004-1-Ig, Proteintech), rabbit anti-RCOR1 (27686-1-AP), mouse anti-YY1 (66281-1-Ig, Proteintech), rabbit anti-Histone-H3 (17168-1-AP, Proteintech).

Following incubation, either with Amersham ECL Plex Cy3 or Cy5 conjugated antibodies or with the appropriate peroxydase conjugate secondary antibodies (Thermo Scientific), fluorescence or luminescence (SuperSignal West Pico reagent, ThermoFisher Scientific) were measured in an Amersham ImageQuant 800 Imaging System (Cytiva). Relative protein expression was calculated by densitometric analysis using the Amersham ImageQuant analysis software. For every band, the ratio between the values obtained for the target protein and the YY1 (nuclear extracts) and the GAPDH (cytoplasmic extracts) normalization controls were calculated. Fold changes were calculated using the control sample as calibrator, control values of uninfected and unstimulated samples being set to 1. Modulations of the LSD1 protein content were statistically determined by the nonparametric Wilcoxon rank-sum test using the GraphPad Prism 7.03 software.

### Determination of LSD1 localization by epifluorescence microscopy analysis

2x10^5^ BMDMs were seeded in 500 µl of complete medium in 24-well plates containing sterile glass coverslips and were infected or not with *L. amazonensis* amastigotes (MOI = 4). At day day 3 PI, cells were fixed in 4% paraformaldehyde (PFA) for 30 min at RT after a PBS wash (1 ml/well, 10 min, RT). Subsequently, quenching solution (50 mM ammonium chloride) was added to block free aldehyde groups (1 ml/well, 15 min, RT). Cells were rinsed twice in PBS and incubated with the Blocking and permeabilization solution for 1 hour at RT (5% donkey normal serum, 0.3% Triton X-100, 5% goat normal serum, in PBS). Primary antibody solution (α-LSD1, C69G12, Cell Signaling, 1/100), Mouse immune serum (1/1000) for parasite detection, 1% Bovine Serum Albumin, 0.3% Triton X-100 in PBS ) was added for an overnight incubation at 4°C. The next day, cells were rinsed 3 times in 1x PBS and subsequently incubated for 90 min at RT in the dark with the secondary antibody solution (AlexaFluor 594 Goat α-mouse Jackson ImmunoResearch, 350µg/ml), FITC Donkey α-rabbit (Jackson ImmunoResearch, 750µg/ml), 1% BSA, 0.3% Triton X-100, PBS). Afterwards, cells were rinsed once in 1x PBS, incubated for 15 min in a 1x PBS solution containing Hoechst 33342 (1mM) and rinsed one last time with 1x PBS for 5 min. Finally, coverslips were rinsed in Milli-Q water, dried and mounted using Slowfade Gold mounting agent on clean glass slides. The slides were sealed with transparent varnish. The slides were examined, and images were captured using an inverted confocal microscope Leica SP8. To generate representative images from our immunofluorescence datasets, we used the Z-Projection function in FIJI (ImageJ). Confocal image stacks were processed using the Stacks → Z-Projection tool, selecting Maximum Intensity Projection as the projection method.

### Determination of parasite load in live macrophages by epifluorescence imaging

Parasite load was quantified by measuring mCherry fluorescence in live infected macrophages. Macrophage Nuclei were stained with 5 mM Hoechst 33342 (ThermoFisher Scientific) for 15 min at room temperature. Live-cell images were acquired on an EVOS M5000 Imaging System 4 days post-infection, following 3 days of treatment, using the DAPI light cube for Hoechst detection (Ex357/44 nm, Em 447/60 nm) and the Texas red light cube for mCherry detection (Ex 585/29 nm, Em 624/40 nm). Five fields were imaged with a 20x objective. Image analysis was performed using the ImageJ software package on a minimum of 100 cells. Parasite load was quantified as the Raw Integrated Density of mCherry fluorescence in macrophages in the absence or presence of LSD1*i*. Statistical analyses (Wilcoxon-Mann-Whitney test) and data visualization were performed in the GraphPad Prism software package.

### Viability assay for *Leishmania* promastigotes and amastigotes

Anti-leishmanial activity of compounds on free parasites was evaluated using an adapted resazurin-based dye reduction (DRA) assay with cell-cycling promastigotes from logarithmic growth phase maintained in M199 supplemented medium.

Fixed concentrations or serial dilutions of the inhibitors were done in triplicate or quadruplicate, at 27°C for promastigotes and 34°C for amastigotes in 96-well plates or 384-well plates. DMSO vehicle and AmB were used as controls. Two days later, resazurin was added (10 μL per well for a final concentration of 5 µg/ml) and resorufin-derived fluorescence intensity was measured 24 h later using a Tecan Safire 2 reader (Ex 558[±[4.5[nm, Em 585[±[10[nm). Following background subtraction (complete parasite culture medium with resazurin in absence of parasites), data were expressed as percentages of growth compared to DMSO-treated controls (POC).

### RNA extraction

Large RNA above 200 nucleotides were isolated from macrophage samples using the NucleoSpin miRNA kit (Macherey-Nagel) according to the manufacturer’s instructions. Evaluation of RNA quality was carried out by optical density measurement using a Nanodrop device (Kisker, http://www.kisker-biotech.com). RNAs were isolated from 3 independent biological replicates of uninfected and *Leishmania*-infected macrophages treated or not with LSD1*i* MC4030 for 24 hrs. RNA isolation was performed at 2 days post-infection.

### RNA-Seq analysis

Library preparation and sequencing was performed at the Biomics platform of Institut Pasteur. Briefly, DNAse-treated RNA extracts were processed for library preparation using the Truseq Stranded mRNA sample preparation kit (Illumina, San Diego, California) according to the manufacturer’s instructions. An initial poly (A)+ RNA isolation step (included in the Illumina protocol) was performed with 1 µg of large RNA to isolate the mRNA fraction and remove ribosomal RNA. The mRNA-enriched fraction was fragmented by divalent ions at high temperatures. The fragmented samples were randomly primed for reverse transcription followed by second-strand synthesis to create double-stranded cDNA fragments. No end repair step was necessary. An adenine was added to the 3’-end and specific Illumina adapters were ligated. Ligation products were submitted to PCR amplification. The quality of the obtained libraries was controlled using a Bioanalyzer DNA1000 Chips (Agilent, # 5067-1504) and quantified by spectrofluorimetric analysis (Quant-iT™ High-Sensitivity DNA Assay Kit, #Q33120, Invitrogen).

Sequencing was performed on the Illumina Hiseq2500 platform to generate single-end 65 bp reads bearing strand specificity. Reads were cleaned using cutadapt version 1.11 and only sequences at least 25 nt in length were considered for further analysis. STAR version 2.5.0a (Dobin et al., 2013), with default parameters, was used for alignment on the reference genome (GRCm38 from Ensembl database 94). Genes were counted using featureCounts version 1.4.6-p3 (Liao et al., 2014) from the Subreads package (parameters: -t gene -s 0). Transcriptomic data are made publicly available at the NCBI’s Gene Expression Omnibus repository (Superseries ongoing submission).

### LSD1 target genes

A list of LSD1 target genes was compiled from bibliographic analyses and presented in Table S3. Target genes were evidenced after LSD1 pharmacological inhibition (Augert et al., 2019; Barth et al., 2019; Egolf et al., 2019; Hoshino et al., 2019; Maiques-Diaz et al., 2018), LSD1 Knocking Out (Barth et al., 2019; Jin et al., 2013), and siRNA LSD1 knockdown (Chen et al., 2017; Jin et al., 2017) using transcriptomic (Augert et al., 2019; Barth et al., 2019; Chen et al., 2017; Egolf et al., 2019; Hoshino et al., 2019; Jin et al., 2013), proteomic (Jin et al., 2017) and Chip Seq (Augert et al., 2019; Chen et al., 2017; Egolf et al., 2019; Jin et al., 2013; Jin et al., 2017; Maiques-Diaz et al., 2018) analyses.

### Gene ontology analysis

Ensembl gene identifiers were translated into ENTREZ gene identifiers prior to the enrichment analysis. ENTREZ genes linked to several Ensembl IDs were associated with the one having the highest expression level. Functional gene-set enrichment analysis was performed using the Fisher statistical test for the over-representation of differentially expressed genes (gene-level adjusted P-value lower than 5%). Several types of gene-sets have been tested:

i. Hallmark (https://doi.org/10.1016%2Fj.cels.2015.12.004),
ii. reactome (https://doi.org/10.1093/nar/gkab1028),
iii. KEGG pathways (https://doi.org/10.1093/nar/gkac963),
iv. GO terms (https://doi.org/10.1093/genetics/iyad031),
v. Wikipathways (https://doi.org/10.1093/NAR/gkaa1024). Only gene-sets with a false discovery rate (FDR) lower than 0.05 were considered significantly enriched in differentially expressed gene sets.

For the analysis of epigenetic regulatory factors, the differential expression values, expressed as log2 fold changes (log2 FC), were computed for each gene, and were visually depicted as a network using the Cytoscape software package (version 3.10.0). Each node within this network projects the log2FC information. Protein-protein interactions were extracted from the STRING Database (version 5.0) (https://string-db.org/). These interactions were filtered based on confidence scores, with a minimum interaction score of 0.700, thus generating an interaction network grounded in known or predicted associations among these genes. Gene Ontology (GO) terms we also incorporated for each network node, facilitating functional annotations pertaining to biological processes, molecular functions, and cellular components.

### Chromatin extraction

Chromatin extraction was performed two days after infection. The medium was removed from each dish, and 10 ml of fixing solution (1% paraformaldehyde in serum-free medium) was added and incubated at room temperature for 5 minutes. Then, 1 ml of quenching solution (1.375 M glycine) was added. The solutions were removed, and the fixed cells were washed twice with 10 ml of ice-cold 1× PBS. After removing the PBS, 2 ml of lysis buffer (50 mM HEPES pH 7.5, 140 mM NaCl, 1 mM EDTA, 10% glycerol, 0.5% NP-40, 0.25% Triton, 1 mM PMSF, and protease inhibitor cocktail) were added, and the samples were incubated for 10 minutes with rocking on ice. Cells were collected with a scraper, transferred to Eppendorf tubes, passed through a syringe with a 27G needle, and sonicated (using Diagenode’s Bioruptor Standard) for 3 cycles at low power (30 s on/30 s off) to better isolate the nuclei. They were then centrifuged at 500 g for 10 minutes at 4 °C. This procedure was repeated until the nuclei preparations were free of debris or “parasite ghosts”. Clean nuclei were centrifuged at 500 g each time, and the pellet was resuspended in 150 µl per tube of sonication buffer (10 mM Tris-HCl pH 8, 100 mM NaCl, 1 mM EDTA, 0.5 mM EGTA, 0.1% sodium deoxycholate, 0.5% N-lauroylsarcosine, 1 mM PMSF, and protease inhibitor cocktail), vortexed, and sonicated for 15 cycles (30 s on/30 s off) to shear the chromatin. Samples were then cleared at 16,000 g for 10 minutes at 4 °C. Sheared chromatin showed fragments between 200 and 500 bp.

### LSD1 co-immunoprecipitation assay

Rapid immunoprecipitation mass spectrometry of endogenous proteins (RIME) for analysis of chromatin complexes (described in DOI: 10.1038/nprot.2016.020) was performed to identify LSD1 partners in uninfected and infected BMDMs, with slight modifications. Briefly, the assay was carried out using five biological replicates of chromatin samples from uninfected and infected BMDMs. Chromatin samples were pre-cleared for 1h with rotation at 4°C using Dynabeads™ Protein G (Invitrogen). The flow-through was then recovered and divided into two tubes per sample. To the first tube, 2 µl of the negative control rabbit IgG pool (Diagenode C15410206) were added, and to the second tube, 10 µl of anti-LSD1 (C69G12, Cell Signaling). Samples were incubated overnight at 4°C with rotation. The following day, Dynabeads were washed with blocking solution (0.5% BSA in 1× PBS) before being added to the tubes containing chromatin plus antibodies. After addition, tubes were incubated for 2h at 4°C with rotation. The supernatant was removed, and the Dynabeads–antibody–antigen complexes were subjected to ten washes in RIPA buffer. Subsequently, they were washed four times with 100 mM ammonium bicarbonate and sent for mass spectrometry analysis.

### LC-MS/MS Analysis

Online chromatography was performed with an RSLCnano system (Ultimate 3000, Thermo Scientific) coupled to an Orbitrap Eclipse mass spectrometer (Thermo Scientific). Peptides were trapped on a 2 cm nanoviper Precolumn (i.d. 75 μm, C18 Acclaim PepMap^TM^ 100, Thermo Scientific) at a flow rate of 3.0 µL/min in buffer A (2/98 MeCN/H2O in 0.1% formic acid) for 4 min to desalt and concentrate the samples. Separation was performed on a 50 cm nanoviper column (i.d. 75 μm, C18, Acclaim PepMap^TM^ RSLC, 2 μm, 100Å, Thermo Scientific) regulated to a temperature of 50°C with a linear gradient from 2% to 25% buffer B (100% MeCN in 0.1% formic acid) at a flow rate of 300 nL/min over 91 min. MS1 data were collected in the Orbitrap (120,000 resolution; maximum injection time 60 ms; AGC 4 x 10^5^). Charges states between 2 and 7 were required for MS2 analysis, and a 60 sec dynamic exclusion window was used. MS2 scan were performed in the ion trap in rapid mode with HCD fragmentation (isolation window 1.2 Da; NCE 30%; maximum injection time 35 ms; AGC 10^4^)

### Proteomics data processing

For identification, the data were searched against the *Mus musculus* (UP000000589_10090) and the *Leishmania amazonensis* MHOMBR71973M2269 in the TriTrypDB (release 60) databases using Sequest HT through proteome discoverer (version 2.4). Enzyme specificity was set to trypsin and a maximum of two miss cleavages sites were allowed. Oxidized methionine, Met-loss, Met-loss-Acetyl and N-terminal acetylation were set as variable modifications. Maximum allowed mass deviation was set to 10 ppm for monoisotopic precursor ions and 0.6 Da for MS/MS peaks. The resulting files were further processed using myProMS v3.10.0 (https://github.com/bioinfo-pf-curie/myproms) (Poullet et al., 2007). FDR calculation used Percolator (The et al., 2016) and was set to 1% at the peptide level for the whole study. The label free quantification was performed by peptide Extracted Ion Chromatograms (XICs), reextracted across all conditions and computed with MassChroQ version 2.2.21 (Valot et al., 2011). For protein quantification, XICs from proteotypic and non-proteotypic peptides were used by assigning to the best protein, peptides shared by multiple match groups, and missed cleavages, charge states and sources were allowed. Label-free quantification (LFQ) was performed following the algorithm as described (Cox et al., 2014).

The final LFQ intensities were used as protein abundance and further analyzed using myProMS. High-confidence LSD1-interacting proteins were identified by applying the following criteria: a fold-change ≥4 in LSD1 IP versus IgG control, adjusted p-value ≤0.01, at least three unique peptides per protein, and detection in at least four biological replicates.

## RESULTS

### *L. amazonensis* subverts the macrophage epigenetic landscape during infection

We apply a multidisciplinary approach to (i) identify potential epigenetic pathways affected by *Leishmania* infection, (ii) reveal their relevance for intracellular parasite survival, and (iii) assess the mechanisms underlying their subversion. We first conducted an RNA-Seq analysis of Bone Marrow-Derived Macrophages (BMDMs) left uninfected (UI) or infected with virulent *Leishmania amazonensis* amastigotes (*Leishmania-* Infected Macrophages, LIMs) at day 2 post-infection (PI) (Figure 1A), which revealed significant expression changes for 5742 genes (Figure S1A1 and S1B1, adjusted p value < 0.05) affecting the macrophage innate immune response, including antigen presentation and toll receptor signaling cascade, as well as metabolic processes such as cholesterol biosynthesis (Figures S2-5). To gain insight into epigenetic subversion strategies, we more specifically investigated the impact of infection on the expression of macrophage Epigenetic Regulatory Factors (EpiRFs), including histone / DNA modifying enzymes, associated transcription factors, and members of epigenetic complexes. From 787 EpiRFs obtained from manual curation of publicly available databases including EpiFactors (http://epifactors.autosome.org., Table S1), 315 showed significant changes in transcript abundance during infection, with increased expression observed for 154 (e.g. *Dnmt1*, FC = +1.39; *Dot1l*, FC = +2.78), and decreased expression affecting 161 (e.g. *Sirt3*, FC = -1.46; *Ehmt,*: FC = -1.38) (Figure 1B). Overall, 39.7 % of EpiRFs were modulated by the parasite, revealing epigenetic regulation as a key target for parasite subversion of host cell immune-metabolomic functions.

**Figure 1:**
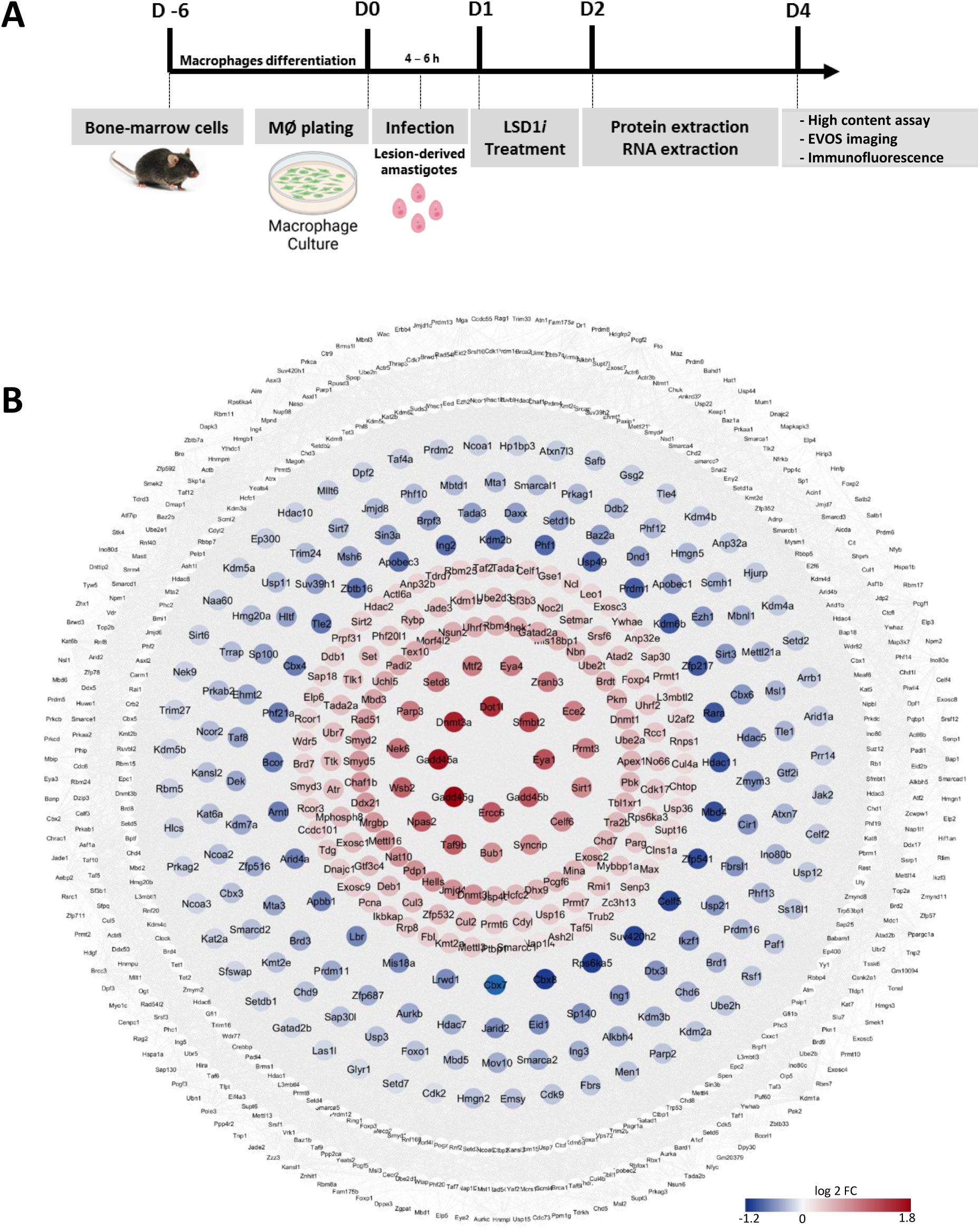
Experimental pipeline (A) and differential expression analysis (B) of Epigenetic Regulatory Factors (EpiRFs) in *Leishmania-*infected macrophages (LIMs). BMDMs were infected with lesion-derived, mCherry-transgenic amastigotes of *L. amazonensis* and compared to uninfected BMDMs by RNA-Seq analysis two days after infection. Out of 787 EpiRFs, 315 were modulated in LIMs compared to uninfected BMDMs. Constitutive (white), downregulated (blue) and upregulated (red) genes are visualized as STRING interaction networks using the the Cytoscape V3.10.0 software package. Log_2_ Fold Change (Log_2_ FC) for differentially expressed EpiRF genes with an adjusted p Value cutoff of 0.05 are shown.

### Pharmacological interference with macrophage epigenetic regulation reduces burden of intracellular *L. amazonensis*

We next used a pharmacological approach to assess the relevance of this epigenetic subversion strategy for intracellular *L. amazonensis* survival. We assessed 425 epigenetic inhibitors provided by the FP7 A-PARADDISE project, which were designed to target various classes of mammalian epigenetic regulators, including sirtuins, Histone Acetyl Transferases (HATs), Histone Deacetylases (HDACs), Histone Methyl Transferases (HMT), Histone DeMethylases (HDMs) and DNA Methyl Transferases (DNMTs) (see https://cordis.europa.eu/project/id/602080). We screened this library by monitoring the intracellular amastigote survival in *L. amazonensis*-infected BMDMs using a high content assay, and extracellular parasite survival using a resazurin-based viability assay applied on promastigotes in culture (Lamotte et al., 2019). This dual strategy allowed us to distinguish anti-leishmanial compounds that affect parasite viability directly (killing both extracellular and intracellular parasites) from compounds that affect parasite viability indirectly via host-dependent mechanisms (killing exclusively intracellular parasites) (Lower right quadrant, Figure 2A).

**Figure 2:**
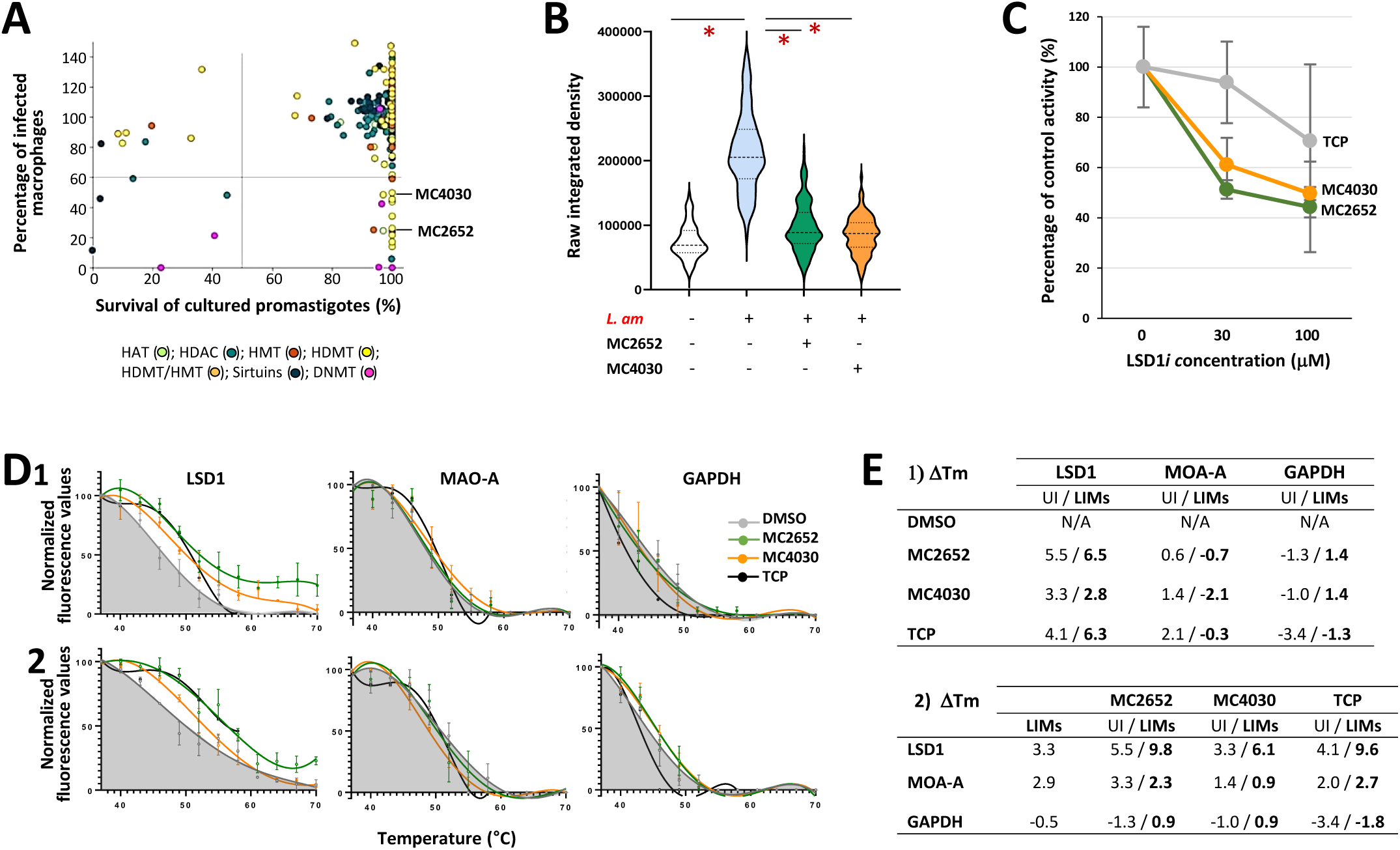
Identification and validation of LSD1 inhibitors with host-dependent, anti-leishmanial activity. **(A) Biparametric dot plot**. The graph shows the survival of *L. amazonensis* parasites after three days of treatment at 4[µM for cultured extracellular promastigotes (X-axis) and at 10 µM for intracellular amastigotes in LIMs (Y-axis) compared to the DMSO control. Data were manually curated to remove compounds toxic for macrophages. Quadrant lines divide the high-low range antileishmanial activity into four sections. Potential host-directed compounds are located in the right lower quadrant. Different classes of inhibitors are represented by different colors. **(B) Violin plot of anti-leishmanial activity for two selected LSD1*i* (MC2652 and MC4030).** Inhibitors were added at a concentration of 10 μM on BMDMs one day post-infection with lesion-derived mCherry-transgenic *L. amazonensis* amastigotes. Epifluorescence microscopy was performed after three days of treatment. Violin plots display the quantification of mCherry fluorescence per cell for each sample in uninfected BMDMs (white), untreated LIMs (grey) and compound-treated LIMs (MC2652, green; MC4030, orange). Raw Integrated Density per cell was determined by ImageJ on 100-180 cells. The p-value is indicated (*: p<0.0001). **(C) LSD1 activity assay.** LSD1 demethylase activity was assessed in 1 µg of nuclear protein extracts from uninfected BMDMs in the absence or presence of increasing concentrations of the LSD1*i* MC2652, MC4030 and TCP. The percentage of inhibition of LSD1 activity was calculated for the two indicated inhibitor doses compared to untreated BMDMs set at 100%. **(D) Cellular thermal shift assay (CETSA).** Uninfected BMDMs (D1) and LIMs (D2) at day 1 post-infection were treated for 24 hours with 1% DMSO as control (grey area), and with 15 µM MC2652 (green line), MC4030 (orange line), and TCP (black line). The results are presented as a percentage of the fluorescent signal detected at the lowest temperature in each melting curve, with values representing the mean and SEM of 2-3 biological replicates. **(E) Table summarizing the thermal shifts observed in the presence of LSD1 inhibitors.** Note that the thermal shifts are specific for LSD1. Thermal shift values were calculated by subtracting the melting temperature (Tm) of the UI sample from the Tm of the sample of interest.

Of particular interest were the 18 compounds that caused significant decrease in intracellular but not extracellular parasite survival. Such putative host-dependent mode of action has been observed for different classes of targets including HMTs, HDMTs and DNMTs, suggesting that interference with macrophage epigenetic regulation both at DNA and histone levels can interfere with intracellular parasite growth or trigger parasite death. Most effective were a series of inhibitors directed against the Lysine-Specific Demethylase 1 (LSD1, a.k.a. KDM1A). This enzyme demethylates histone H3 at K4 and K9 residues, which has been linked to macrophage polarization and the transcription of pro-inflammatory cytokines (Janzer et al., 2012; Tan et al., 2019). The anti-leishmanial effect of these LSD1 inhibitors resonates well with our previous data that correlated *L. amazonensis* infection with macrophage H3K4 and H3K9 hypomethylation (Lecoeur et al., 2020), suggesting that this demethylase may be key for parasite-mediated, epigenetic host cell subversion.

We next focused our attention on two derivatives of the canonical flavoenzyme monoamine oxidase A inhibitor Tranylcypromine (TCP), i.e. the compounds MC2652 and MC4030. These inhibitors were synthesized by replacing the 3-phenyl ring of TCP with distinct heterocyclic thiophene moieties, and display an increased LSD1 inhibitory effect in cancer cell lines (Fioravanti et al., 2022). Surprisingly, the original LSD1 inhibitor TCP did not affect intracellular parasite survival even though it significantly interfered with LSD1 activity using a biochemical assay (data not shown and Figure 2C). In contrast, the more bulky TCP-derivatives MC2652 and MC4030 not only interfered with LSD1 activity but also strongly reduced burden of both intracellular *L. amazonensis* and *L. donovani* as judged by epifluorescence microscopy and (Figures 2B, C and S6).

To determine *in cellula* whether this inhibition could be attributed to target engagement - i.e. direct binding to LSD1, we performed CETSA. The binding of the LSD1*i* to the demethylase was confirmed by LSD1 thermal stabilization in the presence of the inhibitors (Figure 2D and 2E and Figure S9). Importantly, this interaction was specific to LSD1, as no thermal shift was observed for MAO-A or GAPDH in either uninfected (Figure 2D, and 2E) or *L. amazonensis*-infected BMDMs (Figure 2D1 and 2D2, and Figure S12A and S12B).

Unfortunately, all our attempts to phenocopy this pharmacological effect on intracellular parasites by using three independent si-RNAs targeting LSD1/KDM1A failed as reduction of mRNA abundance neither affected protein abundance nor parasite burden (Figures S7A and S8A). To establish an independent validation of the link between inhibition of LSD1 activity and reduced parasite burden, we therefore synthesized a series of novel LSD1 inhibitors with a different structure compared to MC2652 and MC4030. These novel, irreversible compounds are based on a tranylcypromine-derived warhead bearing hydrophobic substituents (e.g., Adamantyl, heptafluorobutyl or other fluorinated groups) (see Figures S9-10). Such hydrophobically tagged compounds (short HyTs) have the capacity to trigger the degradation of targets recruited in the context of a proteolytic chimera by activating quality control pathways (Xie et al., 2023). The AAB-020-derived HyTs bind LSD1, as shown by Cellular Thermal Shift Assay (CETSA) (Figure S11), and specifically inhibit its catalytic activity (Table S3). While neither AAB-020 nor its derived HyT compounds showed direct toxicity toward isolated parasites (amastigotes or promastigotes), only the bulky HyTs reduced the parasite load in LIMs (Figure S10A2, B2). This leishmanicidal effect was not accompanied by LSD1 degradation as judged by Western blot analysis (data not shown), suggesting that these HyTs likely act through target engagement and functional inhibition, possibly by altering LSD1 scaffolding functions. Because MC2652 and MC4030 are easier to synthesize than the HyTs, we pursued subsequent experiments with these compounds.

### *L. amazonensis* affects LSD1 complex formation

Surprisingly, even though our screening data suggest an important role of LSD1 in supporting intracellular *Leishmania* survival, infection itself neither modulated LSD1 protein levels during the first three days of infection (Figure 3A), nor affected LSD1 nuclear localization (Figure 3B). Given the above observed inhibition profile, and since LSD1 activity relies on its scaffolding function interacting with key partners in regulatory complexes (Burg et al., 2015), we next analyzed expression changes of 90 known LSD1-associated proteins during *L. amazonensis* infection mining our RNA-Seq data (Figure 3C). While no significant transcriptional modulation was observed for LSD1 itself, 35 of the LSD1-associated proteins (35.4%) were modulated (Figure 3C), including 19 showing decreased mRNA abundance (e.g. *Zfp217*, FC = -1.87; *Suv39h1*, FC = +1.40), and 16 showing increased mRNA abundance (e.g. *Mbd3*, FC = +1.48; *Gatad2a*, FC = +1.43). Thus, *L. amazonensis* infection changes transcript abundance of known LSD1 interaction partners, possibly subverting the composition of LSD1 complexes (Table 1).

**Figure 3.**
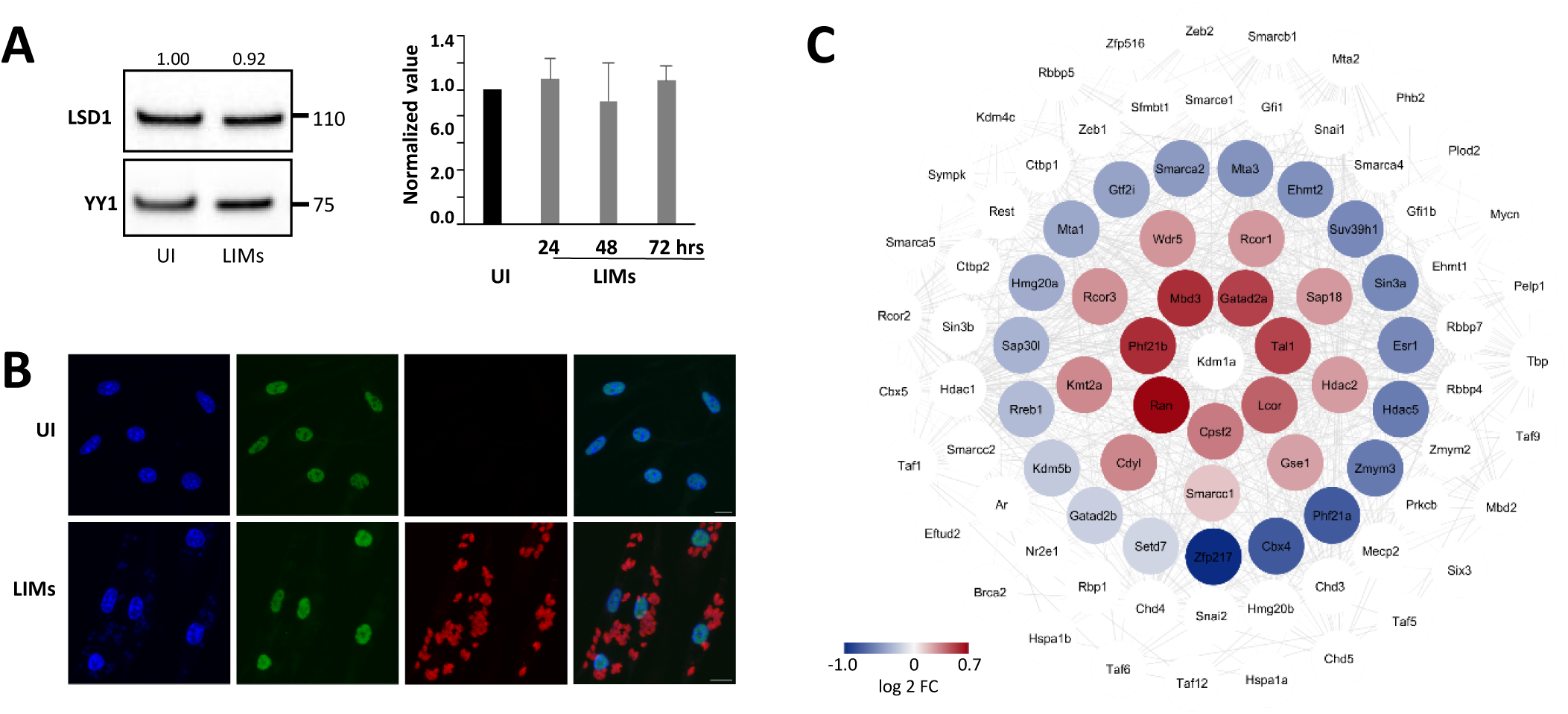
Impact of *L. amazonensis* infection on LSD1 expression, localization and expression of interaction partners. (**A**) **Western Blot analysis. Nuclear proteins were extracted from uninfected (UI) BMDMs and LIMs at day 2 post-infection.** One representative WB analyzing the abundance of LSD1 and YY1 at 2 days PI is shown (left panel). Signals from up to 5 independent replicates were quantified (right panel) by normalizing LSD1 against the YY1 loading control and setting the value for UI BMDMs to 1. (**B**) **Epifluorescence analysis of LSD1 localization in fixed cell samples.** Macrophage nuclei were stained with Hoechst 33342, parasites were visualized by their mCherry fluorescence (red channel), and LSD1 was detected by immunostaining (green channel). Z-projection (max intensity) are shown for each channel and superposed ones. (**C**) **Expression changes observed for LSD1 interaction partners.** BMDMs were infected with lesion-derived mCherry-transgenic *L. amazonensis* amastigotes and compared to UI BMDMs. RNA-Seq analysis of infected and uninfected BMDMS was performed as indicated in legend of Figure 1. Changes in RNA abundance for LSD1 and its known interaction partners are visualized as STRING interaction networks. Differentially expressed genes are defined by a significant modulation, p<0.05). Constitutive transcript abundance, white; decreased transcript abundance, blue; increased transcript abundance, red.

**Table 1.**
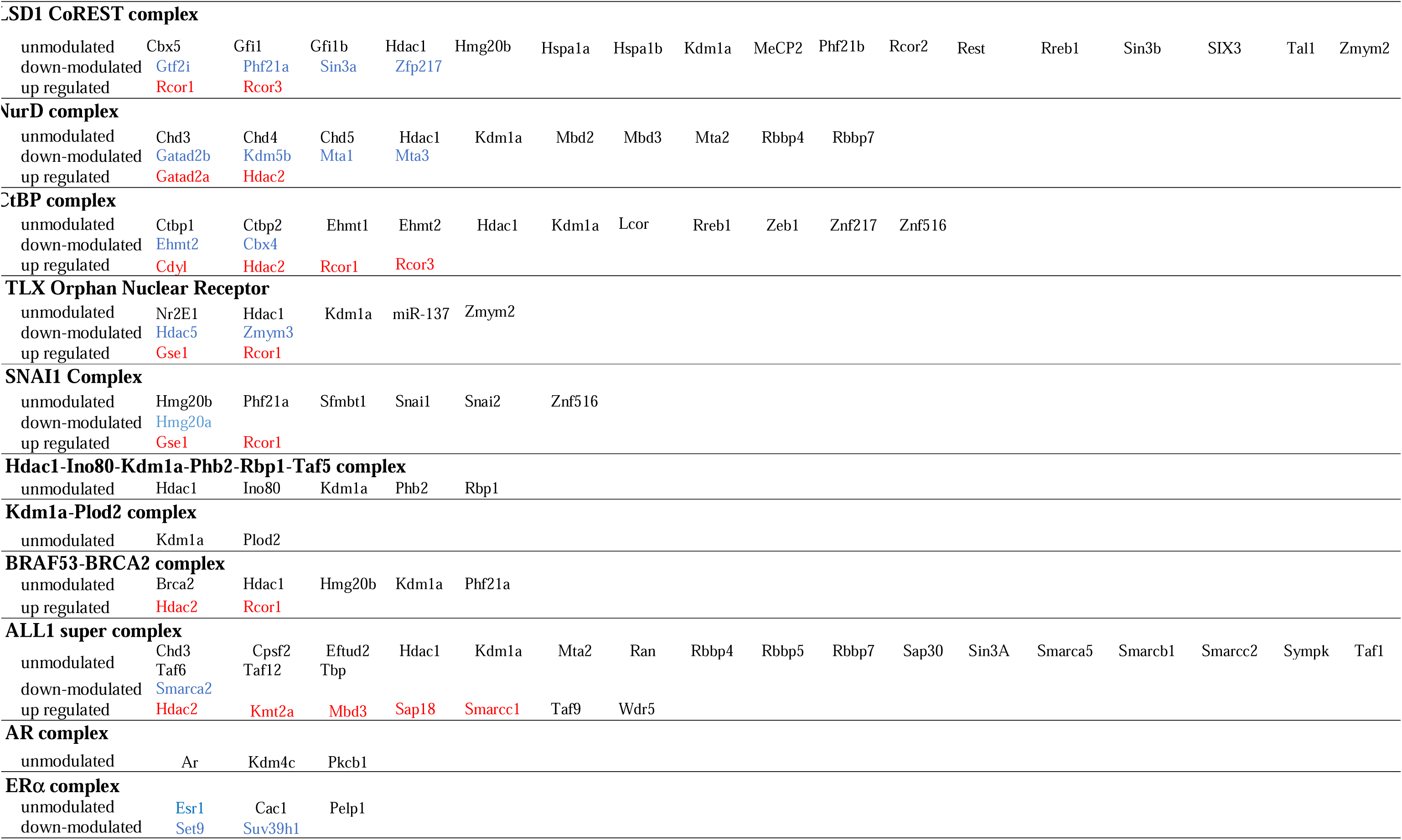
Gutierrez Sanchez et al. : Transcriptional modulation of genes involved in the composition of the main LSD1 complexes.

**Table 2.** Gutierrez Sanchez et al. *Leishmania* LSD1 proteins interactors, co-immunoprecipitated with the specific anti-LSD1 antibody Ab1 using LIMs nuclear protein extracts.

|  |  |
| --- | --- |
| **LAMA_000014200.1** | Tubulin, putative |
| **LAMA_000030000.1** | Ribosomal protein S7e, putative |
| **LAMA_000050800.1** | Hypothetical protein, conserved |
| **LAMA_000069400.1** | Hypothetical protein, conserved |
| **LAMA_000111100.1** | Histone H2A, putative |
| **LAMA_000154400.1** | Core histone H2A/H2B/H3/H4, putative – Annotated as Histone H2B in *L. mexicana* |
| **LAMA_000206700.1** | T-complex protein 10 C-terminus, putative |
| **LAMA_000224800.1** | C2 domain containing protein, putative |
| **LAMA_000261800.1** | Hypothetical protein, conserved |
| **LAMA_000365000.1** | Hypothetical protein, conserved - Annotated as KHARON in *L. mexicana* |
| **LAMA_000388500.1** | Hypothetical protein, conserved |
| **LAMA_000424900.1** | Hypothetical protein, conserved |
| **LAMA_000472900.1** | Hypothetical protein, conserved |
| **LAMA_000475000.1** | Ribosomal proteins L26 eukaryotic, putative |
| **LAMA_000489100.1** | Hypothetical protein, conserved |
| **LAMA_000541300.1** | Hypothetical protein, conserved |
| **LAMA_000574600.1** | Leucine Rich repeat, putative |
| **LAMA_000582300.1** | Hypothetical protein, conserved |
| **LAMA_000615100.1** | Glyceraldehyde 3-phosphate dehydrogenase, putative |
| **LAMA_000622200.1** | Hypothetical protein, conserved |
| **LAMA_000657900.1** | Hypothetical protein, conserved |
| **LAMA_000685500.1** | Leucine-rich repeat/Glycogen recognition site of AMP activated protein kinase, putative |
| **LAMA_000721100.1** | Hypothetical protein, conserved - Annotated as KHassociated protein 1 in *L. mexicana* |
| **LAMA_000780200.1** | TATA box binding protein associated factor (TAF) putative - Annotated as Histone H4 in *L. mexicana* |

Using CETSA, we observed a marked shift in LSD1 thermal stability by 3.3°C in infected versus uninfected BMDMs (Figures 4A), providing initial, indirect evidence that LSD1-interactions and -containing complexes indeed may be remodeled upon infection. To directly assess infection-dependent changes in LSD1 complex composition, we performed LSD1 co-immunoprecipitation (co-IP) using chromatin samples from uninfected and infected BMDMs at day 2 post-infection.

**Figure 4:**
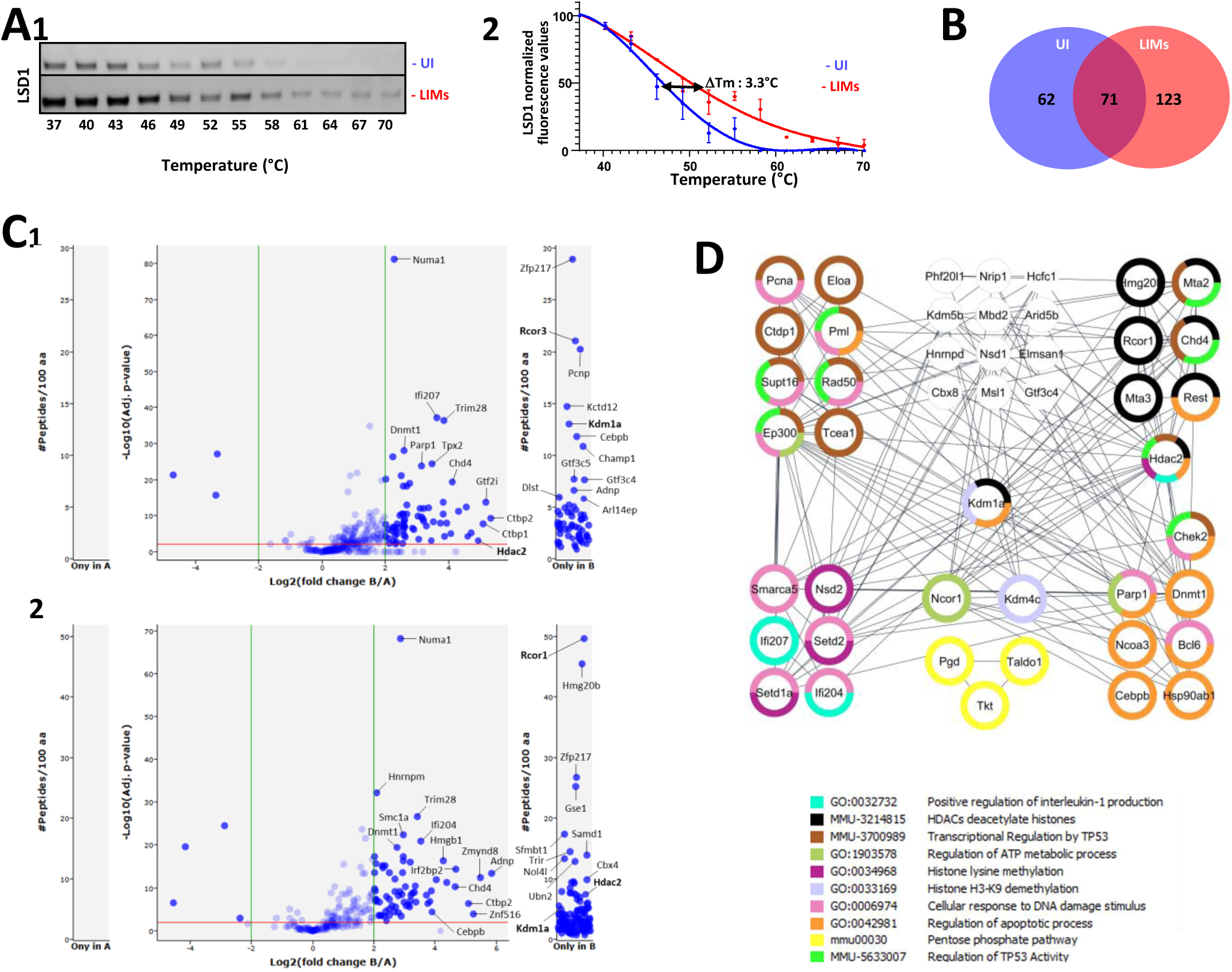
Analyses of LSD1 interactions. **(A) Evidence of a LSD1-specific thermal stabilization in LIMs by CETSA analysis**. Representative Western blot analyses comparing uninfected (UI) BMDMs and LIMs at day 2 PI in presence of DMSO (A 1). Melting curves for both samples are shown with the temperature shift indicated (A2). **(B) Number of identified LSD1-interacting proteins, evidenced after immunoprecipitation in UI BMDMs and LIMs**. Number of identified LSD1-interacting proteins, evidenced after immunoprecipitation in UI BMDMs and LIMs. Chromatin samples were isolated from UI BMDMs and LIMs at day 2 PI. LSD1-containing protein complexes were immunoprecipitated using an anti-LSD1 antibody (n = 5 biological replicates per condition). As a negative control, IP was performed in parallel with rabbit IgGs to account for non-specific binding. Proteins were identified and quantified by mass spectrometry using the following filtering criteria: fold change ≥ 4 (LSD1-specific vs. IgG control), adjusted p-value ≤ 0.01, ≥ 3 unique peptides per protein, detected in at least 4 biological replicates. **(C) Identification of LSD1 interactors in uninfected (UI) BMDMs (C1) and LIMs (C2).** Volcano plots obtained using myProMS software for the protein enrichment in uninfected BMDMs (C1) and LIMs (C2) after LSD1 immunoprecipitation. The X-axis corresponds to the H/L ratio of quantified proteins in log2 scale while the Y-axis corresponds to the p-value for the protein H/L ratio determined using the different H/L peptides ratios for this protein. Vertical green lines correspond to the 2-fold change in protein expression level. Left (Only in A) and right panels (Only in B) correspond to the protein only detected in control IgG and LSD1 immunoprecipitation, respectively. For some dots, protein names are indicated including LSD1 (Kdm1a), Rcor1, and Rcor3. **(D) Gene ontology enrichment analysis of STRING networks identified between LSD1 interaction partners in LIMs using the Cytoscape V3.10.0 software package.** Interactions were retrieved from STRING using a confidence score ≥ 0.4 and visualized in Cytoscape. Each node represents a protein, and colored rings indicate membership in significantly enriched GO biological processes or Reactome pathways. Colors correspond to the following functional categories. Edges represent predicted or experimentally supported functional interactions derived from STRING.

Relative to non-specific IgG controls, we identified 133 and 194 high-confidence LSD1-interactors in uninfected BMDMs and LIMs, respectively (Figure 4C). Seventy-one proteins were shared between conditions, including LSD1/KDM1A itself and the established interactor DNMT1 (Figure 4D). Sixty-two interactors were unique to uninfected BMDMs, whereas 123 were unique to LIMs. LIM-specific partners were enriched for chromatin-regulatory activities, such as histone lysine methylation, HDAC-mediated deacetylation, and regulation of TRP53-dependent transcription, as well as regulators of the the pentose phosphate pathway and apoptotic processes, both known to be modulated in LIMs (Lecoeur et al., 2022b; Zhang et al., 2024).

### LSD1 inhibition counteracts parasite-driven expression changes in LIMs

In order to elucidate host cell pathways that may govern the host-directed, anti-leishmanial effect of compound MC4030, we analyzed the impact of drug treatment on uninfected and infected BMDMs (Figure S1A2 and B2). As noted above, *L. amazonensis* infection alone had a profound effect on the macrophage transcriptome (see Figure 1B and Figure S1A1 and B1), affecting many cellular pathways, including protein kinase activity, sterol biosynthesis, and responses to lipopolysaccharide, hypoxia, and oxidative stress (Figures S2-5). Among the 5742 modulated transcripts, 1101 (19.17%) were previously reported to be regulated by LSD1 (Figure 5A1), which further supports our hypothesis that *Leishmania* targets LSD1 to establish permissive conditions for intracellular macrophage infection.

**Figure 5:**
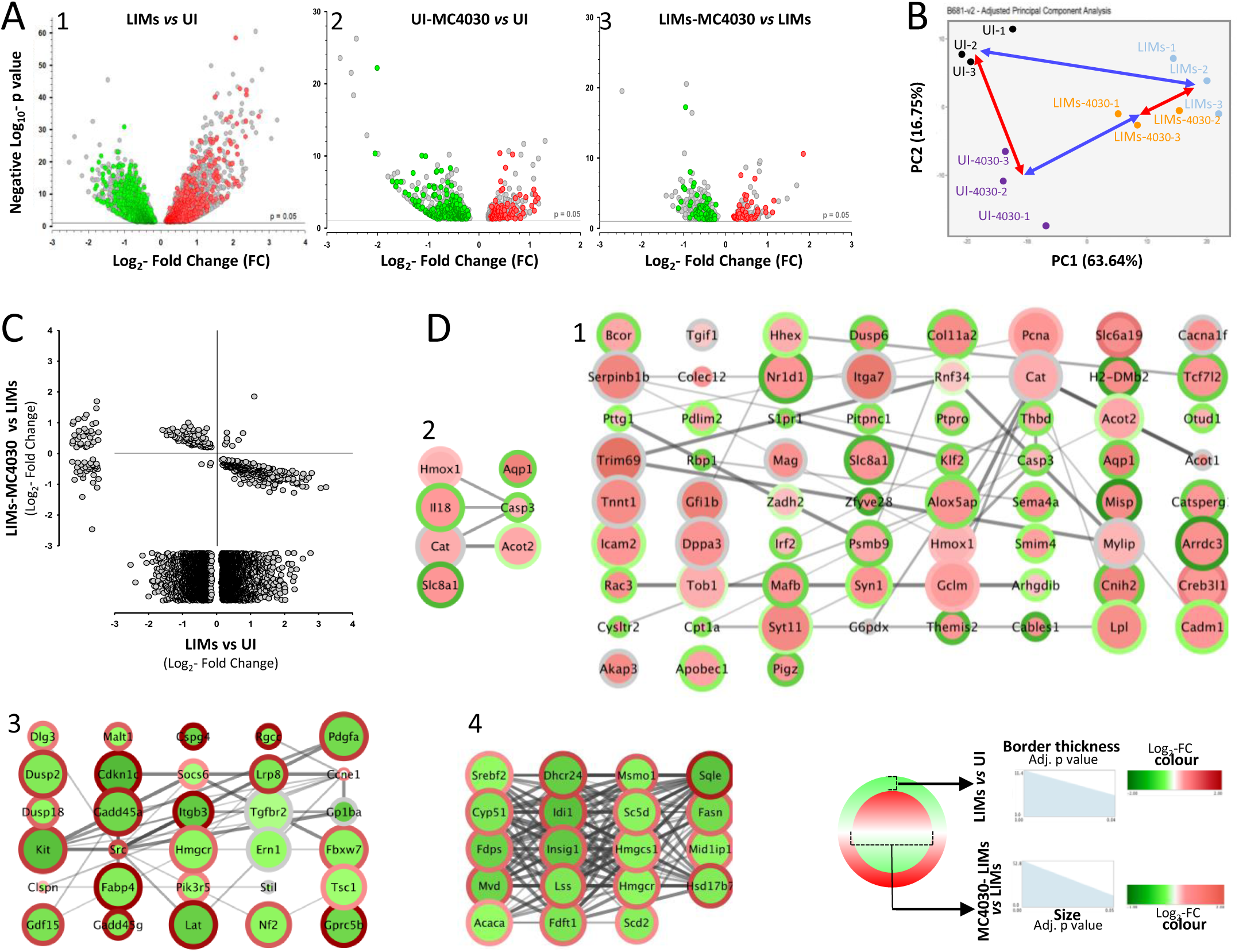
Transcript profiling of uninfected (UI) BMDMs and LIMs at day 2 post-infection treated or not for 1 day with MC4030. **(A) Volcano plots showing differential transcript abundance (adj. p values < 0.05) for the comparisons LIMs vs UI BMDMs (left), MC4030-treated vs UI BMDMs (middle), and MC4030-treated LIMs vs untreated LIMs (right)**. Colored dots correspond to transcripts whose expression is controlled by LSD1. Green, decreased mRNA abundance in a given comparisons; red, increased mRNA abundance in a given comparison. **(B) PCA analysis of the different replicates for each condition.** Arrows indicate the distance between experimental groups. **(C) Double ratio plot displaying the log2 FC ratio of transcript expression in LIMs vs UI BMDMs (X-axis) and MC4030-treated LIMs vs LIMs (Y-axis).** **(D) Visual representation of reversible gene expression changes for genes that share the Gene Ontology terms ‘Response to lipopolysaccharide’ (1), ‘Response to hypoxia’ (2), ‘Regulation of protein kinase activity’ (3), and ‘Steroid biosynthetic process’ (4).** The color of the node border and node filling respectively shows expression changes for the comparisons LIMs *vs* UI BMDMs and M4030-treated LIMs *vs* LIMs, with the border thickness and the node size corresponding to the p-value as indicated in the legend.

MC4030 treatment of uninfected BMDMs resulted in expression changes of 1097 genes, 287 of which are regulated by LSD1 (26.1%) (Figure 5A2). These results further support the specificity of MC4030 to target macrophage LSD1 and sustains a host-directed mechanism for its anti-leishmanial activity. Finally, MC4030 treatment of infected BMDMs affected the abundance of 456 transcripts, 117 (25.7%) of which are regulated by LSD1 (Figure 5A3). MC4030 treatment seems to counteract the infection-induced phenotype as judged by PCA analysis of the three experimental groups, which shows a shorter distance between MC4030-treated infected and uninfected BMDMs compared to the distance between infected and uninfected BMDMs in absence of the inhibitor (Figure 5B). This is further confirmed by the double ratio plot shown in Figure 5C, providing direct evidence that 95% of expression changes caused by the treatment do counteract infection-dependent expression changes observed in infected BMDMs. For example, many genes implicated in the macrophage response to lipopolysaccharide and hypoxia showed reduced expression during infection, which was restored after drug treatment (Figure 5D1 and D2). Conversely, the increased expression of genes implicated in signal transduction and steroid biosynthesis observed in infected BMDMs was significantly reduced upon treatment (Figure 5D3 and D4). Importantly, this reversion was not just the consequence of parasite elimination but a direct effect of LSD1 inhibition given that the treatment had only a slight effect on parasite burden at the time point chosen for the RNA-Seq analysis (i.e. 48h of treatment).

In conclusion, our data provide evidence that MC4030 kills intracellular *Leishmania* by restoring a macrophage expression profile that interferes with the parasite survival by rescuing the host cell immune response and limiting the parasite’s access to nutrients essential for its proliferation.

## DISCUSSION

*Leishmania* infection of macrophages is associated with significant changes in gene expression that contribute to the creation of a favorable environment for parasite survival and replication within host cells. Here we provide pharmacological evidence that *Leishmania* exploits the host demethylase LSD1, raising questions on how *Leishmania* engages with LSD1, how this engagement changes the host cell phenotype, and how interfering with this engagement leads to parasite killing.

*Leishmania* can engage with LSD1 to modulate one or both of its biological functions, i.e. its enzymatic activity and / or its scaffolding properties. The absence of LSD1 modulation with respect to transcript abundance, protein abundance or subcellular localization levels suggested that LSD1 subversion relies on mechanisms independent of its expression, such as complex formation. This hypothesis was confirmed by CETSA analyses that provided first evidence that *Leishmania* infection increases LSD1 thermal stability. This increase likely reflects changes in the nature of LSD1 host complexes. Unlike bacterial ‘nucleomodulins” that can directly engage with nuclear host cells components to subvert their functions (Bierne and Cossart, 2012), no *Leishmania* proteins were co-purified with nuclear LSD1.

Infection-dependent LSD1 interactions directly inform on possible mechanisms of immune subversion. LSD1 exhibits differential cofactor associations in *Leishmania*-infected vs. uninfected macrophages: In LIMs, LSD1 specifically interacts with RCOR1, whereas in uninfected BMDMs, it preferentially binds RCOR3. The RCOR1-LSD1 complex may recruit HDAC2, thereby erasing permissive histone marks (Rivera et al., 2022), a mechanism that could account for the reduced histone acetylation levels we previously observed in LIMs (Lecoeur et al., 2020). Beyond its epigenetic role, HDAC2-mediated repression has been shown to dampen the apoptosis process (Kramer, 2009), a hallmark of LIMs (Fernandes and Zamboni, 2024; Lecoeur et al., 2022b).

LSD1 is an epigenetic factor that positively regulates LPS responses, enhancing pro-inflammatory signaling (Sobczak et al., 2021; Tokarz et al., 2019; Wojtala et al., 2021). However, *L. amazonensis* infection strongly dampens LPS responsiveness in macrophages (Lecoeur et al., 2020). Our transcriptomic data suggest that parasite-driven subversion of LSD1 interactions - shifting from RCOR3 to RCOR1 – may be a key mechanism to impair LPS-induced transcriptional responses and inhibit microbicidal host responses. Our data show that reversion of anti-inflammatory gene expression by pharmacological inhibition of LSD1 rescues macrophage anti-microbial gene expression, which then causes the observed reduction in parasite burden.

In conclusion, our results shed new light on a *Leishmania*-specific epigenetic subversion strategy that targets the LSD1 scaffolding function of the host cell to change transcriptional complexes and establish an infection-permissive phenotype. This strategy was demonstrated in acute myeloid leukemia where the interaction between LSD1 and RCOR1 with the SNAG-domain transcription repressor GFI1 can be dispersed by TCP-derived inhibitors allowing myeloid differentiation (Maiques-Diaz et al., 2018). The identification of distinct, LSD1-targeting compounds with host-directed, anti-leishmanial activity pharmacologically validates host cell epigenetic regulation and downstream-regulated pathways as a fertile ground for the discovery of macrophage drug targets that impact parasite survival, which may be broadly applicable to other intracellular pathogens known to target host cell epigenetic regulation for their survival, such as *Mycobacterium tuberculosis* (Pennini et al., 2006), *Listeria monocytogenes* (Hamon and Cossart, 2008) or *Toxoplasma gondii* (Schneider et al., 2013).

## ACKNOWLEDGMENTS

With financial support from “la Région Île-de-France” (N°EX061034) and ITMO Cancer of Aviesan and INCa on funds administered by Inserm (N°21CQ016-00) for MS analysis, and from A-PARADDISE/602080 FP7 Project, FISR2019_00374 MeDyCa, and Progetto di Ateneo Sapienza 2021 (no. RM12117A61C811CE) for the preparation of LSD1 inhibitors. This work was also supported by the French National Research Agency (ANR) through the project ELATION (grant number ANR-21-CE15-0046). Maria Gutierrez-Sanchez was supported by a PhD scholarship from the Mexican National Council for Science and Technology” (scholarship number 2019-000004-01EXTF-00037).

## SUPPLEMENTARY FIGURE LEGENDS

**Figure S1. Heat maps of mRNA expression values in BMDM samples.**

BMDMs were left uninfected (UI BMDMs) (A) or infected with lesion-derived mCherry-transgenic amastigotes of *L. amazonensis* (LIMs) (B). One day after infection, MC4030 was added or not to the samples. RNA-Seq was performed one day after this treatment using three independent biological replicates. A Z-score normalization is performed on the normalized read counts across samples for each gene. Z-scores are computed on a gene-by-gene (row-by-row) basis and used to plot heatmap (add the color legend with values).

**Figures S2-5: Gene Set Enrichment Analysis (GSEA) of transcriptionally modulated genes in LIMs.** GSEA analysis of the RNASeq data of UI BMDMs and LIMs. Upregulated (Figure S2) and downregulated (Figure S3) processes are indicated. Sub-panels of the analyses are shown in Figures S4 and S5. Downregulated and upregulated processes are indicated in blue and red colors, respectively.

**Figure S6: Analysis of the impact of LSD1*i* on parasite load in *Leishmania-*infected BMDMs.** BMDMs were infected with mCherry-transgenic amastigotes of *L. amazonensis* isolated from infected mouse footpads (A) or *L. donovani* isolated from infected hamster spleens (B1). After one day of infection, MC2652 and MC4030 were added at a concentration of 10 μM for 3 days. Merged fluorescence and bright field microscopy images of each sample are shown. Macrophage nuclei and parasites were visualized using Hoechst 33342 staining (blue fluorescence) and mCherry expression (red fluorescence), respectively. Green fluorescence corresponds to dead macrophages as revealed by YOPRO-1 incorporation. (B2) Violin plots showing the level the mCherry fluorescence as a readout for *L. donovani* burden. Grey, BMDMs treated with DMSO treated; violet, MC2652-treated, green; MC4030-treated. Raw Integrated Density per cell was determined by ImageJ. The p-value is indicated (*: p<0.0001).

**Figure S7: Effect of a transfection with *Lsd1*/*Kdm1a* siRNAs performed before parasite infection.**

Experimental workflow (A) and analysis of uninfected BMDMs (B) and LIMs (C) in response to transfection with two siRNAs (si1 and si3 versus a control siRNA, siCtrl). Multiparametric analyses included RTqPCR of *Lsd1/Kdm1a* transcripts and negative control transcripts (*Rcor1, Cat* and *Ptgs1*) (B1 and C1), Western blot analysis and relative quantitation of LSD1 protein (B2 and C2), and analysis on cell morphology (B3 and C3) and parasite load by epifluorescnce microscopy analysis of parasite -related mCherry fluorescence (C3).

**Figure S8: Effect of a transfection with *Lsd1/Kdm1a* siRNAs performed after parasite infection**

Experimental workflow (A) and analysis of uninfected BMDMS (B) and LIMs (C) in response to transfection with two siRNAS (si1 and si3 versus a control siRNA siCtrl). Multiparametric analyses included RTqPCR of *Kdm1a* transcripts and negative control transcripts (*Rcor1, Cat* and *Ptgs1*) (B1 and C1), Western blot analysis and relative quantitation of LSD1 protein (B2 and C2) and analysis on cell morphology (B3 and C3) and parasite load (epifluorescnce microscopy analysis of parasite -related mCherry fluorescence (C3).

**Figure S9: Reaction conditions for the synthesis of HyTs.**

a) HATU, *N, N-*diisopropylethylamine, dimethylformamid/dichloromethane (1:1), rt, 16 h, 81%; b) NaN_3_, acetonitrile, 80 °C, 16 h, 40%; c) Trimethylsulfoxonium iodide, NaH, DMSO, rt, 48 h, 93%; d) KOH, MeOH, rt, 16 h, 100%; e) i) diphenylphosphoryl azide, *tert*-butanol, 60 °C, 3 h, ii) Boc_2_O, 80 °C, 16 h, 76%; f) 4-(bromomethyl)-1,2-dichlorobenzene, NaH, dimethylformamid, 0 °C to rt, 4 h, 95%; g) CuI, (Pd(PPh[)[), Et_3_N, dioxane, 80°C, 16 h, 81%; h) K_2_CO_3_, MeOH/ dichloromethane (1:1), rt, 5 h, 80%; i) intermediate 2, CuSO_4_, NaAscorbat, tris((1-benzyl-4-triazolyl)methyl)amine, water/acetonitrile (1:1), rt, 16 h; j) TFA, DCM, rt, 2h, 40% over two steps.

**Figure S10: Structure and biological effect of Hydrophobic Tags (HyTs) on *L.am* promastigotes in culture or amastigotes in LIMs.**

High Content Assay (HCA) and Dye Reduction Assay (DRA) results at 2.5 µM, 5 µM, 10 µM or 20 µM of the indicated compound. The level of host cell toxicity or anti-leishmanial activity is indicated by the red intensity: red, > 70%; light red ,40% - 70%, grey < 40% (considered negligible). All data data was normalized to the DMSO control. Mø stands for uninfected macrophages, PV for parasitophorous vacuoles and Am for amastigotes. Mø, PV and Am were evaluated by HCA assay. Free, lesion-derived amastigotes and cultured promastigotes were measured by DRA assay. (A) Depiction of the data for HyTs containing adamantane as hydrophobic group as dichlorobenzyl derivatives. AAB-362 serves as a negative control. (B) HyTs with flourene and heptafluorobutyl groups are depicted. They are linked either to a dichlorobenzyl derivative as warhead or a benzensulfonamide derivative. AAB-020 is the parent compound without a hydrophobic group. The grey zone corresponds to the hydrophobic tags and green zone to the warhead.

**Figure S11: LSD1 cellular thermal shift asssay for compounds NT24 (A) and AABO20**

**(B) in THP-1 cells.** The melting curves for LSD1 in the presence of DMSO (control) is shown in black. Compounds at 30 µM were incubated for 2 hours in two independent experiments. The difference in protein stability between the DMSO and compound-treated samples is represented as ΔTm.

**Figure S12: Cellular thermal shift assay of LSD1, MAO-A, and GAPDH in protein lysates from uninfected (UI) BMDM and in LIMs in presence of the LSD1 inhibitors.**

UI BMDMs (A) and LIMs at day 1 post-infection (B) were treated for 24 hours with 1% DMSO as control or with 15 µM MC2652 or MC4030. Extracts were resolved by SDS-PAGE and proteins of interest revealed by Western blot analysis using specific antibodies. One representative Western blot out of 2-3 independent experiments performed is shown.

**Figure S13: Quality control of nuclear and cytoplasmic extracts isolated from uninfected (UI) BMDMs and LIMs.**

**(A)** Microscopic images of live BMDMs (upper panel) and isolated nuclei (lower panel).

**(B)** Western blot analysis of GAPDH in nuclear and cytoplasmic extracts of 5 biological replicates. The absence of GAPDH indicates the absence of cytoplasmic contamination in nuclear extracts.

## SUPPLEMENTARY INFORMATION ON CHEMISTRY

**(Hydrophobic Tags synthesis).**

For synthesis of Hydrophobic Tags, all reactions were carried out in glassware under inert nitrogen atmosphere. The chemicals and reagents used were purchased by BLDPharm, FischerScientific, SigmaAldrich, Arcros organics and MedChemExpress were used without further purifications. For the reaction monitoring, thinlayer chromatography (TLC) was performed with Merck alumina plates coated with silica gel 60 F254 and silica gel 60 RP-18 F254s and visualized under UV light (254 nm and 365 nm). Staining agents such as KMNO4, Bromocresol green, ninhydrine and triphenylphosphine were used. The mobile phases were adjusted to the corresponding compounds properties. The purification was performed as flash column chromatography with Biotage® Isolera Prime/One purification instrument using 40-60 µm pre-packed silica gel columns from Biotage®, Sfär Silica D60 µm, Sfär KP amino D 50 µM or Sfär Silica HC D 20 µM. For the product identification of the compounds, NMR spectroscopy and mass spectroscopy were used. NMR spectra were acquired on a BRUKER Avance 400 spectrometer (400 MHz and 100.6 MHz for 1H, 19F and 13C) and BRUKER 700 spectrometer (700 MHz for HMBC, HSQC and NOESY) at a temperature of 303 K using DMSO-d6, methanol-d4 and acetonitrile-d3 solvents. Chemical shifts (0) are reported in ppm, multiplicity is designated as broad singlet (bs), singlet (s), doublet (d), doublet of doublets (dd), doublet of doublet of doublets (ddd), doublet of triplets (dt), triplet (t), triplet of doublets (td), quartet (q), multiplet (m), coupling constant (J) are expressed in Hz. The 1H assignment resulted from COSY experiments. Mass spectra were recorded on an Advion expression CMS using an ASAP® (Atmospheric Solids Analysis Probe; a.k.a APCI: Atmospheric Pressure Chemical Ionization) as ion source, on a Thermo Scientific Exactive mass spectrometer using electrospray ionization (ESI) as ion source or HR-MS were obtained on a THERMO SCIENTIFIC Advantage. HPLC analysis was performed to determine the purity of all final compounds on an Agilent Technologies 1260 Infinity II system using diode array detector (DAD) UV detection at 210, 230, 248, 254, 260 and 280 nm. The following parameters were used: XBridge ®Shield RP18 5 µm, 130 Å, 4.6 x 150 mm column from waters, eluent A was H2O containing 0.05% trifluoroacetic acid (TFA) and eluent B was acetonitrile. Gradient: 0-4 min: 90:10 (A/B); 4-19 min: 90:10 → 0:100 (A/B); 19-21 min: 0:100 (A/B); 21-21.5 min: 0:100 → 90:10 (A/B); 21.5-25 min: 90:10 (A/B) with a flowrate of 1 mL/min. For the final purification preparative HPLC was performed for all final compounds on an Agilent 1260 Infinity II at 210 nm using the same parameters. All compounds were solubilized in DMSO at a 10 mM concentration and stored as aliquots at -20°C in 1.5 mL Eppendorf tubes. Compounds targeting LSD1 (LSD1*i*) were used at different concentrations depending on the assay as indicated in Results.

Method A: XBridge ®Shield RP18 5 µm, 130 Å, 4.6 x 150 mm column from waters, eluent A was H2O containing 0.05% trifluoroacetic acid (TFA) and eluent B was acetonitrile. Gradient: 0-4 min: 90:10 (A/B); 4-19 min: 90:10 → 0:100 (A/B); 19-21 min: 0:100 (A/B); 21-21.5 min: 0:100 → 90:10 (A/B); 21.5-25 min: 90:10 (A/B) with a flowrate of 1 mL/min.

Method B: XBridge® Prep Shield RP18 5 µm OBD™, 130 Å, 19 x 150 mm column was used, eluent A was H2O containing 0.05% trifluoracetic acid (TFA) and eluent B was acetonitrile. In general the gradient was: 0- 4 min: 90:10 (A/B); 4-19 min: 90:10 → 0:100 (A/B); 19-21 min: 0:100 (A/B); 21-21.5 min: 0:100 → 90:10 (A/B); 21.5-25 min: 90:10 (A/B) with a flowrate of 17.1 mL/min. For each final compound the gradient conditions were optimized and are listed in the final compound description.

N-(((3r,5r,7r)-adamantan-1-yl)methyl)-5-bromopentanamide--methane (1)

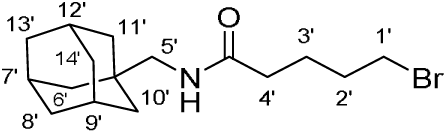

Compound (1) was synthesized from 5-bromovaleric acid (150 mg, 0.82 mmol), 1-adamantanmethylamin (137 mg, 0.82 mmol), HATU (473 mg, 1.23 mmol, 1.5 eq.) and DIPEA (423 µL, 2.46 mmol, 3.00 eq.) in DCM/DMF (1:1, 8.2 mL, 0.1 M). The reaction mixture stirred for 16 h at 23°C (room temperature). After extraction with DCM and evaporating to dryness, the crude product was purified by flash chromatography (n-heptane/EtOAc, 0-40%) to obtain the product as colourless oil (200 mg, 74%.).1H NMR (400 MHz, DMSO-d6) δ 7.65 (t, J = 6.3 Hz, 1H, CONH-), 3.53 (t, J = 6.7 Hz, 2H, H1’), 2.75 (d, J = 6.3 Hz, 2H, H5’), 2.13 (t, J = 7.3 Hz, 2H, H4’), 2.01 – 1.87 (m, 3H, H7’, H9’, H12’), 1.71 – 1.53 (m, 10H, H8’, H13’, H14’, H2’, H3’), 1.41 (d, J = 2.8 Hz, 6H, H6’ H10’, H11’). APCI calc. for C16H27BrNO [M+H]+: 328.12, found: 328.6/330.6 [M+H]+. N-(((3r,5r,7r)-adamantan-1-yl)methyl)-5-azidopentanamide--methane (2)

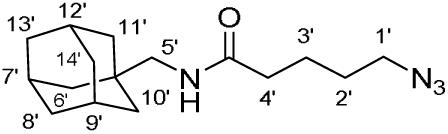

To a solution of compound 1 (200 mg, 0.60 mmol, 1.00 eq.) in ACN (0.8 mL, 0.8 M), sodium azide (119 mg, 1.80 mmol, 3.00 eq.) was added. The reaction was heated to 80 °C and stirred for 16 h. After it was left to cool down to room temperature, the white suspension was diluted with water and extracted with EtOAC (3 x 50 mL). The combined organic layers were dried with sodium sulfate, filtrated and concentrated under reduced pressure. The crude product was purified by flash chromatography (n-heptane/EtOAc 0-40 %) to receive the product as colourless oil (90 mg, 40 %).1H NMR (400 MHz, DMSO-d6) δ 7.66 (t, J = 6.4 Hz, 1H, - CONH-), 3.35 (s, 2H, H1’ under water peak), 2.77 (d, J = 6.2 Hz, 2H, H5’), 2.20 – 2.05 (m, 2H, H4’), 1.92 (p, J = 3.1 Hz, 3H, H7’, H9’, H12’), 1.71 – 1.63 (m, 4H, H8’a, H13’a, H14’a), 1.63 – 1.37 (m, 7H, H8’b, H13’b, H14’b, H2’, H3’), 1.42 (d, J = 2.8 Hz, 6H, H6’, H10’, H11’). APCI calc. for C16H27N4O [M+H]+: 291.21, found: 291.3 [M+H]+. Ethyl 2-(4-bromophenyl)cyclopropane-1-carboxylate rac. (3)

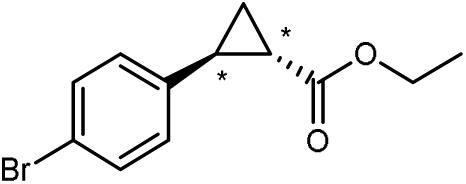

In a 250 mL round bottom flask, sodium hydride 60 % in mineral oil (392 mg, 9.80 mmol, 0.1 eq.) was suspended in anhydrous DMSO (39.2 mL, 0.25 M). Trimethylsulfoxonium iodide (2.40 g, 10.78 mmol, 1.1 eq.) was added portions wise and the solution stirred for 30 min at rt. A solution of ethyl (E)-3-(4-bromophenyl)acrylate (2.50 g, 9.80 mmol, 1.00 eq.) in DMSO (71.9 mL) was then transferred dropwise over 20 minutes. The reaction flask was left to stir at 23°C (room temperature, rt) for 16 hours. When the reaction was completed, the clear solution had turned pale yellow. The product was extracted 3 times using 150 mL of ethyl acetate (EtOAc), the combined organic layers were washed using a saturated aqueous solution of saturated NaCl (brine) and dried over anhydrous sodium sulfate. The product was filtered, dried in vacuo and purified using column chromatography (n-heptane/EtOAc; 0-50%). After drying, a clear oil was obtained. Yield: 71-93 %. 1H NMR (400 MHz, DMSO-d6) δ 7.49 – 7.40 (m, 2H, m-Ar), 7.19 – 7.11 (m, 2H, o-Ar), 4.09 (qd, J = 7.1, 0.7 Hz, 2H, CH2), 2.43 (ddd, J = 9.1, 6.5, 4.1 Hz, 1H, H2), 1.94 (ddd, J = 8.4, 5.3, 4.2 Hz, 1H, H1), 1.46 (ddd, J = 9.2, 5.3, 4.5 Hz, 1H, H3a), 1.37 (ddd, J = 8.4, 6.5, 4.5 Hz, 1H, H3b), 1.20 (t, J = 7.1 Hz, 3H, CH3). APCI calc. for C12H14BrO2 [M+H]+: 269.01, found: 268.8/270.8 [M+H]+. (1S*,2R*)-2-(4-Bromophenyl)cyclopropane-1-carboxylic acid rac. (4)

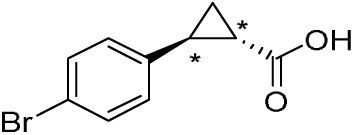

To a solution of compound 3 (2.21 g, 8.21 mmol, 1.000 eq.) in EtOH (32.8 mL, 0.25 M) an aqueous KOH solution (1.0 M, 16.42 mmol, 2.00 eq.) was added. The reaction mixture was stirred for 16 h at room temperature and then concentrated in vacuo. The mixture was acidified with an aqueous solution of HCl (1.00 M) to pH 1-2, as indicated by pH paper resulting in the formation of a white precipitate. The product was extracted with EtOAc (25 mL x 4). The combined organic layers were dried over anhydrous sodium sulfate, filtered, and concentrated in vacuo again. The product was a white solid which was used for the next reaction step without further purification. Yield: 95%. 1H NMR (400 MHz, DMSO-d6) δ 12.32 (s, 1H, COOH), 7.49 – 7.40 (m, 2H, m-Ar), 7.18 – 7.11 (m, 2H, o-Ar), 2.39 (ddd, J = 9.1, 6.4, 4.1 Hz, 1H, H2), 1.81 (ddd, J = 8.4, 5.3, 4.1 Hz, 1H, H1), 1.45 – 1.39 (m, 1H, H3), 1.32 (ddd, J = 8.4, 6.4, 4.4 Hz, 1H, H3b). APCI: calc. for C10H10BrO2 [M+H]+: 240.98, found 240.9/242.9 [M+H]+. tert-Butyl ((1S*,2R*)-2-(4-bromophenyl)cyclopropyl)carbamate rac. (5)

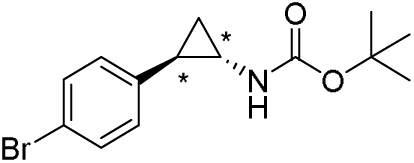

Under nitrogen atmosphere in a dry heated flask was charged with compound 4 (2.00 g, 8.30 mmol, 1.00 eq.) in tert-butanol (30 mL) and triethylamine (2.3 mL, 16.6 mmol, 2.00 eq.). Afterwards Diphenylphosphoryl azide (DPPA) (1.96 mL, 9.12 mmol, 1.10 eq.) and molecular sieves were added. The reaction mixture was stirred at 80 °C for 24 h. Then the mixture was allowed to cool down to room temperature and tert-butanol was removed in vacuo. The resulting brown slurry suspension was filtered and washed with EtOAc (20 mL x 3). The combined organic layers were dried over anhydrous sodium sulfate, filtered, and concentrated in vacuo again. The crude product was purified by silica gel column chromatography (cyclohexane/EtOAc; 0-50%), to afford the brown solid 5 (76 %). 1H NMR (400 MHz, DMSO-d6) δ 7.45 – 7.38 (m, 2H, m-Ar), 7.09 – 7.02 (m, 2H, o-Ar), 2.59 (d, J = 6.9 Hz, 1H, H1), 1.88 (ddd, J = 9.4, 6.3, 3.2 Hz, 1H, H2), 1.37 (s, 9H, C(CH3)3), 1.12 (ddd, J = 9.6, 5.5, 4.6 Hz, 1H, H3a), 1.06 (q, J = 6.5 Hz, 1H, H3b). APCI: calc. for C14H19BrNO2 [M+H]+: 312.05, found: 255.9/257.9 [M(-tert-butyl) +H]+, 212.0/213.9 [M(-Boc)+H]+. tert-Butyl ((1S*,2R*)-2-(4-bromophenyl)cyclopropyl)(3,4-dichlorobenzyl)carbamate rac. (6)

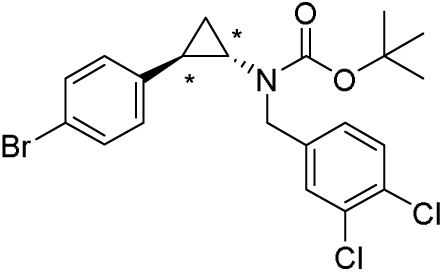

Compound 6 was obtained starting from compound 5 (300 mg, 0.96 mmol, 1.00 eq.) dissolved in DMF (0.25 M). Sodium hydride (60% in mineral oil, 1.1 eq.) was added portionwise at 0 °C and 4-(bromomethyl)-1,2-dichlorobenzen was then added to the reaction mixture. Purification after flash column chromatography on silica (n- heptane/EtOAc; 0-50%) gave a colourless oil. Yield: 95%. 1H NMR (400 MHz, DMSO-d6) δ 7.60 (d, J = 8.3 Hz, 1H, H5-benzyl), 7.44 (d, J = 2.0 Hz, 1H, H2-benzyl), 7.42 – 7.37 (m, 2H, m-Ar), 7.20 (dd, J = 8.3, 2.1 Hz, 1H, H6-benzyl), 7.07 (d, J = 8.4 Hz, 2H, o-Ar), 4.51 (d, J = 15.9 Hz, 1H, NCH), 4.32 (d, J = 15.9 Hz, 1H, NCH’), 2.62 (ddd, J = 7.7, 4.6, 3.3 Hz, 1H, H1), 2.15 (ddd, J = 9.8, 6.6, 3.3 Hz, 1H, H2), 1.33 (s, 10H, H3, C(CH3)3 ), 1.17 (td, J = 7.1, 1.5 Hz, 1H, H3b). APCI: calc. for C21H23BrCl2NO2 [M+H]+: 470.02, found: 413.8/415.8/417.8/419.8 [M(-tert-butyl) +H]+, 369.8/371.8/373.8/375.8 [M(-Boc)+H]+. tert-Butyl (3,4-dichlorobenzyl)((1S*,2R*)-2(4((trimethylsilyl)ethynyl)phenyl)cyclopropyl) carbamate rac. (7)

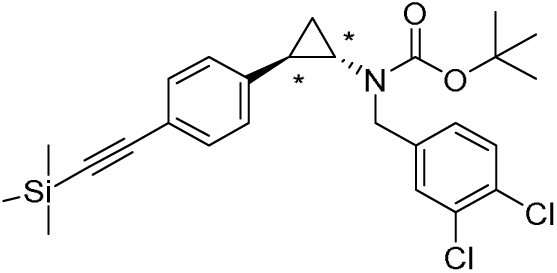

Compound 6 (350 mg, 0.74 mmol, 1.00 eq.) was dissolved with copper(I) iodide (14 mg, 0.07 mmol, 0.10 eq.) and tetrakis(triphenylphosphin)palladium(0) (86 mg, 0.07 mmol. 0.10 eq.) in dry dioxane/trimethylamine (7.4 mL, 0.1 M, 1:1). The reaction mixture stirred at 80 °C for 16 h. After extraction with ethyl acetate, the product was isolated as a brown-orange oil after flash chromatography (n-heptane/EtOAc; 0-80%). Yield: 293 mg; 81%. 1H NMR (400 MHz, DMSO-d6) δ 7.62 (t, J = 8.5 Hz, 1H, H5-benzyl), 7.45 (d, J = 2.0 Hz, 1H, H2-benzyl), 7.34 – 7.28 (m, 2H, m-Ar), 7.20 (dd, J = 8.3, 2.1 Hz, 1H, H6-benzyl), 7.09 (d, J = 8.0 Hz, 2H, o-Ar), 4.53 (d, J = 17.9 Hz, 1H NCH), 4.31 (d, J = 16.0 Hz, 1H, NCH’), 2.69 – 2.62 (m, 1H, H1), 2.17 (ddd, J = 9.7, 6.5, 3.3 Hz, 1H, H2), 1.33 (s, 1H, H3a), 1.32 (s, 9H, C(CH3)3), 1.22 (d, J = 8.3 Hz, 1H, H3b), 0.21 (s, 9H, Si(CH3)3). APCI: calc. C26H32Cl2NO2Si [M+H]+: 488.15, found: 432.1/434.1/436.0 [M(-tert-butyl) +H]+, 388.0/390.0/392.0 [M(-Boc)+H]+. tert-Butyl (3,4-dichlorobenzyl)((1S*,2R*)-2-(4-ethynylphenyl)cyclopropyl)carbamate rac. (8)

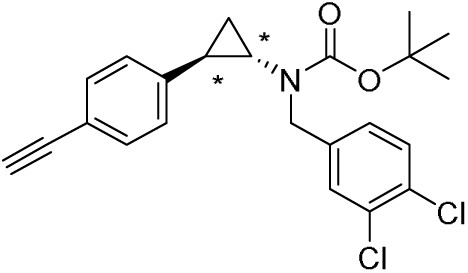

Compound 7 (250 mg, 0.51 mmol, 1.00 eq.) was stirred in MeOH (0.50 M, 1.00 mL) and K2CO3 (317 mg, 1.02 mmol, 2.00 eq.) at rt for 16 h. After completion, the reaction was extracted with ethyl acetate and the solvent was evaporated in vacuo. The product was isolated after the purification (n-heptane / EtOAc; 0-80%), as a brown-orange oil. Yield: 170 mg, 60-80%. 1H NMR (400 MHz, DMSO-d6) δ 7.61 (d, J = 8.2 Hz, 1H, H5-benzyl), 7.44 (d, J = 2.1 Hz, 1H, H2-benzyl), 7.37 – 7.30 (m, 2H, m-Ar), 7.20 (dd, J = 8.3, 2.1 Hz, 1H, H6- benzyl), 7.10 (d, J = 8.2 Hz, 2H, o-Ar), 4.52 (d, J = 15.8 Hz, 1H, NCH), 4.32 (d, J = 15.9 Hz, 1H, NCH), 4.11 (s, 1H, CCH), 2.65 (dt, J = 7.9, 3.0 Hz, 1H, H1), 2.23 – 2.10 (m, 1H, H2), 1.35 (m, 1H, H1), 1.33 (d, J = 2.7 Hz, 9H, C(CH3)3), 1.26 – 1.21 (m, 1H, H1’). APCI: calc. for C23H24Cl2NO2 [M+H]+: 416.11, found: 360.1/362.1/364.1 [M(-tert-butyl) +H]+, 316.1/318.1/320.1 [M(-Boc)+H]+. N-(((3r,5r,7r)-adamantan-1-yl)methyl)-5-(4-(4-((1R*,2S*)-2-((3,4-dichlorobenzyl)amino)cyclopropyl)phenyl)-1H-1,2,3-triazol-1-yl)pentanamide—methane rac. (10) (NT24)

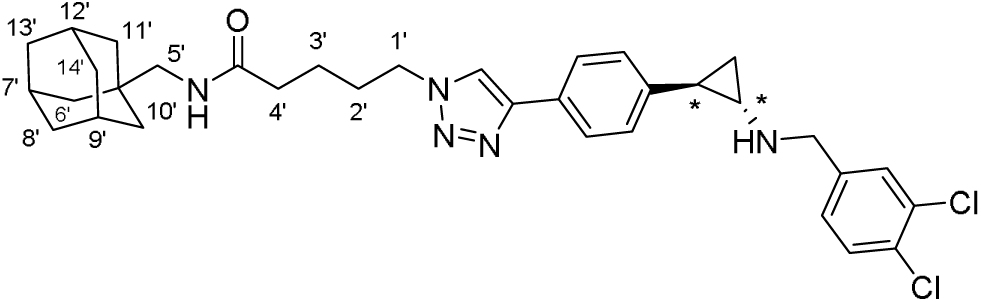

In a 10 mL-25 mL flask, the alkyne derivative (8) (25.0 mg, 0.06 mmol, 1.00 eq.) and the corresponding azide (2) (17.0 mg, 0.06 mmol, 1.00 eq.) were suspended in a 1:1 mixture of demin. water and acetonitrile (0.05 M). Afterwards first a 1 M aqueous copper(II) sulfate solution (0.20 eq.) was added, directly followed by sodium ascorbate (0.40 eq.). (Tris(benzyltriazolylmethyl)amine (TBTA, 0.10 eq.) was dissolved in DMF (0.02 M) and then added. The reaction mixture was stirred for 2-16 h at room temperature until complete conversion of the reaction. The reaction was monitored by analytical HPLC. After extraction with dichloromethane and water, the organic layers were concentrated under reduced pressure. The crude product was either directly used in the next reaction. The subsequent deprotection was carried out with DCM/TFA (1:1; 1.0 eq.) and without further purification, the reaction mixture was immediately purified by preparative HPLC method B. 1H NMR (400 MHz, DMSO-d6) δ 9.40 (d, J = 37.0 Hz, 2H, NH2¬+), 8.55 (s, 1H, H5 triazole), 7.80 (d, J = 2.0 Hz, 1H, H5 benzyl), 7.80 – 7.72 (m, 2H, o-Ar triazole), 7.72 (d, J = 8.2 Hz, 1H, H2 benzyl), 7.65 (t, J = 6.3 Hz, 1H, CONH), 7.48 (dd, J = 8.3, 2.1 Hz, 1H, H6 benzyl), 7.24 – 7.14 (m, 2H, m-Ar triazole), 4.44 – 4.30 (m, 4H, NCH2, H1’), 3.00 (m, 1H, H1), 2.74 (d, J = 6.3 Hz, 2H, H5’), 2.39 (ddd, J = 10.2, 6.6, 3.5 Hz, 1H, H2), 2.15 (t, J = 7.3 Hz, 2H, H4’), 1.88 (s, 5H, H2’, H9’, H7’, H12’), 1.89 –1.62 (d, J = 12.0 Hz, 6H, H8’, H13’, H14’), 1.54 (d, J = 11.1 Hz, 2H, H3), 1.53 – 1.41 (m, 7H), 1.39 (d, J = 2.7 Hz, 7H, H, H6’, H11’, H10’). HRMS calc. for C35H46Cl2N5O [M+H]+: 606.27, found: 606.2761/608.2734/610.2700 [M+H]+. HPLC (method A): tR = 12.991 min; UV purity at 210 nm: 96%. (1S*,2R*)-2-(4-(1-(2-(2-(((3s,5s,7s)-adamantan-1-yl)oxy)ethoxy)ethyl)-1H-1,2,3-triazol-4-yl)phenyl)-N-(3,4-dichlorobenzyl)cyclopropan-1-amine rac. NT29

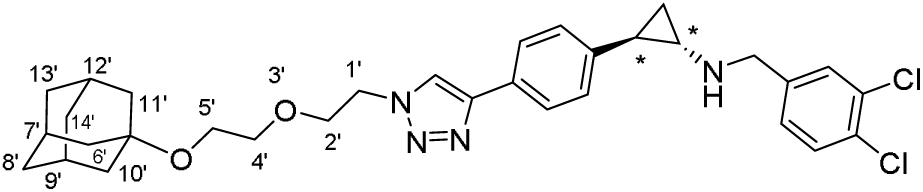

In a 10 mL-25 mL flask, the alkyne derivative (8) (25.0 mg, 0.06 mmol, 1.00 eq.) and the corresponding azide (15.0 mg, 0.06 mmol, 1.00 eq.) were suspended in a 1:1 mixture of demin. water and acetonitrile (0.05 M). Afterwards first a 1 M aqueous copper(II) sulfate solution (0.20 eq.) was added, directly followed by sodium ascorbate (0.40 eq.). (Tris(benzyltriazolylmethyl)amine (TBTA, 0.10 eq.) was dissolved in DMF (0.02 M) and then added. The reaction mixture was stirred for 2-16 h at room temperature until complete conversion of the reaction. The reaction was monitored by analytical HPLC. After extraction with dichloromethane and water, the organic layers were concentrated under reduced pressure. The crude product was either directly used in the next reaction. The subsequent deprotection was carried out with DCM/TFA (1:1; 1.0 eq.) and without further purification, the reaction mixture was immediately purified by preparative HPLC method B. 1H NMR (400 MHz, DMSO-d6) δ 9.38 (s, 2H, -NH2+-), 8.49 (s, 1H, H5 triazole), 7.80 (d, J = 2.0 Hz, 1H, H2 benzyl), 7.78 – 7.74 (m, 2H, o-Ar triazole), 7.72 (d, J = 8.3 Hz, 1H, H5 benzyl), 7.48 (dd, J = 8.3, 2.1 Hz, 1H, H6 benzyl), 7.23 – 7.13 (m, 2H, m-Ar triazole), 4.55 (t, J = 5.1 Hz, 2H, H1’), 4.36 (s, 2H, -NCH2-), 3.90 – 3.80 (m, 2H, H2’), 3.48 (td, J = 4.7, 1.3 Hz, 2H, H4’), 3.42 (td, J = 4.7, 1.4 Hz, 2H, H5’), 2.98 (s, 1H, H1), 2.39 (ddd, J = 10.1, 6.5, 3.5 Hz, 1H, H2), 2.02 (s, 3H, H7’, H9’, H12’), 1.59 (d, J = 2.9 Hz, 6H, H6’, H10’, H11’), 1.54 (d, J = 12.8 Hz, 3H, H8’a, H13’a, H14’a), 1.51 – 1.41 (m, 4H, H3a H8’b, H13’b, H14’b), 1.35 (q, J = 6.8 Hz, 1H, H3b). HRMS calc. for C32H39Cl2N4O2 [M+H]+: 581.24, found: 581.2454 [M+H]+. HPLC (method B): tR = 12.748 min; UV purity at 210 nm: 95%.

2-((3r,5r,7r)-adamantan-1-yl)-N-((4’-((1R*,2S*)-2-((3,4-dichlorobenzyl)amino)cyclopropyl)-[1,1’-biphenyl]-3-yl)methyl)acetamide rac. AAB175

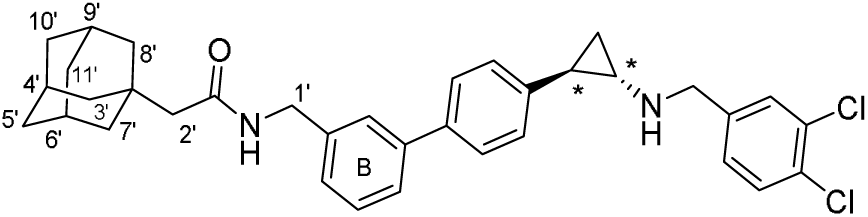

1H NMR (400 MHz, Methanol-d4) δ 7.68 (d, J = 2.1 Hz, 1H, H2 benzyl), 7.61 (d, J = 8.2 Hz, 1H, H6 benzyl), 7.56 (dd, J = 8.3, 1.9 Hz, 3H, o-Ar B, H2 B), 7.48 (ddd, J = 7.7, 1.9, 1.2 Hz, 1H, H4 B), 7.45 – 7.34 (m, 2H, H5 B, H6 benzyl), 7.28 (ddd, J = 7.5, 1.8, 1.1 Hz, 1H, H6 B), 7.17 – 7.12 (m, 2H, m-Ar B), 4.45 – 4.34 (m, 4H, NCH2, H1’), 3.01 (ddd, J = 8.0, 4.5, 3.6 Hz, 1H, H1), 2.40 (ddd, J = 10.4, 6.7, 3.6 Hz, 1H, H2), 2.00 (s, 2H, H2’), 1.94 (d, J = 14.5 Hz, 3H, H4’, H6’, H9’), 1.77 – 1.67 (m, 6H, H5’, H10’, H11’), 1.65 (d, J = 2.9 Hz, 11H, H3’, H7’, H8’), 1.52 (ddd, J = 10.3, 6.9, 4.5 Hz, 1H, H3a), 1.44 (dt, J = 7.8, 6.8 Hz, 1H, H3b). HRMS calc. for C35H39Cl2N2O [M+H]+: 573.24, found: 573.2443 [M+H]+. HPLC (method B): tR = 13.974 min; UV purity at 210 nm: 96%.

7-(2-((3r,5r,7r)-adamantan-1-yl)acetamido)-N-((4’-((1R*,2S)-2-((3,4-dichlorobenzyl)amino)cyclopropyl)-[1,1’-biphenyl]-3-yl)methyl)heptanamide rac. (AAB167)

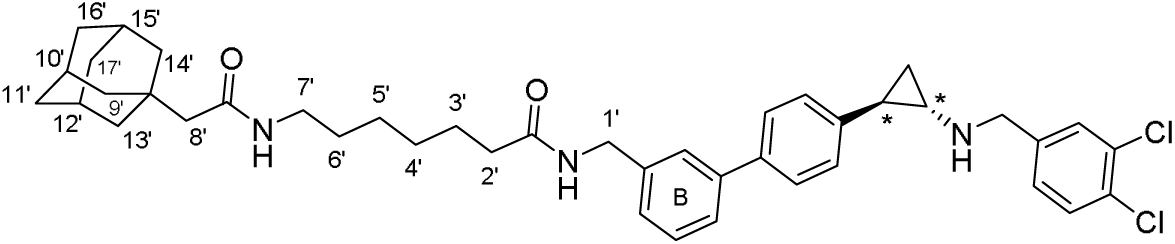

1H NMR (400 MHz, Methanol-d4) δ 7.69 (d, J = 2.1 Hz, 1H, H2 benzyl), 7.62 (d, J = 8.3 Hz, 1H, H6 benzyl), 7.58 – 7.53 (m, 2H, o-Ar B), 7.52 (dt, J = 1.9, 1.0 Hz, 1H, H2 B), 7.48 (dt, J = 7.8, 1.5 Hz, 1H, H4 B), 7.43 (dd, J = 8.3, 2.1 Hz, 1H, H5 benzyl), 7.39 (t, J = 7.6 Hz, 1H, H5 B), 7.26 (dt, J = 7.7, 1.3 Hz, 1H, H6 B), 7.20 – 7.14 (m, 2H, m-Ar B), 4.42 (d, J = 5.6 Hz, 4H, NCH2, H1’), 3.10 (t, J = 6.8 Hz, 2H, H7’), 3.07 – 2.99 (m, 1H, H1), 2.41 (ddd, J = 10.4, 6.7, 3.6 Hz, 1H, H2), 1.93 (d, J = 3.9 Hz, 5H, H2’, H10’, H12’, H15’), 1.90 (s, 2H, H8’), 1.73 (d, J = 12.3 Hz, 3H, H11’a, H16’a, H17’a), 1.69 – 1.58 (m, 11H, H3’, H9’, H11’b, H13’, H14’, H16’b, H17’b), 1.53 (ddd, J = 10.3, 6.9, 4.5 Hz, 1H, H3a), 1.46 (dt, J = 7.7, 6.8 Hz, 1H, H3b), 1.39 – 1.26 (m, 6H, H4’-H6’). HRMS calc. for C42H52Cl2N3O2 [M+H]+: 700.34, found: 700.3426 [M+H]+. HPLC (method B): tR = 13.856 min; UV purity at 210 nm: 95%.

N-(4-(4-(4-((1R*,2S*)-2-((3,4-dichlorobenzyl)amino)cyclopropyl)phenyl)-1H-1,2,3-triazol-1-yl)butyl)-2-(9H-fluoren-9-yl)acetamide rac. (PB17)

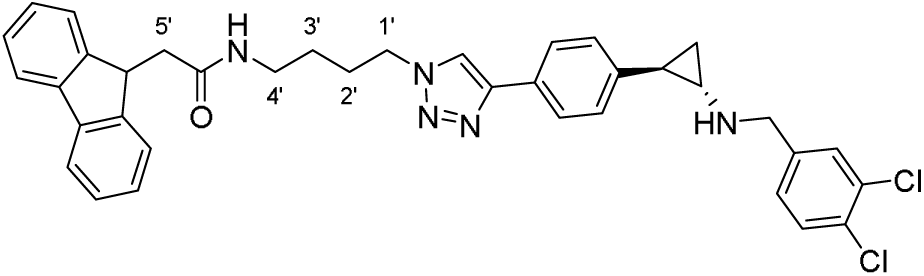

In a 10 mL-25 mL flask, the alkyne derivative (8) (20.0 mg, 0.04 mmol, 1.00 eq.) and the corresponding azide (20.0 mg, 0.04 mmol, 1.00 eq.) were suspended in a 1:1 mixture of demin. water and acetonitrile (0.05 M). Afterwards first a 1 M aqueous copper(II) sulfate solution (0.20 eq.) was added, directly followed by sodium ascorbate (0.40 eq.). (Tris(benzyltriazolylmethyl)amine (TBTA, 0.10 eq.) was dissolved in DMF (0.02 M) and then added. The reaction mixture was stirred for 2-16 h at room temperature until complete conversion of the reaction. The reaction was monitored by analytical HPLC. After extraction with dichloromethane and water, the organic layers were concentrated under reduced pressure. The crude product was either directly used in the next reaction. The subsequent deprotection was carried out with DCM/TFA (1:1; 1.0 eq.) and without further purification, the reaction mixture was immediately purified by preparative HPLC method B. 1H NMR (400 MHz, DMSO-d6) δ 9.40 (d, J = 49.8 Hz, 2H, -NH2+-), 8.59 (s, 1H, H5 triazole), 7.99 (t, J = 5.6 Hz, 1H, -CONH-), 7.85 (dt, J = 7.6, 0.9 Hz, 2H, H4 and H5 fluoren), 7.82 – 7.75 (m, 3H, H2 benzyl, o-Ar triazole), 7.75 – 7.70 (m, 1H, H5 benzyl), 7.53 – 7.44 (m, 3H, H6 benzyl, H1 and H8 fluoren), 7.35 (tt, J = 7.4, 0.9 Hz, 2H, H3 and H6 fluoren), 7.26 (td, J = 7.4, 1.2 Hz, 2H, H2 and H7 fluoren), 7.23 – 7.15 (m, 2H, m-Ar triazole), 4.44 (t, J = 6.9 Hz, 2H, H1’), 4.40 – 4.30 (m, 3H, NCH2, H9 fluoren), 3.26 – 3.18 (m, 2H, H4’), 3.00 (d, J = 4.0 Hz, 1H, H1), 2.48 (s, 2H, H5’ under DMSO peak), 2.39 (ddd, J = 10.1, 6.5, 3.4 Hz, 1H, H2), 1.98 – 1.81 (m, 3H, H2’), 1.55 – 1.40 (m, 3H, H3a, H3’), 1.36 (dt, J = 7.8, 6.5 Hz, 1H, H3b). HRMS calc. for C37H36Cl2N5O [M+H]+: 366.22, found: 363.2304 [M+H]+. HPLC (method A): tR = 14.875 min; UV purity at 210 nm: 100%.

(1S*,2R*)-N-(3,4-dichlorobenzyl)-2-(3’-((2,2,3,3,4,4,4-heptafluorobutoxy)methyl)-[1,1’-biphenyl]-4-yl)cyclopropan-1-amine rac. (AAB243)

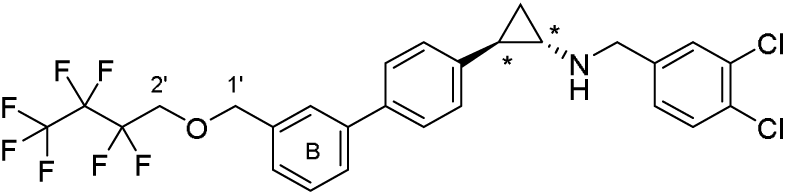

1H NMR (400 MHz, Methanol-d4) δ 7.69 (d, J = 2.1 Hz, 1H, H2 benzyl), 7.62 (d, J = 8.3 Hz, 1H, H5 benzyl), 7.60 – 7.53 (m, 3H, o-Ar B, H2 B), 7.49 – 7.41 (m, 2H, H4-5 B), 7.34 (dt, J = 7.8, 1.3 Hz, 1H, H6 B), 7.20 – 7.15 (m, 2H, m-Ar B), 4.74 (s, 2H, H1’), 4.46 – 4.35 (m, 2H, - NCH2-), 4.16 – 4.04 (m, 2H, H2’), 3.03 (ddd, J = 8.0, 4.5, 3.6 Hz, 1H, H1), 2.40 (ddd, J = 10.4, 6.7, 3.7 Hz, 1H, H2), 1.52 (ddd, J = 10.2, 6.9, 4.5 Hz, 1H, H3a), 1.46 (dt, J = 7.8, 6.8 Hz, 1H, H3b). HRMS calc. for C27H22Cl2F7NO [M+H]+: 580.10, found: 508.1036 [M+H]+. HPLC (method B): tR = 14.399 min; UV purity at 210 nm: 96%.

4-((((1S*,2R*)-2-(3’-((2,2,3,3,4,4,4-heptafluorobutoxy)methyl)-[1,1’-biphenyl]-4-yl)cyclopropyl)amino)methyl)benzenesulfonamide rac. (AAB252)

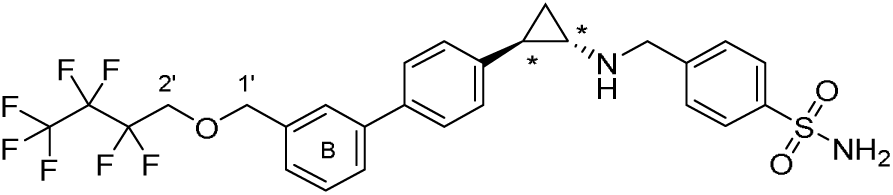

1H NMR (400 MHz, Methanol-d4) δ 8.03 – 7.92 (m, 2H, o-benzyl), 7.72 – 7.64 (m, 2H, m-benzyl), 7.58 (dq, J = 7.8, 1.8 Hz, 4H, o-Ar B, H2 B, H4 B), 7.45 (t, J = 7.6 Hz, 1H, H5 B), 7.34 (d, J = 7.7 Hz, 1H, H6 B), 7.20 – 7.15 (m, 2H, m-Ar), 4.74 (s, 2H, H1’), 4.55 – 4.42 (m, 2H, NCH2), 4.10 (tt, J = 13.8, 1.5 Hz, 2H, H2’), 3.08 – 2.98 (m, 1H, H1), 2.47 – 2.37 (m, 1H, H2), 1.60 – 1.49 (m, 1H, H3a), 1.49 – 1.41 (m, 1H, H3b). HRMS calc. for C27H25F7N2O3S [M+H]+: 591.15, found: 591.1554 [M+H]+. HPLC (method B): tR = 13.336 min; UV purity at 210 nm: 97%.

(1S*,2R*)-N-(3,4-dichlorobenzyl)-2-phenylcyclopropan-1-amine rac. AAB20

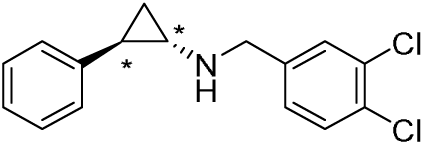

To trans 2-phenylcyclopropan-1-amine (150 mg, 1.13 mmol, 1.00 eq.) in a round-bottom flask was added acetic acid (1.10 eq.) and 3,4-dichlorobenzaldehyde (197 mg, 1.13 mmol 1.00 eq.) in DCM (8.5 mL, 0.15 M) and stirred for two hours at room temperature. Sodium triacetoxyborohydride (7.16 mg, 3.38 mmol, 3.00 eq.) was added in portions and the reaction mixture was then stirred for a further 2 h before the reaction was quenched with 10 mL of 5% sodium hydrogen carbonate solution. The organic phase was separated, and the aqueous phase was extracted three times with DCM. The organic layers were combined, dried with sodium sulfate, filtrated and concentrated under reduced pressure before being purified by column chromatography over silica gel (n-heptane/EtOAc; 20-80%). The white product was then purified by preparative HPLC method B to afford the product as a pure white TFA salt (140 mg, 43 %). 1H NMR (400 MHz, DMSO-d6) δ 9.45 (s, 2H, -NH2+-), 7.78 (d, J = 2.0 Hz, 1H, H2 benzyl), 7.70 (d, J = 8.3 Hz, 1H, H5 benzyl), 7.47 (dd, J = 8.3, 2.1 Hz, 1H, H6 benzyl), 7.32 – 7.26 (m, 2H, H3 Ar, H5 Ar), 7.26 – 7.15 (m, 1H, H4 Ar), 7.15 – 7.06 (m, 2H, H2 Ar, H6 Ar), 4.42 – 4.27 (m, 2H, -NH2+CH2-), 2.95 – 2.88 (m, 1H, H1), 2.35 (ddt, J = 9.2, 6.2, 3.0 Hz, 1H, H2), 1.44 (ddd, J = 10.4, 6.4, 4.4 Hz, 1H, H3a), 1.30 (dt, J = 7.9, 6.5 Hz, 1H, H3b). APCI: calc. for C16H16Cl2N [M+H]+: 292.06, found: 292.0/293.9 [M+H]+. HPLC: tR = 19.92 min. (method A). UV-purity at 210 nm 100 %.

tert-Butyl ((1S*,2R*)-2-(4-(1-(5-((((3r,5r,7r)-adamantan-1-yl)methyl)amino)-5-oxopentyl)-1H-1,2,3-triazol-4-yl)phenyl)cyclopropyl)(3,4-dichlorobenzyl)carbamate—methane rac. (AAB362)

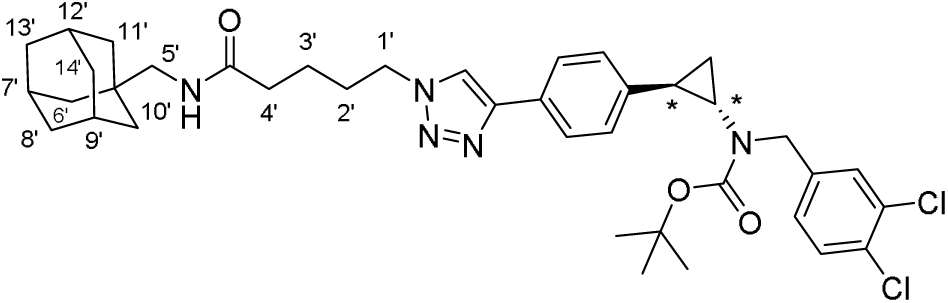

In a 10 mL-25 mL flask, the alkyne derivative (8) (25.0 mg, 0.06 mmol, 1.00 eq.) and the corresponding azide (15.0 mg, 0.06 mmol, 1.00 eq.) were suspended in a 1:1 mixture of demin. water and acetonitrile (0.05 M). Afterwards first a 1 M aqueous copper(II) sulfate solution (0.20 eq.) was added, directly followed by sodium ascorbate (0.40 eq.). (Tris(benzyltriazolylmethyl)amine (TBTA, 0.10 eq.) was dissolved in DMF (0.02 M) and then added. The reaction mixture was stirred for 2-16 h at room temperature until complete conversion of the reaction. The reaction was monitored by analytical HPLC. After extraction with dichloromethane and water, the organic layers were concentrated under reduced pressure. The crude product was immediately purified by preparative HPLC method B. 1H NMR (400 MHz, Methanol-d4) δ 8.27 (s, 1H, H5 triazole), 7.74 – 7.64 (m, 2H, o-Ar triazole), 7.48 (d, J = 8.3 Hz, 1H, H5 benzyl), 7.41 (d, J = 2.0 Hz, 1H, H2 benzyl), 7.25 – 7.18 (m, 1H, H6 benzyl), 7.18 – 7.12 (m, 2H, m-Ar triazole), 4.56 (d, J = 15.9 Hz, 1H, -NCHa-), 4.50 – 4.40 (m, 3H, -NCHb-, H1’), 2.85 (s, 2H, H5’), 2.70 (ddd, J = 7.7, 4.6, 3.4 Hz, 1H, H1), 2.27 (t, J = 7.4 Hz, 2H, H4’), 2.21 (ddd, J = 9.9, 6.5, 3.4 Hz, 1H, H2), 2.04 – 1.94 (m, 2H, H2’), 1.91 (d, J = 4.1 Hz, 3H, H7’, H9’, H12’), 1.77 – 1.57 (m, 8H, H3’, H8’, H13’, H14’), 1.48 (d, J = 2.8 Hz, 6H, H6’, H10’, H11’), 1.43 (s, 9H, C(CH3)3), 1.37 (ddd, J = 9.8, 6.0, 4.6 Hz, 1H, H3a), 1.33 – 1.18 (m, 1H, H3b). HRMS calc. for C39H50Cl2N5O3 [M+H]+: 706.33, found: 706.3293 [M+H]+. HPLC (method A): tR = 20.788 min; UV purity at 210 nm: 100%.

