## Supplementary Table 1 for "Inhibition of the macrophage demethylase LSD1 reverses *Leishmania amazonensis*-induced transcriptomic changes and causes a decrease in parasite load"

| Ensembl_Id | Mouse_Gene name |
|---|---|
| ENSMUSG00000020831 | 0610010K14Rik |
| ENSMUSG00000052595 | A1cf |
| ENSMUSG00000035234 | Abraxas1 |
| ENSMUSG00000030965 | Abraxas2 |
| ENSMUSG00000022185 | Acin1 |
| ENSMUSG00000029580 | Actb |
| ENSMUSG00000027671 | Actl6a |
| ENSMUSG00000029712 | Actl6b |
| ENSMUSG00000056367 | Actr3b |
| ENSMUSG00000037761 | Actr5 |
| ENSMUSG00000019948 | Actr6 |
| ENSMUSG00000015971 | Actr8 |
| ENSMUSG00000051149 | Adnp |
| ENSMUSG00000030232 | Aebp2 |
| ENSMUSG00000040627 | Aicda |
| ENSMUSG00000000731 | Aire |
| ENSMUSG00000079036 | Alkbh1 |
| ENSMUSG00000039754 | Alkbh4 |
| ENSMUSG00000042650 | Alkbh5 |
| ENSMUSG00000032249 | Anp32a |
| ENSMUSG00000028333 | Anp32b |
| ENSMUSG00000015749 | Anp32e |
| ENSMUSG00000037032 | Apbb1 |
| ENSMUSG00000035960 | Apex1 |
| ENSMUSG00000040613 | Apobec1 |
| ENSMUSG00000040694 | Apobec2 |
| ENSMUSG00000009585 | Apobec3 |
| ENSMUSG00000007880 | Arid1a |
| ENSMUSG00000069729 | Arid1b |
| ENSMUSG00000033237 | Arid2 |
| ENSMUSG00000048118 | Arid4a |
| ENSMUSG00000039219 | Arid4b |
| ENSMUSG00000055116 | Arntl |
| ENSMUSG00000018909 | Arrb1 |
| ENSMUSG00000019857 | Asf1a |
| ENSMUSG00000005470 | Asf1b |
| ENSMUSG00000028053 | Ash1l |
| ENSMUSG00000031575 | Ash2l |
| ENSMUSG00000042548 | Asxl1 |
| ENSMUSG00000037486 | Asxl2 |
| ENSMUSG00000045215 | Asxl3 |
| ENSMUSG00000022360 | Atad2 |
| ENSMUSG00000052812 | Atad2b |
| ENSMUSG00000027104 | Atf2 |
| ENSMUSG00000030213 | Atf7ip |
| ENSMUSG00000034218 | Atm |
| ENSMUSG00000004263 | Atn1 |
| ENSMUSG00000032409 | Atr |
| ENSMUSG00000031229 | Atrx |

| | |
|---|---|
| ENSMUSG00000021738 | Atxn7 |
| ENSMUSG00000059995 | Atxn7l3 |
| ENSMUSG00000027496 | Aurka |
| ENSMUSG00000020897 | Aurkb |
| ENSMUSG00000070837 | Aurkc |
| ENSMUSG00000031820 | Babam1 |
| ENSMUSG00000052139 | Babam2 |
| ENSMUSG00000040007 | Bahd1 |
| ENSMUSG00000025316 | Banp |
| ENSMUSG00000021901 | Bap1 |
| ENSMUSG00000026196 | Bard1 |
| ENSMUSG00000035021 | Baz1a |
| ENSMUSG00000002748 | Baz1b |
| ENSMUSG00000040054 | Baz2a |
| ENSMUSG00000026987 | Baz2b |
| ENSMUSG00000040363 | Bcor |
| ENSMUSG00000036959 | Bcorl1 |
| ENSMUSG00000026739 | Bmi1 |
| ENSMUSG00000040481 | Bptf |
| ENSMUSG00000017146 | Brca1 |
| ENSMUSG00000041147 | Brca2 |
| ENSMUSG00000031201 | Brcc3 |
| ENSMUSG00000022387 | Brd1 |
| ENSMUSG00000024335 | Brd2 |
| ENSMUSG00000026918 | Brd3 |
| ENSMUSG00000024002 | Brd4 |
| ENSMUSG00000031660 | Brd7 |
| ENSMUSG00000003778 | Brd8 |
| ENSMUSG00000057649 | Brd9 |
| ENSMUSG00000029279 | Brdt |
| ENSMUSG00000080268 | Brms1 |
| ENSMUSG00000012076 | Brms1l |
| ENSMUSG00000001632 | Brpf1 |
| ENSMUSG00000063952 | Brpf3 |
| ENSMUSG00000022914 | Brwd1 |
| ENSMUSG00000063663 | Brwd3 |
| ENSMUSG00000027379 | Bub1 |
| ENSMUSG00000032185 | Carm1 |
| ENSMUSG00000020659 | Cbll1 |
| ENSMUSG00000018666 | Cbx1 |
| ENSMUSG00000025577 | Cbx2 |
| ENSMUSG00000029836 | Cbx3 |
| ENSMUSG00000039989 | Cbx4 |
| ENSMUSG00000009575 | Cbx5 |
| ENSMUSG00000089715 | Cbx6 |
| ENSMUSG00000053411 | Cbx7 |
| ENSMUSG00000025578 | Cbx8 |
| ENSMUSG00000017499 | Cdc6 |
| ENSMUSG00000026361 | Cdc73 |
| ENSMUSG00000019942 | Cdk1 |

| | |
|---|---|
| ENSMUSG00000020015 | Cdk17 |
| ENSMUSG00000025358 | Cdk2 |
| ENSMUSG00000092300 | Cdk3 |
| ENSMUSG00000028969 | Cdk5 |
| ENSMUSG00000069089 | Cdk7 |
| ENSMUSG00000009555 | Cdk9 |
| ENSMUSG00000059288 | Cdyl |
| ENSMUSG00000031758 | Cdyl2 |
| ENSMUSG00000071226 | Cecr2 |
| ENSMUSG00000005506 | Celf1 |
| ENSMUSG00000002107 | Celf2 |
| ENSMUSG00000028137 | Celf3 |
| ENSMUSG00000024268 | Celf4 |
| ENSMUSG00000034818 | Celf5 |
| ENSMUSG00000032297 | Celf6 |
| ENSMUSG00000029253 | Cenpc1 |
| ENSMUSG00000002835 | Chaf1a |
| ENSMUSG00000022945 | Chaf1b |
| ENSMUSG00000023852 | Chd1 |
| ENSMUSG00000028089 | Chd1l |
| ENSMUSG00000078671 | Chd2 |
| ENSMUSG00000018474 | Chd3 |
| ENSMUSG00000063870 | Chd4 |
| ENSMUSG00000005045 | Chd5 |
| ENSMUSG00000057133 | Chd6 |
| ENSMUSG00000041235 | Chd7 |
| ENSMUSG00000053754 | Chd8 |
| ENSMUSG00000056608 | Chd9 |
| ENSMUSG00000032113 | Chek1 |
| ENSMUSG00000068391 | Chrac1 |
| ENSMUSG00000001017 | Chtop |
| ENSMUSG00000025199 | Chuk |
| ENSMUSG00000041777 | Cir1 |
| ENSMUSG00000029516 | Cit |
| ENSMUSG00000025439 | Clns1a |
| ENSMUSG00000029238 | Clock |
| ENSMUSG00000035403 | Crb2 |
| ENSMUSG00000022521 | Crebbp |
| ENSMUSG00000074698 | Csnk2a1 |
| ENSMUSG00000037373 | Ctbp1 |
| ENSMUSG00000030970 | Ctbp2 |
| ENSMUSG00000005698 | Ctcf |
| ENSMUSG00000070495 | Ctcfl |
| ENSMUSG00000005609 | Ctr9 |
| ENSMUSG00000029686 | Cul1 |
| ENSMUSG00000024231 | Cul2 |
| ENSMUSG00000004364 | Cul3 |
| ENSMUSG00000031446 | Cul4a |
| ENSMUSG00000031095 | Cul4b |
| ENSMUSG00000032030 | Cul5 |

| | |
|---|---|
| ENSMUSG00000024560 | Cxxc1 |
| ENSMUSG00000034974 | Dapk3 |
| ENSMUSG00000002307 | Daxx |
| ENSMUSG00000024740 | Ddb1 |
| ENSMUSG00000002109 | Ddb2 |
| ENSMUSG00000055065 | Ddx17 |
| ENSMUSG00000020075 | Ddx21 |
| ENSMUSG00000020719 | Ddx5 |
| ENSMUSG00000020076 | Ddx50 |
| ENSMUSG00000021377 | Dek |
| ENSMUSG00000042699 | Dhx9 |
| ENSMUSG00000009640 | Dmap1 |
| ENSMUSG00000026740 | Dnajc1 |
| ENSMUSG00000029014 | Dnajc2 |
| ENSMUSG00000044595 | Dnd1 |
| ENSMUSG00000004099 | Dnmt1 |
| ENSMUSG00000020661 | Dnmt3a |
| ENSMUSG00000027478 | Dnmt3b |
| ENSMUSG00000000730 | Dnmt3l |
| ENSMUSG00000039756 | Dnttip2 |
| ENSMUSG00000061589 | Dot1l |
| ENSMUSG00000030584 | Dpf1 |
| ENSMUSG00000024826 | Dpf2 |
| ENSMUSG00000021221 | Dpf3 |
| ENSMUSG00000046323 | Dppa3 |
| ENSMUSG00000024067 | Dpy30 |
| ENSMUSG00000029265 | Dr1 |
| ENSMUSG00000049502 | Dtx3l |
| ENSMUSG00000064061 | Dzip3 |
| ENSMUSG00000057469 | E2F6 |
| ENSMUSG00000030619 | Eed |
| ENSMUSG00000080115 | Eef1akmt3 |
| ENSMUSG00000115219 | Eef1akmt4 |
| ENSMUSG00000036893 | Ehmt1 |
| ENSMUSG00000013787 | Ehmt2 |
| ENSMUSG00000091337 | Eid1 |
| ENSMUSG00000046058 | Eid2 |
| ENSMUSG00000070705 | Eid2b |
| ENSMUSG00000025580 | Eif4a3 |
| ENSMUSG00000028431 | Elp1 |
| ENSMUSG00000024271 | Elp2 |
| ENSMUSG00000022031 | Elp3 |
| ENSMUSG00000027167 | Elp4 |
| ENSMUSG00000018565 | Elp5 |
| ENSMUSG00000054836 | Elp6 |
| ENSMUSG00000035401 | Emsy |
| ENSMUSG00000022338 | Eny2 |
| ENSMUSG00000055024 | Ep300 |
| ENSMUSG00000029505 | Ep400 |
| ENSMUSG00000024240 | Epc1 |

| | |
|---|---|
| ENSMUSG00000069495 | Epc2 |
| ENSMUSG00000062209 | Erbb4 |
| ENSMUSG00000054051 | Ercc6 |
| ENSMUSG00000034321 | Exosc1 |
| ENSMUSG00000039356 | Exosc2 |
| ENSMUSG00000028322 | Exosc3 |
| ENSMUSG00000034259 | Exosc4 |
| ENSMUSG00000061286 | Exosc5 |
| ENSMUSG00000109941 | Exosc6 |
| ENSMUSG00000025785 | Exosc7 |
| ENSMUSG00000027752 | Exosc8 |
| ENSMUSG00000027714 | Exosc9 |
| ENSMUSG00000025932 | Eya1 |
| ENSMUSG00000017897 | Eya2 |
| ENSMUSG00000028886 | Eya3 |
| ENSMUSG00000010461 | Eya4 |
| ENSMUSG00000006920 | Ezh1 |
| ENSMUSG00000029687 | Ezh2 |
| ENSMUSG00000046865 | Fbl |
| ENSMUSG00000042423 | Fbrs |
| ENSMUSG00000043323 | Fbrsl1 |
| ENSMUSG00000035451 | Foxa1 |
| ENSMUSG00000044167 | Foxo1 |
| ENSMUSG00000030067 | Foxp1 |
| ENSMUSG00000029563 | Foxp2 |
| ENSMUSG00000039521 | Foxp3 |
| ENSMUSG00000023991 | Foxp4 |
| ENSMUSG00000055932 | Fto |
| ENSMUSG00000036390 | Gadd45a |
| ENSMUSG00000015312 | Gadd45b |
| ENSMUSG00000021453 | Gadd45g |
| ENSMUSG00000007415 | Gatad1 |
| ENSMUSG00000036180 | Gatad2a |
| ENSMUSG00000042390 | Gatad2b |
| ENSMUSG00000029275 | Gfi1 |
| ENSMUSG00000026815 | Gfi1b |
| ENSMUSG00000022536 | Glyr1 |
| ENSMUSG00000100622 | Gm20379 |
| ENSMUSG00000031822 | Gse1 |
| ENSMUSG00000060261 | Gtf2i |
| ENSMUSG00000035666 | Gtf3c4 |
| ENSMUSG00000050107 | Haspin |
| ENSMUSG00000027018 | Hat1 |
| ENSMUSG00000031386 | Hcfc1 |
| ENSMUSG00000020246 | Hcfc2 |
| ENSMUSG00000028800 | Hdac1 |
| ENSMUSG00000062906 | Hdac10 |
| ENSMUSG00000034245 | Hdac11 |
| ENSMUSG00000019777 | Hdac2 |
| ENSMUSG00000024454 | Hdac3 |

| | |
|---|---|
| ENSMUSG00000026313 | Hdac4 |
| ENSMUSG00000008855 | Hdac5 |
| ENSMUSG00000031161 | Hdac6 |
| ENSMUSG00000022475 | Hdac7 |
| ENSMUSG00000067567 | Hdac8 |
| ENSMUSG00000004698 | Hdac9 |
| ENSMUSG00000004897 | Hdgf |
| ENSMUSG00000002833 | Hdgfl2 |
| ENSMUSG00000025001 | Hells |
| ENSMUSG00000036450 | Hif1an |
| ENSMUSG00000032119 | Hinfp |
| ENSMUSG00000022702 | Hira |
| ENSMUSG00000042606 | Hirip3 |
| ENSMUSG00000044783 | Hjurp |
| ENSMUSG00000040820 | Hlcs |
| ENSMUSG00000002428 | Hltf |
| ENSMUSG00000032329 | Hmg20a |
| ENSMUSG00000020232 | Hmg20b |
| ENSMUSG00000066551 | Hmgb1 |
| ENSMUSG00000040681 | Hmgn1 |
| ENSMUSG00000003038 | Hmgn2 |
| ENSMUSG00000066456 | Hmgn3 |
| ENSMUSG00000031245 | Hmgn5 |
| ENSMUSG00000015165 | Hnrnpl |
| ENSMUSG00000059208 | Hnrnpm |
| ENSMUSG00000039630 | Hnrnpu |
| ENSMUSG00000028759 | Hp1bp3 |
| ENSMUSG00000022096 | Hr |
| ENSMUSG00000091971 | Hspa1a |
| ENSMUSG00000090877 | Hspa1b |
| ENSMUSG00000025261 | Huwe1 |
| ENSMUSG00000018654 | Ikzf1 |
| ENSMUSG00000018168 | Ikzf3 |
| ENSMUSG00000045969 | Ing1 |
| ENSMUSG00000063049 | Ing2 |
| ENSMUSG00000029670 | Ing3 |
| ENSMUSG00000030330 | Ing4 |
| ENSMUSG00000026283 | Ing5 |
| ENSMUSG00000034154 | Ino80 |
| ENSMUSG00000030034 | Ino80b |
| ENSMUSG00000047989 | Ino80c |
| ENSMUSG00000040865 | Ino80d |
| ENSMUSG00000030689 | Ino80e |
| ENSMUSG00000025764 | Jade1 |
| ENSMUSG00000020387 | Jade2 |
| ENSMUSG00000037315 | Jade3 |
| ENSMUSG00000024789 | Jak2 |
| ENSMUSG00000038518 | Jarid2 |
| ENSMUSG00000034271 | Jdp2 |
| ENSMUSG00000037876 | Jmjd1c |

| | |
|---|---|
| ENSMUSG00000036819 | Jmjd4 |
| ENSMUSG00000056962 | Jmjd6 |
| ENSMUSG00000098789 | Jmjd7 |
| ENSMUSG00000025736 | Jmjd8 |
| ENSMUSG00000018412 | Kansl1 |
| ENSMUSG00000022992 | Kansl2 |
| ENSMUSG00000010453 | Kansl3 |
| ENSMUSG00000020918 | Kat2a |
| ENSMUSG00000000708 | Kat2b |
| ENSMUSG00000024926 | Kat5 |
| ENSMUSG00000031540 | Kat6a |
| ENSMUSG00000021767 | Kat6b |
| ENSMUSG00000038909 | Kat7 |
| ENSMUSG00000030801 | Kat8 |
| ENSMUSG00000036940 | Kdm1a |
| ENSMUSG00000038080 | Kdm1b |
| ENSMUSG00000054611 | Kdm2a |
| ENSMUSG00000029475 | Kdm2b |
| ENSMUSG00000053470 | Kdm3a |
| ENSMUSG00000038773 | Kdm3b |
| ENSMUSG00000033326 | Kdm4a |
| ENSMUSG00000024201 | Kdm4b |
| ENSMUSG00000028397 | Kdm4c |
| ENSMUSG00000053914 | Kdm4d |
| ENSMUSG00000030180 | Kdm5a |
| ENSMUSG00000042207 | Kdm5b |
| ENSMUSG00000025332 | Kdm5c |
| ENSMUSG00000056673 | Kdm5d |
| ENSMUSG00000037369 | Kdm6a |
| ENSMUSG00000018476 | Kdm6b |
| ENSMUSG00000042599 | Kdm7a |
| ENSMUSG00000030752 | Kdm8 |
| ENSMUSG00000003308 | Keap1 |
| ENSMUSG00000028790 | Khdrbs1 |
| ENSMUSG00000002028 | Kmt2a |
| ENSMUSG00000006307 | Kmt2b |
| ENSMUSG00000038056 | Kmt2c |
| ENSMUSG00000048154 | Kmt2d |
| ENSMUSG00000029004 | Kmt2e |
| ENSMUSG00000049327 | Kmt5a |
| ENSMUSG00000045098 | Kmt5b |
| ENSMUSG00000059851 | Kmt5c |
| ENSMUSG00000035576 | L3mbtl1 |
| ENSMUSG00000022394 | L3mbtl2 |
| ENSMUSG00000039089 | L3mbtl3 |
| ENSMUSG00000041565 | L3mbtl4 |
| ENSMUSG00000057421 | Las1l |
| ENSMUSG00000004880 | Lbr |
| ENSMUSG00000042487 | Leo1 |
| ENSMUSG00000029703 | Lrwd1 |

| | |
|---|---|
| ENSMUSG00000028609 | Magoh |
| ENSMUSG00000028284 | Map3k7 |
| ENSMUSG00000032577 | Mapkapk3 |
| ENSMUSG00000026779 | Mastl |
| ENSMUSG00000059436 | Max |
| ENSMUSG00000030678 | Maz |
| ENSMUSG00000024561 | Mbd1 |
| ENSMUSG00000024513 | Mbd2 |
| ENSMUSG00000035478 | Mbd3 |
| ENSMUSG00000030322 | Mbd4 |
| ENSMUSG00000036792 | Mbd5 |
| ENSMUSG00000025409 | Mbd6 |
| ENSMUSG00000021028 | Mbip |
| ENSMUSG00000027763 | Mbnl1 |
| ENSMUSG00000036109 | Mbnl3 |
| ENSMUSG00000059474 | Mbtd1 |
| ENSMUSG00000037570 | Mcrs1 |
| ENSMUSG00000061607 | Mdc1 |
| ENSMUSG00000028863 | Meaf6 |
| ENSMUSG00000031393 | Mecp2 |
| ENSMUSG00000024947 | Men1 |
| ENSMUSG00000040113 | Mettl11b |
| ENSMUSG00000026694 | Mettl13 |
| ENSMUSG00000028114 | Mettl14 |
| ENSMUSG00000010554 | Mettl16 |
| ENSMUSG00000025956 | Mettl21a |
| ENSMUSG00000022160 | Mettl3 |
| ENSMUSG00000055660 | Mettl4 |
| ENSMUSG00000033943 | Mga |
| ENSMUSG00000026743 | Miit10 |
| ENSMUSG00000022978 | Mis18a |
| ENSMUSG00000047534 | Mis18bp1 |
| ENSMUSG00000024212 | Mllt1 |
| ENSMUSG00000038437 | Mllt6 |
| ENSMUSG00000062270 | Morf4l1 |
| ENSMUSG00000031422 | Morf4l2 |
| ENSMUSG00000002227 | Mov10 |
| ENSMUSG00000079184 | Mphosph8 |
| ENSMUSG00000003199 | Mpnd |
| ENSMUSG00000027569 | Mrgbp |
| ENSMUSG00000005370 | Msh6 |
| ENSMUSG00000052915 | Msl1 |
| ENSMUSG00000066415 | Msl2 |
| ENSMUSG00000031358 | Msl3 |
| ENSMUSG00000032591 | Mst1 |
| ENSMUSG00000021144 | Mta1 |
| ENSMUSG00000071646 | Mta2 |
| ENSMUSG00000055817 | Mta3 |
| ENSMUSG00000029267 | Mtf2 |
| ENSMUSG00000040463 | Mybbp1a |

| | |
|---|---|
| ENSMUSG00000017774 | Myo1c |
| ENSMUSG00000062627 | Mysm1 |
| ENSMUSG00000005982 | Naa60 |
| ENSMUSG00000058799 | Nap1l1 |
| ENSMUSG00000082229 | Nap1l2 |
| ENSMUSG00000059119 | Nap1l4 |
| ENSMUSG00000028693 | Nasp |
| ENSMUSG00000027185 | Nat10 |
| ENSMUSG00000028224 | Nbn |
| ENSMUSG00000026234 | Ncl |
| ENSMUSG00000020647 | Ncoa1 |
| ENSMUSG00000005886 | Ncoa2 |
| ENSMUSG00000027678 | Ncoa3 |
| ENSMUSG00000038369 | Ncoa6 |
| ENSMUSG00000018501 | Ncor1 |
| ENSMUSG00000029478 | Ncor2 |
| ENSMUSG00000026749 | Nek6 |
| ENSMUSG00000034290 | Nek9 |
| ENSMUSG00000042185 | Nfrkb |
| ENSMUSG00000020248 | Nfyb |
| ENSMUSG00000032897 | Nfyc |
| ENSMUSG00000022141 | Nipbl |
| ENSMUSG00000095567 | Noc2l |
| ENSMUSG00000026077 | Npas2 |
| ENSMUSG00000057113 | Npm1 |
| ENSMUSG00000047911 | Npm2 |
| ENSMUSG00000021488 | Nsd1 |
| ENSMUSG00000057406 | Nsd2 |
| ENSMUSG00000054823 | Nsd3 |
| ENSMUSG00000062510 | Nsl1 |
| ENSMUSG00000037958 | Nsrp1 |
| ENSMUSG00000021595 | Nsun2 |
| ENSMUSG00000026707 | Nsun6 |
| ENSMUSG00000026857 | Ntmt1 |
| ENSMUSG00000063550 | Nup98 |
| ENSMUSG00000034160 | Ogt |
| ENSMUSG00000072980 | Oip5 |
| ENSMUSG00000025329 | Padi1 |
| ENSMUSG00000028927 | Padi2 |
| ENSMUSG00000025328 | Padi3 |
| ENSMUSG00000025330 | Padi4 |
| ENSMUSG00000003437 | Paf1 |
| ENSMUSG00000030680 | Pagr1 |
| ENSMUSG00000022781 | Pak2 |
| ENSMUSG00000021911 | Parg |
| ENSMUSG00000026496 | Parp1 |
| ENSMUSG00000036023 | Parp2 |
| ENSMUSG00000023249 | Parp3 |
| ENSMUSG00000002221 | Paxip1 |
| ENSMUSG00000022033 | Pbk |

| | |
|---|---|
| ENSMUSG00000042323 | Pbrm1 |
| ENSMUSG00000069678 | Pcgf1 |
| ENSMUSG00000018537 | Pcgf2 |
| ENSMUSG00000033623 | Pcgf3 |
| ENSMUSG00000024805 | Pcgf5 |
| ENSMUSG00000025050 | Pcgf6 |
| ENSMUSG00000027342 | Pcna |
| ENSMUSG00000049225 | Pdp1 |
| ENSMUSG00000018921 | Pelp1 |
| ENSMUSG00000098387 | Pet117 |
| ENSMUSG00000040669 | Phc1 |
| ENSMUSG00000028796 | Phc2 |
| ENSMUSG00000037652 | Phc3 |
| ENSMUSG00000024193 | Phf1 |
| ENSMUSG00000023883 | Phf10 |
| ENSMUSG00000037791 | Phf12 |
| ENSMUSG00000047777 | Phf13 |
| ENSMUSG00000029629 | Phf14 |
| ENSMUSG00000026873 | Phf19 |
| ENSMUSG00000038025 | Phf2 |
| ENSMUSG00000038116 | Phf20 |
| ENSMUSG00000072501 | Phf20l1 |
| ENSMUSG00000058318 | Phf21a |
| ENSMUSG00000041229 | Phf8 |
| ENSMUSG00000032253 | Phip |
| ENSMUSG00000036912 | Piwil4 |
| ENSMUSG00000032294 | Pkm |
| ENSMUSG00000057672 | Pkn1 |
| ENSMUSG00000038902 | Pogz |
| ENSMUSG00000028394 | Pole3 |
| ENSMUSG00000029167 | Ppargc1a |
| ENSMUSG00000029147 | Ppm1g |
| ENSMUSG00000020349 | Ppp2ca |
| ENSMUSG00000030697 | Ppp4c |
| ENSMUSG00000052144 | Ppp4r2 |
| ENSMUSG00000041846 | Ppp4r3a |
| ENSMUSG00000020463 | Ppp4r3b |
| ENSMUSG00000031157 | Pqbp1 |
| ENSMUSG00000038151 | Prdm1 |
| ENSMUSG00000075028 | Prdm11 |
| ENSMUSG00000079466 | Prdm12 |
| ENSMUSG00000040478 | Prdm13 |
| ENSMUSG00000042414 | Prdm14 |
| ENSMUSG00000039410 | Prdm16 |
| ENSMUSG00000057637 | Prdm2 |
| ENSMUSG00000035529 | Prdm4 |
| ENSMUSG00000029913 | Prdm5 |
| ENSMUSG00000069378 | Prdm6 |
| ENSMUSG00000035456 | Prdm8 |
| ENSMUSG00000051977 | Prdm9 |

| | |
|---|---|
| ENSMUSG00000050697 | Prkaa1 |
| ENSMUSG00000028518 | Prkaa2 |
| ENSMUSG00000029513 | Prkab1 |
| ENSMUSG00000038205 | Prkab2 |
| ENSMUSG00000067713 | Prkag1 |
| ENSMUSG00000028944 | Prkag2 |
| ENSMUSG00000006542 | Prkag3 |
| ENSMUSG00000050965 | Prkca |
| ENSMUSG00000052889 | Prkcb |
| ENSMUSG00000021948 | Prkcd |
| ENSMUSG00000022672 | Prkdc |
| ENSMUSG00000109324 | Prmt1 |
| ENSMUSG00000020230 | Prmt2 |
| ENSMUSG00000030505 | Prmt3 |
| ENSMUSG00000023110 | Prmt5 |
| ENSMUSG00000049300 | Prmt6 |
| ENSMUSG00000060098 | Prmt7 |
| ENSMUSG00000030350 | Prmt8 |
| ENSMUSG00000037134 | Prmt9 |
| ENSMUSG00000008373 | Prpf31 |
| ENSMUSG00000030822 | Prr14 |
| ENSMUSG00000028484 | Psip1 |
| ENSMUSG00000006498 | Ptbp1 |
| ENSMUSG00000002524 | Puf60 |
| ENSMUSG00000020156 | Pwwp3a |
| ENSMUSG00000027323 | Rad51 |
| ENSMUSG00000078773 | Rad54b |
| ENSMUSG00000028702 | Rad54l |
| ENSMUSG00000040661 | Rad54l2 |
| ENSMUSG00000061311 | Rag1 |
| ENSMUSG00000032864 | Rag2 |
| ENSMUSG00000062115 | Rai1 |
| ENSMUSG00000037992 | Rara |
| ENSMUSG00000022105 | Rb1 |
| ENSMUSG00000057236 | Rbbp4 |
| ENSMUSG00000026439 | Rbbp5 |
| ENSMUSG00000031353 | Rbbp7 |
| ENSMUSG00000008658 | Rbfox1 |
| ENSMUSG00000032940 | Rbm11 |
| ENSMUSG00000048109 | Rbm15 |
| ENSMUSG00000074102 | Rbm15b |
| ENSMUSG00000037197 | Rbm17 |
| ENSMUSG00000038132 | Rbm24 |
| ENSMUSG00000010608 | Rbm25 |
| ENSMUSG00000094936 | Rbm4 |
| ENSMUSG00000032580 | Rbm5 |
| ENSMUSG00000042396 | Rbm7 |
| ENSMUSG00000038374 | Rbm8a |
| ENSMUSG00000022400 | Rbx1 |
| ENSMUSG00000028896 | Rcc1 |

| | |
|---|---|
| ENSMUSG00000037896 | Rcor1 |
| ENSMUSG00000037395 | Rcor3 |
| ENSMUSG00000029249 | Rest |
| ENSMUSG00000024325 | Ring1 |
| ENSMUSG00000046791 | Riox1 |
| ENSMUSG00000022724 | Riox2 |
| ENSMUSG00000056537 | Rlim |
| ENSMUSG00000035367 | Rmi1 |
| ENSMUSG00000014074 | Rnf168 |
| ENSMUSG00000026484 | Rnf2 |
| ENSMUSG00000028309 | Rnf20 |
| ENSMUSG00000030816 | Rnf40 |
| ENSMUSG00000090083 | Rnf8 |
| ENSMUSG00000034681 | Rnps1 |
| ENSMUSG00000031309 | Rps6ka3 |
| ENSMUSG00000118668 | Rps6ka4 |
| ENSMUSG00000021180 | Rps6ka5 |
| ENSMUSG00000051169 | Rpusd3 |
| ENSMUSG00000030888 | Rrp8 |
| ENSMUSG00000035623 | Rsf1 |
| ENSMUSG00000034544 | Rsrc1 |
| ENSMUSG00000030079 | Ruvbl1 |
| ENSMUSG00000003868 | Ruvbl2 |
| ENSMUSG00000072872 | Rybp |
| ENSMUSG00000071054 | Safb |
| ENSMUSG00000024260 | Sap130 |
| ENSMUSG00000021963 | Sap18 |
| ENSMUSG00000061104 | Sap18b |
| ENSMUSG00000079165 | Sap25 |
| ENSMUSG00000031609 | Sap30 |
| ENSMUSG00000020519 | Sap30l |
| ENSMUSG00000023927 | Satb1 |
| ENSMUSG00000038331 | Satb2 |
| ENSMUSG00000000085 | Scmh1 |
| ENSMUSG00000000037 | Scml2 |
| ENSMUSG00000044770 | Scml4 |
| ENSMUSG00000033075 | Senp1 |
| ENSMUSG00000005204 | Senp3 |
| ENSMUSG00000054766 | Set |
| ENSMUSG00000042308 | Setd1a |
| ENSMUSG00000038384 | Setd1b |
| ENSMUSG00000044791 | Setd2 |
| ENSMUSG00000056770 | Setd3 |
| ENSMUSG00000022948 | Setd4 |
| ENSMUSG00000034269 | Setd5 |
| ENSMUSG00000031671 | Setd6 |
| ENSMUSG00000037111 | Setd7 |
| ENSMUSG00000015697 | Setdb1 |
| ENSMUSG00000071350 | Setdb2 |
| ENSMUSG00000034639 | Setmar |

| | |
|---|---|
| ENSMUSG00000025982 | Sf3b1 |
| ENSMUSG00000033732 | Sf3b3 |
| ENSMUSG00000006527 | Sfmbt1 |
| ENSMUSG00000061186 | Sfmbt2 |
| ENSMUSG00000028820 | Sfpq |
| ENSMUSG00000029439 | Sfswap |
| ENSMUSG00000030714 | Sgf29 |
| ENSMUSG00000090112 | Shprh |
| ENSMUSG00000042557 | Sin3a |
| ENSMUSG00000031622 | Sin3b |
| ENSMUSG00000020063 | Sirt1 |
| ENSMUSG00000015149 | Sirt2 |
| ENSMUSG00000025486 | Sirt3 |
| ENSMUSG00000034748 | Sirt6 |
| ENSMUSG00000025138 | Sirt7 |
| ENSMUSG00000036309 | Skp1 |
| ENSMUSG00000021597 | Slf1 |
| ENSMUSG00000020409 | Slu7 |
| ENSMUSG00000031099 | Smarca1 |
| ENSMUSG00000024921 | Smarca2 |
| ENSMUSG00000032187 | Smarca4 |
| ENSMUSG00000031715 | Smarca5 |
| ENSMUSG00000029920 | Smarcad1 |
| ENSMUSG00000039354 | Smarcal1 |
| ENSMUSG00000000902 | Smarcb1 |
| ENSMUSG00000032481 | Smarcc1 |
| ENSMUSG00000025369 | Smarcc2 |
| ENSMUSG00000023018 | Smarcd1 |
| ENSMUSG00000078619 | Smarcd2 |
| ENSMUSG00000028949 | Smarcd3 |
| ENSMUSG00000037935 | Smarce1 |
| ENSMUSG00000055027 | Smyd1 |
| ENSMUSG00000026603 | Smyd2 |
| ENSMUSG00000055067 | Smyd3 |
| ENSMUSG00000018809 | Smyd4 |
| ENSMUSG00000033706 | Smyd5 |
| ENSMUSG00000022676 | Snai2 |
| ENSMUSG00000001280 | Sp1 |
| ENSMUSG00000026222 | Sp100 |
| ENSMUSG00000070031 | Sp140 |
| ENSMUSG00000040761 | Spen |
| ENSMUSG00000057522 | Spop |
| ENSMUSG00000053877 | Srcap |
| ENSMUSG00000063919 | Srrm4 |
| ENSMUSG00000018379 | Srsf1 |
| ENSMUSG00000028676 | Srsf10 |
| ENSMUSG00000054679 | Srsf12 |
| ENSMUSG00000071172 | Srsf3 |
| ENSMUSG00000016921 | Srsf6 |
| ENSMUSG00000039086 | Ss18l1 |

| | |
|---|---|
| ENSMUSG00000032526 | Ss18l2 |
| ENSMUSG00000027067 | Ssrp1 |
| ENSMUSG00000018209 | Stk4 |
| ENSMUSG00000066900 | Suds3 |
| ENSMUSG00000035726 | Supt16 |
| ENSMUSG00000038954 | Supt3 |
| ENSMUSG00000002052 | Supt6 |
| ENSMUSG00000053134 | Supt7l |
| ENSMUSG00000039231 | Suv39h1 |
| ENSMUSG00000026646 | Suv39h2 |
| ENSMUSG00000017548 | Suz12 |
| ENSMUSG00000032423 | Syncrip |
| ENSMUSG00000026563 | Tada1 |
| ENSMUSG00000018651 | Tada2a |
| ENSMUSG00000029196 | Tada2b |
| ENSMUSG00000048930 | Tada3 |
| ENSMUSG00000031314 | Taf1 |
| ENSMUSG00000043866 | Taf10 |
| ENSMUSG00000028899 | Taf12 |
| ENSMUSG00000037343 | Taf2 |
| ENSMUSG00000025782 | Taf3 |
| ENSMUSG00000039117 | Taf4 |
| ENSMUSG00000025049 | Taf5 |
| ENSMUSG00000038697 | Taf5l |
| ENSMUSG00000036980 | Taf6 |
| ENSMUSG00000003680 | Taf6l |
| ENSMUSG00000051316 | Taf7 |
| ENSMUSG00000023980 | Taf8 |
| ENSMUSG00000052293 | Taf9 |
| ENSMUSG00000047242 | Taf9b |
| ENSMUSG00000027630 | Tbl1xr1 |
| ENSMUSG00000034674 | Tdg |
| ENSMUSG00000022019 | Tdrd3 |
| ENSMUSG00000035517 | Tdrd7 |
| ENSMUSG00000041912 | Tdrkh |
| ENSMUSG00000047146 | Tet1 |
| ENSMUSG00000040943 | Tet2 |
| ENSMUSG00000034832 | Tet3 |
| ENSMUSG00000028345 | Tex10 |
| ENSMUSG00000038482 | Tfdp1 |
| ENSMUSG00000006335 | Tfpt |
| ENSMUSG00000043962 | Thrap3 |
| ENSMUSG00000008305 | Tle1 |
| ENSMUSG00000034771 | Tle2 |
| ENSMUSG00000024642 | Tle4 |
| ENSMUSG00000041997 | Tlk1 |
| ENSMUSG00000020694 | Tlk2 |
| ENSMUSG00000026182 | Tnp1 |
| ENSMUSG00000043050 | Tnp2 |
| ENSMUSG00000059323 | Tonsl |

| | |
|---|---|
| ENSMUSG00000020914 | Top2a |
| ENSMUSG00000017485 | Top2b |
| ENSMUSG00000043909 | Tp53bp1 |
| ENSMUSG00000022858 | Tra2b |
| ENSMUSG00000047821 | Trim16 |
| ENSMUSG00000029833 | Trim24 |
| ENSMUSG00000021326 | Trim27 |
| ENSMUSG00000005566 | Trim28 |
| ENSMUSG00000033014 | Trim33 |
| ENSMUSG00000059552 | Trp53 |
| ENSMUSG00000045482 | Trrap |
| ENSMUSG00000039826 | Trub2 |
| ENSMUSG00000047654 | Tssk6 |
| ENSMUSG00000038379 | Ttk |
| ENSMUSG00000048495 | Tyw5 |
| ENSMUSG00000030435 | U2af2 |
| ENSMUSG00000016308 | Ube2a |
| ENSMUSG00000020390 | Ube2b |
| ENSMUSG00000019927 | Ube2d1 |
| ENSMUSG00000078578 | Ube2d3 |
| ENSMUSG00000021774 | Ube2e1 |
| ENSMUSG00000039159 | Ube2h |
| ENSMUSG00000074781 | Ube2n |
| ENSMUSG00000026429 | Ube2t |
| ENSMUSG00000039473 | Ubn1 |
| ENSMUSG00000023977 | Ubr2 |
| ENSMUSG00000037487 | Ubr5 |
| ENSMUSG00000041712 | Ubr7 |
| ENSMUSG00000018189 | Uchl5 |
| ENSMUSG00000001228 | Uhrf1 |
| ENSMUSG00000024817 | Uhrf2 |
| ENSMUSG00000025878 | Uimc1 |
| ENSMUSG00000031066 | Usp11 |
| ENSMUSG00000029640 | Usp12 |
| ENSMUSG00000020124 | Usp15 |
| ENSMUSG00000025616 | Usp16 |
| ENSMUSG00000053483 | Usp21 |
| ENSMUSG00000042506 | Usp22 |
| ENSMUSG00000032376 | Usp3 |
| ENSMUSG00000033909 | Usp36 |
| ENSMUSG00000020020 | Usp44 |
| ENSMUSG00000054814 | Usp46 |
| ENSMUSG00000090115 | Usp49 |
| ENSMUSG00000022710 | Usp7 |
| ENSMUSG00000068457 | Uty |
| ENSMUSG00000022479 | Vdr |
| ENSMUSG00000040720 | Virma |
| ENSMUSG00000008958 | Vps72 |
| ENSMUSG00000021115 | Vrk1 |
| ENSMUSG00000024283 | Wac |

| | |
|---|---|
| ENSMUSG00000026917 | Wdr5 |
| ENSMUSG00000000561 | WDR77 |
| ENSMUSG00000020257 | Wdr82 |
| ENSMUSG00000029364 | Wsb2 |
| ENSMUSG00000060475 | Wtap |
| ENSMUSG00000022634 | Yaf2 |
| ENSMUSG00000041215 | Yeats2 |
| ENSMUSG00000020171 | Yeats4 |
| ENSMUSG00000035851 | Ythdc1 |
| ENSMUSG00000018326 | Ywhab |
| ENSMUSG00000020849 | Ywhae |
| ENSMUSG00000022285 | Ywhaz |
| ENSMUSG00000021264 | Yy1 |
| ENSMUSG00000066687 | Zbtb16 |
| ENSMUSG00000048047 | Zbtb33 |
| ENSMUSG00000035011 | Zbtb7a |
| ENSMUSG00000044646 | Zbtb7c |
| ENSMUSG00000022000 | Zc3h13 |
| ENSMUSG00000037108 | Zcwpw1 |
| ENSMUSG00000052056 | Zfp217 |
| ENSMUSG00000070902 | Zfp352 |
| ENSMUSG00000058881 | Zfp516 |
| ENSMUSG00000042439 | Zfp532 |
| ENSMUSG00000078796 | Zfp541 |
| ENSMUSG00000036036 | Zfp57 |
| ENSMUSG00000005621 | Zfp592 |
| ENSMUSG00000019338 | Zfp687 |
| ENSMUSG00000025529 | Zfp711 |
| ENSMUSG00000055150 | Zfp78 |
| ENSMUSG00000027582 | Zgpat |
| ENSMUSG00000022361 | Zhx1 |
| ENSMUSG00000021945 | Zmym2 |
| ENSMUSG00000031310 | Zmym3 |
| ENSMUSG00000021156 | Zmynd11 |
| ENSMUSG00000039671 | Zmynd8 |
| ENSMUSG00000059518 | Znhit1 |
| ENSMUSG00000036086 | Zranb3 |
| ENSMUSG00000039068 | Zzz3 |
