## Supplementary Table 2 for "Inhibition of the macrophage demethylase LSD1 reverses *Leishmania amazonensis*-induced transcriptomic changes and causes a decrease in parasite load"

**~~Table S2,~~ Gutierrez Sanchez et al.**

**Composition of Lysis (LBI) and Extraction (EBI) Buffers for nuclear and cytoplasmic protein extraction.**

| | | |
|---|---|---|
| **Common components among buffers** | - 25 mM β-glycerophosphate | - 1 mM TSPP |
| | - 2 mM Phenanthroline | - 0.5 mM Spermidine |
| | - 5 mM Na3VO4 | - 0.15 mM Spermine |
| | - 1 mM PMSF | |
| | - 1x Protein Inhibitor Cocktail (PIC) (cOmplete Roche tablets for Lysis Buffers and HaltTM Thermo Fisher Scientific for Extraction Buffers) | |
| **Lysis Buffer I (LBI) Activity Assay, Western Blot** | - 1 mM KCl | - 0.5 mM EDTA pH 8.0 |
| | - 20 mM Hepes pH 7.5 | - 1 mM MgCl2 |
| | - 1 mM DTT | - 0.5% NP-40 |
| **Extraction Buffer I (EBI) LSD1 Western Blot Analysis & LSD1 Activity Assay** | - 600 mM NaCl | - 1 mM MgCl2 |
| | - 25 mM Hepes pH 7.5 | - 1% Triton |
| | - 1 mM DTT | |
