## Supplementary Table 3 for "Inhibition of the macrophage demethylase LSD1 reverses *Leishmania amazonensis*-induced transcriptomic changes and causes a decrease in parasite load"

### Slide 1
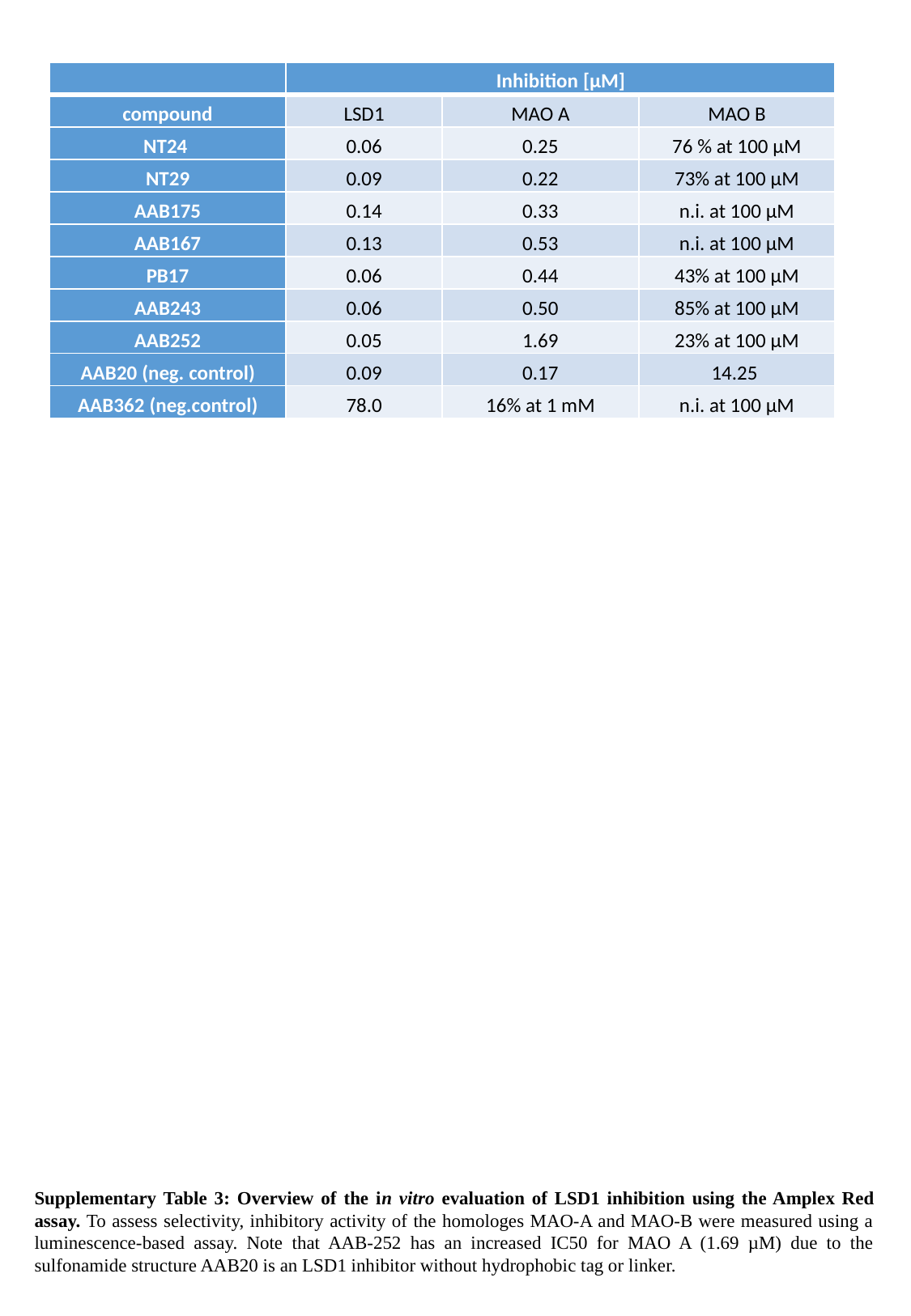

| | Inhibition [µM] | | |
| --- | --- | --- | --- |
| compound | LSD1 | MAO A | MAO B |
| NT24 | 0.06 | 0.25 | 76 % at 100 µM |
| NT29 | 0.09 | 0.22 | 73% at 100 µM |
| AAB175 | 0.14 | 0.33 | n.i. at 100 µM |
| AAB167 | 0.13 | 0.53 | n.i. at 100 µM |
| PB17 | 0.06 | 0.44 | 43% at 100 µM |
| AAB243 | 0.06 | 0.50 | 85% at 100 µM |
| AAB252 | 0.05 | 1.69 | 23% at 100 µM |
| AAB20 (neg. control) | 0.09 | 0.17 | 14.25 |
| AAB362 (neg.control) | 78.0 | 16% at 1 mM | n.i. at 100 µM |
Supplementary Table 3: Overview of the in vitro evaluation of LSD1 inhibition using the Amplex Red assay. To assess selectivity, inhibitory activity of the homologes MAO-A and MAO-B were measured using a luminescence-based assay. Note that AAB-252 has an increased IC50 for MAO A (1.69 µM) due to the sulfonamide structure AAB20 is an LSD1 inhibitor without hydrophobic tag or linker.
