## Supplementary Figures for "Inhibition of the macrophage demethylase LSD1 reverses *Leishmania amazonensis*-induced transcriptomic changes and causes a decrease in parasite load"

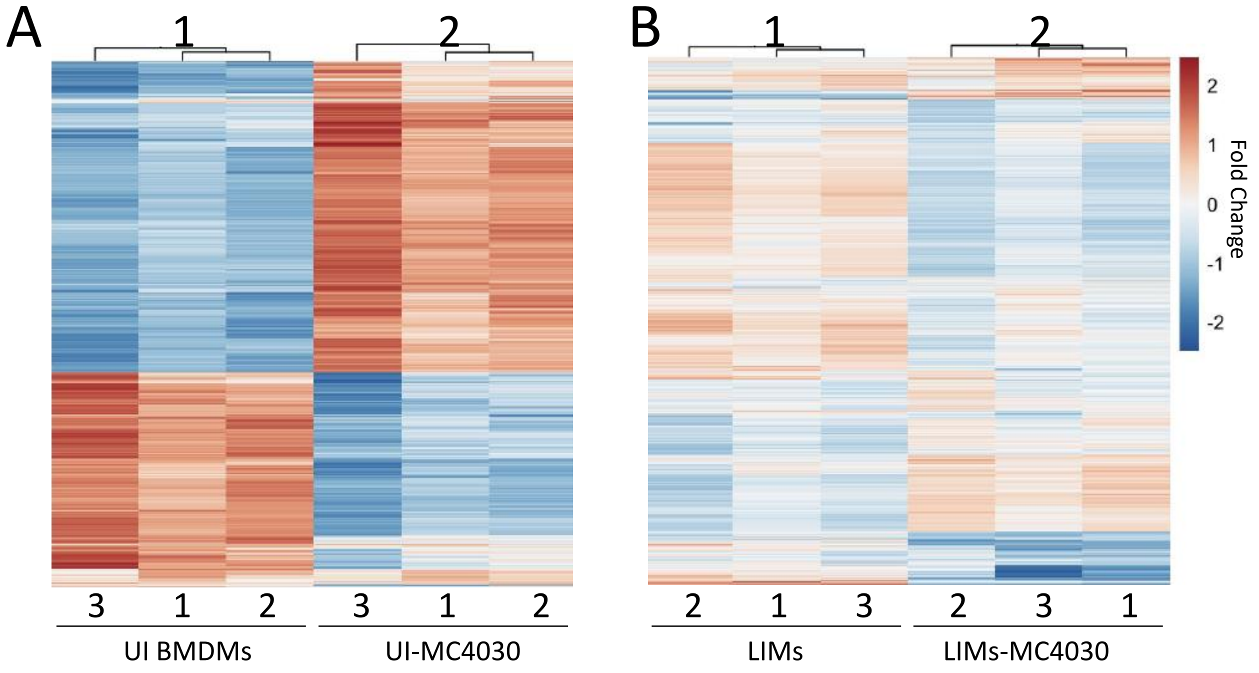


**Figure S1. Gutiérrez-Sanchez M.** ***et al.***

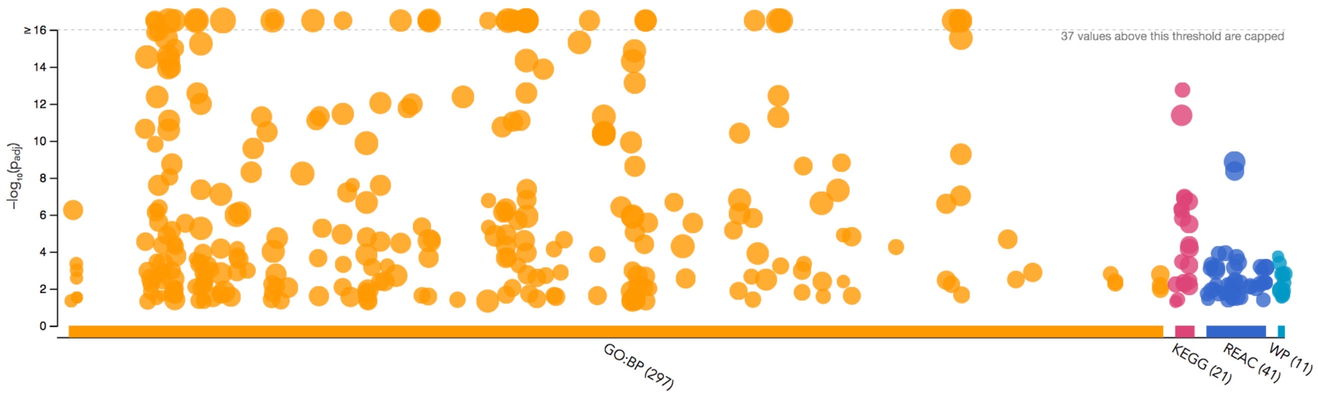


GO:BP — stats

| Term name | Term ID | $p_{adj}$ |
|---|---|---|
| ribonucleoprotein complex biogenesis | GO:0022613 | $8.507\times10^{-30}$ |
| ribosome biogenesis | GO:0042254 | $3.186\times10^{-28}$ |
| ncRNA metabolic process | GO:0034660 | $9.006\times10^{-24}$ |
| rRNA processing | GO:0006364 | $2.554\times10^{-22}$ |
| ncRNA processing | GO:0034470 | $1.169\times10^{-21}$ |
| rRNA metabolic process | GO:0016072 | $4.954\times10^{-21}$ |
| peptide biosynthetic process | GO:0043043 | $7.272\times10^{-17}$ |
| translation | GO:0006412 | $1.334\times10^{-16}$ |
| mitochondrion organization | GO:0007005 | $9.999\times10^{-16}$ |
| tRNA metabolic process | GO:0006399 | $1.539\times10^{-10}$ |
| organophosphate biosynthetic process | GO:0090407 | $1.557\times10^{-9}$ |
| establishment of protein localization to organelle | GO:0072594 | $2.336\times10^{-9}$ |
| mitochondrial transport | GO:0006839 | $9.090\times10^{-9}$ |
| maturation of SSU-rRNA | GO:0030490 | $2.510\times10^{-8}$ |
| ribosomal large subunit biogenesis | GO:0042273 | $1.714\times10^{-7}$ |
| nucleobase-containing small molecule metabolic process | GO:0055086 | $2.131\times10^{-7}$ |
| cellular amino acid metabolic process | GO:0006520 | $4.377\times10^{-7}$ |
| tRNA aminoacylation | GO:0043039 | $5.305\times10^{-7}$ |
| tRNA aminoacylation for protein translation | GO:0006418 | $5.778\times10^{-7}$ |
| regulation of translation | GO:0006417 | $7.362\times10^{-7}$ |
| amino acid activation | GO:0043038 | $1.031\times10^{-6}$ |
| regulation of mitotic cell cycle | GO:0007346 | $2.907\times10^{-6}$ |
| regulation of cellular amide metabolic process | GO:0034248 | $4.452\times10^{-6}$ |
| ribosomal small subunit biogenesis | GO:0042274 | $4.828\times10^{-6}$ |
| process utilizing autophagic mechanism | GO:0061919 | $6.950\times10^{-6}$ |
| autophagy | GO:0006914 | $6.950\times10^{-6}$ |
| RNA phosphodiester bond hydrolysis | GO:0090501 | $1.255\times10^{-5}$ |
| nucleoside phosphate metabolic process | GO:0006753 | $1.284\times10^{-5}$ |
| response to endoplasmic reticulum stress | GO:0034976 | $2.220\times10^{-5}$ |
| ATP metabolic process | GO:0046034 | $2.274\times10^{-5}$ |
| generation of precursor metabolites and energy | GO:0006091 | $2.855\times10^{-5}$ |
| mitochondrial membrane organization | GO:0007006 | $3.703\times10^{-5}$ |
| protein targeting | GO:0006605 | $4.381\times10^{-5}$ |
| vacuole organization | GO:0007033 | $5.154\times10^{-5}$ |
| mitochondrial gene expression | GO:0140053 | $5.600\times10^{-5}$ |
| negative regulation of autophagy | GO:0010507 | $6.975\times10^{-5}$ |
| positive regulation of translation | GO:0045727 | $7.315\times10^{-5}$ |
| nucleotide metabolic process | GO:0009117 | $1.205\times10^{-4}$ |
| organelle fusion | GO:0048284 | $1.420\times10^{-4}$ |
| regulation of autophagy | GO:0010506 | $1.856\times10^{-4}$ |
| proteasome-mediated ubiquitin-dependent protein cata... | GO:0043161 | $1.995\times10^{-4}$ |
| ribose phosphate metabolic process | GO:0019693 | $2.228\times10^{-4}$ |
| mitochondrial fusion | GO:0008053 | $2.387\times10^{-4}$ |
| protein folding | GO:0006457 | $3.585\times10^{-4}$ |
| phospholipid biosynthetic process | GO:0008654 | $3.899\times10^{-4}$ |
| maturation of SSU-rRNA from tricistronic rRNA transcri... | GO:0000462 | $4.486\times10^{-4}$ |
| establishment of protein localization to mitochondrion | GO:0072655 | $4.584\times10^{-4}$ |
| ribonucleoprotein complex assembly | GO:0022618 | $4.862\times10^{-4}$ |
| dicarboxylic acid metabolic process | GO:0043648 | $5.499\times10^{-4}$ |
| ribonucleotide metabolic process | GO:0009259 | $5.542\times10^{-4}$ |
| ribonucleoprotein complex subunit organization | GO:0071826 | $5.892\times10^{-4}$ |
| mitochondrial translation | GO:0032543 | $6.026\times10^{-4}$ |
| regulation of protein stability | GO:0031647 | $6.869\times10^{-4}$ |
| RNA modification | GO:0009451 | $9.777\times10^{-4}$ |
| maturation of LSU-rRNA | GO:0000470 | $1.007\times10^{-3}$ |
| regulation of mitochondrion organization | GO:0010821 | $1.042\times10^{-3}$ |
| nuclear transport | GO:0051169 | $1.045\times10^{-3}$ |
| nucleocytoplasmic transport | GO:0006913 | $1.045\times10^{-3}$ |
| purine-containing compound metabolic process | GO:0072521 | $1.056\times10^{-3}$ |
| purine nucleotide metabolic process | GO:0006163 | $1.104\times10^{-3}$ |
| protein targeting to mitochondrion | GO:0006626 | $1.134\times10^{-3}$ |
| proteasomal protein catabolic process | GO:0010498 | $1.221\times10^{-3}$ |
| positive regulation of mitotic cell cycle | GO:0045931 | $1.331\times10^{-3}$ |
| serine family amino acid metabolic process | GO:0009069 | $1.426\times10^{-3}$ |
| mitochondrial transmembrane transport | GO:1990542 | $1.599\times10^{-3}$ |
| regulation of apoptotic signaling pathway | GO:2001233 | $1.709\times10^{-3}$ |
| ribonucleoside triphosphate metabolic process | GO:0009199 | $1.714\times10^{-3}$ |
| cellular respiration | GO:0045333 | $1.965\times10^{-3}$ |
| organelle localization | GO:0051640 | $2.140\times10^{-3}$ |
| protein localization to mitochondrion | GO:0070585 | $2.226\times10^{-3}$ |
| translational initiation | GO:0006413 | $2.426\times10^{-3}$ |
| mitotic cell cycle phase transition | GO:0044772 | $2.582\times10^{-3}$ |
| cleavage involved in rRNA processing | GO:0000469 | $2.658\times10^{-3}$ |
| regulation of protein modification by small protein conju... | GO:1903320 | $3.180\times10^{-3}$ |
| carbohydrate derivative biosynthetic process | GO:1901137 | $3.479\times10^{-3}$ |
| phospholipid metabolic process | GO:0006644 | $3.564\times10^{-3}$ |
| purine ribonucleotide metabolic process | GO:0009150 | $3.705\times10^{-3}$ |
| RNA phosphodiester bond hydrolysis, endonucleolytic | GO:0090502 | $4.127\times10^{-3}$ |
| cellular response to leukemia inhibitory factor | GO:1990830 | $4.166\times10^{-3}$ |
| methylation | GO:0032259 | $4.248\times10^{-3}$ |
| nucleoside triphosphate metabolic process | GO:0009141 | $5.113\times10^{-3}$ |
| response to leukemia inhibitory factor | GO:1990823 | $5.177\times10^{-3}$ |
| DNA replication | GO:0006260 | $5.518\times10^{-3}$ |
| viral process | GO:0016032 | $5.666\times10^{-3}$ |
| nucleoside phosphate biosynthetic process | GO:1901293 | $5.796\times10^{-3}$ |
| nucleotide biosynthetic process | GO:0009165 | $6.074\times10^{-3}$ |
| positive regulation of hydrolase activity | GO:0051345 | $6.781\times10^{-3}$ |
| negative regulation of apoptotic signaling pathway | GO:2001234 | $6.997\times10^{-3}$ |
| macroautophagy | GO:0016236 | $7.598\times10^{-3}$ |
| tRNA processing | GO:0008033 | $7.752\times10^{-3}$ |
| protein maturation | GO:0051604 | $8.181\times10^{-3}$ |
| regulation of cell morphogenesis | GO:0022604 | $8.209\times10^{-3}$ |
| inner mitochondrial membrane organization | GO:0007007 | $9.020\times10^{-3}$ |
| macromolecule methylation | GO:0043414 | $9.134\times10^{-3}$ |
| apoptotic mitochondrial changes | GO:0008637 | $9.427\times10^{-3}$ |
| modulation of process of other organism involved in sy... | GO:0051817 | $9.696\times10^{-3}$ |
| vacuolar transport | GO:0007034 | $9.718\times10^{-3}$ |
| regulation of extrinsic apoptotic signaling pathway | GO:2001236 | $1.113\times10^{-2}$ |
| cellular response to chemical stress | GO:0062197 | $1.337\times10^{-2}$ |
| response to oxidative stress | GO:0006979 | $1.407\times10^{-2}$ |

KEGG — stats

| Term name | Term ID | $p_{adj}$ |
|---|---|---|
| Ribosome biogenesis in eukaryotes | KEGG:03008 | $1.783\times10^{-13}$ |
| Protein processing in endoplasmic reticulum | KEGG:04141 | $1.155\times10^{-7}$ |
| Lysosome | KEGG:04142 | $1.308\times10^{-7}$ |
| Parkinson disease | KEGG:05012 | $1.947\times10^{-7}$ |
| Aminoacyl-tRNA biosynthesis | KEGG:00970 | $4.781\times10^{-7}$ |
| Carbon metabolism | KEGG:01200 | $5.704\times10^{-7}$ |
| RNA transport | KEGG:03013 | $1.544\times10^{-6}$ |
| Amyotrophic lateral sclerosis | KEGG:05014 | $3.253\times10^{-6}$ |
| Prion disease | KEGG:05020 | $4.297\times10^{-5}$ |
| Pathways of neurodegeneration - multiple diseases | KEGG:05022 | $6.607\times10^{-5}$ |
| Biosynthesis of amino acids | KEGG:01230 | $3.464\times10^{-4}$ |
| Alzheimer disease | KEGG:05010 | $5.392\times10^{-4}$ |
| Spinocerebellar ataxia | KEGG:05017 | $3.933\times10^{-3}$ |
| Cell cycle | KEGG:04110 | $4.333\times10^{-3}$ |
| Proteasome | KEGG:03050 | $5.793\times10^{-3}$ |
| Oxidative phosphorylation | KEGG:00190 | $5.820\times10^{-3}$ |
| Ferroptosis | KEGG:04216 | $6.801\times10^{-3}$ |
| Huntington disease | KEGG:05016 | $7.411\times10^{-3}$ |
| Amino sugar and nucleotide sugar metabolism | KEGG:00520 | $3.860\times10^{-2}$ |
| Steroid biosynthesis | KEGG:00100 | $4.995\times10^{-2}$ |

1 to 20 of 20 Page 1 of 1

REAC — stats

| Term name | Term ID | $p_{adj}$ |
|---|---|---|
| G1/S Transition | REAC:R-MMU-... | $1.218\times10^{-4}$ |
| DNA Replication Pre-Initiation | REAC:R-MMU-... | $1.264\times10^{-4}$ |
| Mitotic G1 phase and G1/S transition | REAC:R-MMU-... | $1.991\times10^{-4}$ |
| Neddylation | REAC:R-MMU-... | $3.853\times10^{-4}$ |
| Assembly of the pre-replicative complex | REAC:R-MMU-... | $4.911\times10^{-4}$ |
| The role of GTSE1 in G2/M progression after G2 checkp... | REAC:R-MMU-... | $6.511\times10^{-4}$ |
| rRNA processing in the nucleus and cytosol | REAC:R-MMU-... | $7.109\times10^{-4}$ |
| rRNA processing | REAC:R-MMU-... | $7.109\times10^{-4}$ |
| Major pathway of rRNA processing in the nucleolus and ... | REAC:R-MMU-... | $7.109\times10^{-4}$ |
| Cellular responses to external stimuli | REAC:R-MMU-... | $7.250\times10^{-4}$ |
| Amino acids regulate mTORC1 | REAC:R-MMU-... | $8.182\times10^{-4}$ |
| Transferrin endocytosis and recycling | REAC:R-MMU-... | $9.611\times10^{-4}$ |
| Cellular responses to stress | REAC:R-MMU-... | $1.005\times10^{-3}$ |
| Metabolism of polyamines | REAC:R-MMU-... | $3.178\times10^{-3}$ |
| Orc1 removal from chromatin | REAC:R-MMU-... | $4.517\times10^{-3}$ |
| p53-Independent G1/S DNA damage checkpoint | REAC:R-MMU-... | $4.710\times10^{-3}$ |
| p53-Independent DNA Damage Response | REAC:R-MMU-... | $4.710\times10^{-3}$ |
| Ubiquitin Mediated Degradation of Phosphorylated Cdc... | REAC:R-MMU-... | $4.710\times10^{-3}$ |
| Metabolism of amino acids and derivatives | REAC:R-MMU-... | $5.113\times10^{-3}$ |
| Cholesterol biosynthesis | REAC:R-MMU-... | $5.504\times10^{-3}$ |
| Insulin receptor recycling | REAC:R-MMU-... | $5.565\times10^{-3}$ |
| Glyoxylate metabolism and glycine degradation | REAC:R-MMU-... | $5.565\times10^{-3}$ |
| The citric acid (TCA) cycle and respiratory electron tran... | REAC:R-MMU-... | $5.569\times10^{-3}$ |
| Regulation of mRNA stability by proteins that bind AU-ri... | REAC:R-MMU-... | $7.187\times10^{-3}$ |
| Regulation of ornithine decarboxylase (ODC) | REAC:R-MMU-... | $8.263\times10^{-3}$ |
| Switching of origins to a post-replicative state | REAC:R-MMU-... | $9.608\times10^{-3}$ |
| CDT1 association with the CDC6:ORC:origin complex | REAC:R-MMU-... | $1.091\times10^{-2}$ |
| Autophagy | REAC:R-MMU-... | $1.206\times10^{-2}$ |
| Macroautophagy | REAC:R-MMU-... | $1.206\times10^{-2}$ |
| G1/S DNA Damage Checkpoints | REAC:R-MMU-... | $1.754\times10^{-2}$ |
| DNA Replication | REAC:R-MMU-... | $1.758\times10^{-2}$ |
| Glycolysis | REAC:R-MMU-... | $1.878\times10^{-2}$ |
| AUF1 (hnRNP D0) binds and destabilizes mRNA | REAC:R-MMU-... | $1.878\times10^{-2}$ |
| Oxygen-dependent proline hydroxylation of Hypoxia-in... | REAC:R-MMU-... | $2.953\times10^{-2}$ |
| Iron uptake and transport | REAC:R-MMU-... | $3.226\times10^{-2}$ |
| Activation of ATR in response to replication stress | REAC:R-MMU-... | $3.608\times10^{-2}$ |
| Hedgehog 'off' state | REAC:R-MMU-... | $3.874\times10^{-2}$ |
| Ubiquitin-dependent degradation of Cyclin D | REAC:R-MMU-... | $4.029\times10^{-2}$ |

1 to 38 of 38 Page 1 of 1

WP — stats

| Term name | Term ID | $p_{adj}$ |
|---|---|---|
| Cholesterol Biosynthesis | WP:WP103 | $1.825\times10^{-4}$ |
| Glycolysis and Gluconeogenesis | WP:WP157 | $4.225\times10^{-4}$ |
| Amino Acid metabolism | WP:WP662 | $1.587\times10^{-3}$ |
| Cholesterol metabolism (includes both Bloch and Kand... | WP:WP4346 | $1.639\times10^{-3}$ |
| Translation Factors | WP:WP307 | $1.639\times10^{-3}$ |
| Proteasome Degradation | WP:WP519 | $4.746\times10^{-3}$ |
| TNF-alpha NF-kB Signaling Pathway | WP:WP246 | $1.126\times10^{-2}$ |
| mRNA processing | WP:WP310 | $1.304\times10^{-2}$ |
| TCA Cycle | WP:WP434 | $1.882\times10^{-2}$ |
| G1 to S cell cycle control | WP:WP413 | $2.258\times10^{-2}$ |
| Fatty Acid Biosynthesis | WP:WP336 | $2.851\times10^{-2}$ |

1 to 11 of 11 Page 1 of 1

**Figure S2. Gutiérrez-Sanchez M. *et al.***

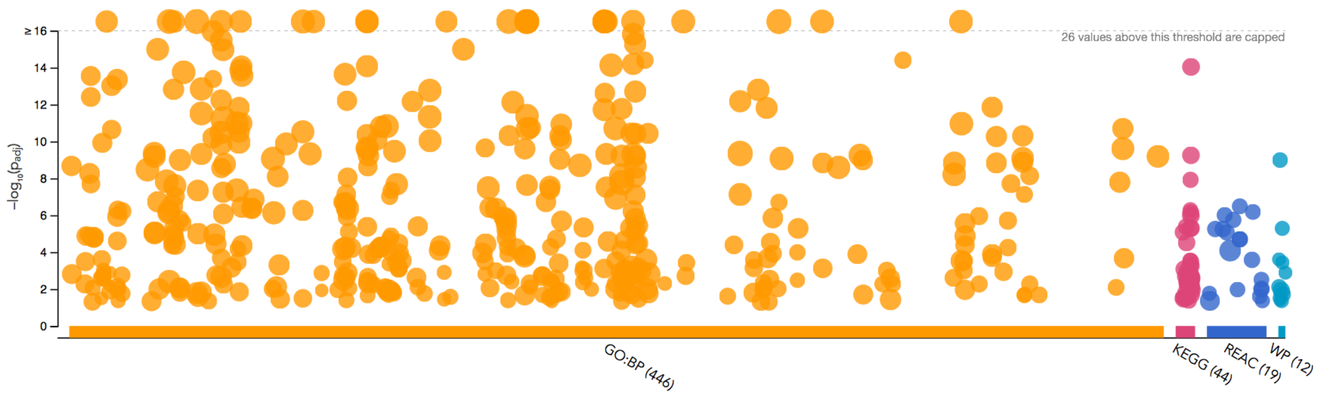


**GO:BP**

| Term name | Term ID | $p_{adj}$ |
|---|---|---|
| defense response to symbiont | GO:0140546 | $3.969\times10^{-15}$ |
| defense response to virus | GO:0051607 | $3.969\times10^{-15}$ |
| response to virus | GO:0009615 | $4.099\times10^{-14}$ |
| T cell activation | GO:0042110 | $2.266\times10^{-10}$ |
| regulation of hemopoiesis | GO:1903706 | $7.277\times10^{-9}$ |
| mononuclear cell differentiation | GO:1903131 | $1.944\times10^{-8}$ |
| positive regulation of cytokine production | GO:0001819 | $2.006\times10^{-8}$ |
| regulation of tumor necrosis factor production | GO:0032680 | $4.464\times10^{-8}$ |
| regulation of tumor necrosis factor superfamily cytokin... | GO:1903555 | $7.435\times10^{-8}$ |
| actin filament organization | GO:0007015 | $9.491\times10^{-8}$ |
| tumor necrosis factor production | GO:0032640 | $9.555\times10^{-8}$ |
| regulation of cell-cell adhesion | GO:0022407 | $1.898\times10^{-7}$ |
| tumor necrosis factor superfamily cytokine production | GO:0071706 | $1.996\times10^{-7}$ |
| lymphocyte differentiation | GO:0030098 | $2.468\times10^{-7}$ |
| regulation of immune effector process | GO:0002697 | $5.131\times10^{-7}$ |
| regulation of response to biotic stimulus | GO:0002831 | $6.018\times10^{-7}$ |
| regulation of leukocyte differentiation | GO:1902105 | $1.105\times10^{-6}$ |
| myeloid cell differentiation | GO:0030099 | $1.206\times10^{-6}$ |
| defense response to protozoan | GO:0042832 | $1.522\times10^{-6}$ |
| leukocyte cell-cell adhesion | GO:0007159 | $1.700\times10^{-6}$ |
| regulation of leukocyte cell-cell adhesion | GO:1903037 | $1.943\times10^{-6}$ |
| I-kappaB kinase/NF-kappaB signaling | GO:0007249 | $2.614\times10^{-6}$ |
| T cell differentiation | GO:0030217 | $7.908\times10^{-6}$ |
| response to protozoan | GO:0001562 | $1.107\times10^{-5}$ |
| regulation of interleukin-6 production | GO:0032675 | $1.305\times10^{-5}$ |
| negative regulation of cytokine production | GO:0001818 | $1.579\times10^{-5}$ |
| positive regulation of cell adhesion | GO:0045785 | $1.934\times10^{-5}$ |
| interleukin-6 production | GO:0032635 | $1.941\times10^{-5}$ |
| regulation of T cell activation | GO:0050863 | $2.170\times10^{-5}$ |
| negative regulation of immune system process | GO:0002683 | $2.469\times10^{-5}$ |
| cellular response to organonitrogen compound | GO:0071417 | $2.940\times10^{-5}$ |
| regulation of actin cytoskeleton organization | GO:0032956 | $3.242\times10^{-5}$ |
| process utilizing autophagic mechanism | GO:0061919 | $4.033\times10^{-5}$ |
| autophagy | GO:0006914 | $4.033\times10^{-5}$ |
| regulation of lymphocyte differentiation | GO:0045619 | $5.194\times10^{-5}$ |
| regulation of I-kappaB kinase/NF-kappaB signaling | GO:0043122 | $5.194\times10^{-5}$ |
| positive regulation of leukocyte cell-cell adhesion | GO:1903039 | $5.664\times10^{-5}$ |
| positive regulation of autophagy | GO:0010508 | $6.632\times10^{-5}$ |
| positive regulation of response to external stimulus | GO:0032103 | $6.720\times10^{-5}$ |
| alpha-beta T cell activation | GO:0046631 | $7.848\times10^{-5}$ |
| signal transduction by p53 class mediator | GO:0072331 | $1.027\times10^{-4}$ |
| regulation of actin filament-based process | GO:0032970 | $1.106\times10^{-4}$ |
| cellular response to external stimulus | GO:0071496 | $1.193\times10^{-4}$ |
| regulation of interleukin-12 production | GO:0032655 | $1.236\times10^{-4}$ |
| negative regulation of protein modification process | GO:0031400 | $1.290\times10^{-4}$ |
| positive regulation of I-kappaB kinase/NF-kappaB signa... | GO:0043123 | $1.311\times10^{-4}$ |
| cellular response to extracellular stimulus | GO:0031668 | $1.454\times10^{-4}$ |
| negative regulation of intracellular signal transduction | GO:1902532 | $1.964\times10^{-4}$ |
| positive regulation of catabolic process | GO:0009896 | $2.088\times10^{-4}$ |
| response to molecule of bacterial origin | GO:0002237 | $2.369\times10^{-4}$ |
| regulation of leukocyte proliferation | GO:0070663 | $2.633\times10^{-4}$ |
| response to peptide | GO:1901652 | $2.980\times10^{-4}$ |
| multicellular organismal homeostasis | GO:0048871 | $2.990\times10^{-4}$ |
| cellular response to hormone stimulus | GO:0032870 | $3.123\times10^{-4}$ |
| T cell proliferation | GO:0042098 | $3.153\times10^{-4}$ |
| angiogenesis | GO:0001525 | $3.237\times10^{-4}$ |
| regulation of autophagy | GO:0010506 | $3.364\times10^{-4}$ |
| positive regulation of cell-cell adhesion | GO:0022409 | $3.364\times10^{-4}$ |
| bone development | GO:0060348 | $3.734\times10^{-4}$ |
| interleukin-12 production | GO:0032615 | $4.390\times10^{-4}$ |
| positive regulation of hydrolase activity | GO:0051345 | $6.541\times10^{-4}$ |
| regulation of viral entry into host cell | GO:0046596 | $6.830\times10^{-4}$ |
| mononuclear cell proliferation | GO:0032943 | $7.047\times10^{-4}$ |
| leukocyte proliferation | GO:0070661 | $7.147\times10^{-4}$ |
| positive regulation of T cell activation | GO:0050870 | $7.382\times10^{-4}$ |
| leukocyte migration | GO:0050900 | $7.996\times10^{-4}$ |
| regulation of mononuclear cell proliferation | GO:0032944 | $8.150\times10^{-4}$ |
| alpha-beta T cell activation involved in immune response | GO:0002287 | $9.628\times10^{-4}$ |
| cellular response to peptide | GO:1901653 | $9.678\times10^{-4}$ |
| regulation of actin filament organization | GO:0110053 | $1.006\times10^{-3}$ |
| response to starvation | GO:0042594 | $1.016\times10^{-3}$ |
| lymphocyte proliferation | GO:0046651 | $1.030\times10^{-3}$ |
| regulation of cytoskeleton organization | GO:0051493 | $1.038\times10^{-3}$ |
| regulation of supramolecular fiber organization | GO:1902903 | $1.152\times10^{-3}$ |
| regulation of lymphocyte proliferation | GO:0050670 | $1.231\times10^{-3}$ |
| response to interferon-beta | GO:0035456 | $1.247\times10^{-3}$ |
| antigen processing and presentation of exogenous pept... | GO:0002478 | $1.250\times10^{-3}$ |
| antigen processing and presentation | GO:0019882 | $1.352\times10^{-3}$ |
| T cell differentiation involved in immune response | GO:0002292 | $1.532\times10^{-3}$ |
| T-helper cell differentiation | GO:0042093 | $1.538\times10^{-3}$ |
| positive regulation of DNA-binding transcription factor ... | GO:0051091 | $1.624\times10^{-3}$ |
| regulation of DNA-binding transcription factor activity | GO:0051090 | $1.696\times10^{-3}$ |
| regulation of innate immune response | GO:0045088 | $1.739\times10^{-3}$ |
| regulation of adaptive immune response | GO:0002819 | $1.739\times10^{-3}$ |
| negative regulation of cell-cell adhesion | GO:0022408 | $1.745\times10^{-3}$ |
| regulation of defense response to virus | GO:0050688 | $1.813\times10^{-3}$ |
| cell chemotaxis | GO:0060326 | $1.929\times10^{-3}$ |
| myeloid cell homeostasis | GO:0002262 | $2.071\times10^{-3}$ |
| CD4-positive, alpha-beta T cell differentiation involved i... | GO:0002294 | $2.124\times10^{-3}$ |
| positive regulation of defense response | GO:0031349 | $2.165\times10^{-3}$ |
| positive regulation of NF-kappaB transcription factor ac... | GO:0051092 | $2.302\times10^{-3}$ |
| viral entry into host cell | GO:0046718 | $2.459\times10^{-3}$ |
| regulation of vasculature development | GO:1901342 | $2.466\times10^{-3}$ |
| pattern recognition receptor signaling pathway | GO:0002221 | $2.498\times10^{-3}$ |
| plasma membrane bounded cell projection assembly | GO:0120031 | $2.557\times10^{-3}$ |
| regulation of angiogenesis | GO:0045765 | $2.788\times10^{-3}$ |
| alpha-beta T cell differentiation involved in immune res... | GO:0002293 | $2.907\times10^{-3}$ |
| cellular response to lipid | GO:0071396 | $2.962\times10^{-3}$ |
| intrinsic apoptotic signaling pathway by p53 class medi... | GO:0072332 | $3.309\times10^{-3}$ |
| regulation of erythrocyte differentiation | GO:0045646 | $3.437\times10^{-3}$ |

**KEGG**

| Term name | Term ID | $p_{adj}$ |
|---|---|---|
| Coronavirus disease - COVID-19 | KEGG:05171 | $9.095\times10^{-15}$ |
| Epstein-Barr virus infection | KEGG:05169 | $5.587\times10^{-10}$ |
| Toxoplasmosis | KEGG:05145 | $1.198\times10^{-8}$ |
| Leishmaniasis | KEGG:05140 | $5.080\times10^{-7}$ |
| Influenza A | KEGG:05164 | $7.373\times10^{-7}$ |
| Kaposi sarcoma-associated herpesvirus infection | KEGG:05167 | $1.074\times10^{-6}$ |
| Osteoclast differentiation | KEGG:04380 | $4.191\times10^{-6}$ |
| Tuberculosis | KEGG:05152 | $4.928\times10^{-6}$ |
| Human T-cell leukemia virus 1 infection | KEGG:05166 | $4.999\times10^{-6}$ |
| Ribosome | KEGG:03010 | $8.285\times10^{-6}$ |
| NOD-like receptor signaling pathway | KEGG:04621 | $3.043\times10^{-5}$ |
| Pertussis | KEGG:05133 | $2.747\times10^{-4}$ |
| Hepatitis B | KEGG:05161 | $3.205\times10^{-4}$ |
| Yersinia infection | KEGG:05135 | $5.512\times10^{-4}$ |
| Human cytomegalovirus infection | KEGG:05163 | $5.545\times10^{-4}$ |
| Chemokine signaling pathway | KEGG:04062 | $8.292\times10^{-4}$ |
| C-type lectin receptor signaling pathway | KEGG:04625 | $1.041\times10^{-3}$ |
| Th1 and Th2 cell differentiation | KEGG:04658 | $1.424\times10^{-3}$ |
| Measles | KEGG:05162 | $1.615\times10^{-3}$ |
| Cellular senescence | KEGG:04218 | $2.006\times10^{-3}$ |
| Viral carcinogenesis | KEGG:05203 | $2.252\times10^{-3}$ |
| Inflammatory bowel disease | KEGG:05321 | $2.312\times10^{-3}$ |
| TNF signaling pathway | KEGG:04668 | $2.365\times10^{-3}$ |
| Herpes simplex virus 1 infection | KEGG:05168 | $2.494\times10^{-3}$ |
| Human papillomavirus infection | KEGG:05165 | $3.097\times10^{-3}$ |
| Antigen processing and presentation | KEGG:04612 | $3.315\times10^{-3}$ |
| Notch signaling pathway | KEGG:04330 | $4.056\times10^{-3}$ |
| Acute myeloid leukemia | KEGG:05221 | $4.456\times10^{-3}$ |
| Th17 cell differentiation | KEGG:04659 | $5.352\times10^{-3}$ |
| Toll-like receptor signaling pathway | KEGG:04620 | $5.407\times10^{-3}$ |
| Allograft rejection | KEGG:05330 | $5.439\times10^{-3}$ |
| Intestinal immune network for IgA production | KEGG:04672 | $5.555\times10^{-3}$ |
| AGE-RAGE signaling pathway in diabetic complications | KEGG:04933 | $6.607\times10^{-3}$ |
| Salmonella infection | KEGG:05132 | $7.932\times10^{-3}$ |
| Viral myocarditis | KEGG:05416 | $7.983\times10^{-3}$ |
| Longevity regulating pathway | KEGG:04211 | $1.515\times10^{-2}$ |
| Human immunodeficiency virus 1 infection | KEGG:05170 | $2.039\times10^{-2}$ |
| EGFR tyrosine kinase inhibitor resistance | KEGG:01521 | $2.834\times10^{-2}$ |
| Circadian rhythm | KEGG:04710 | $2.925\times10^{-2}$ |
| Cell adhesion molecules | KEGG:04514 | $2.967\times10^{-2}$ |
| MAPK signaling pathway | KEGG:04010 | $3.212\times10^{-2}$ |
| p53 signaling pathway | KEGG:04115 | $3.485\times10^{-2}$ |
| Insulin resistance | KEGG:04931 | $4.246\times10^{-2}$ |

1 to 43 of 43 Page 1 of 1

**REAC**

| Term name | Term ID | $p_{adj}$ |
|---|---|---|
| Nonsense Mediated Decay (NMD) independent of the E... | REAC:R-MMU-... | $3.268\times10^{-7}$ |
| SRP-dependent cotranslational protein targeting to me... | REAC:R-MMU-... | $6.556\times10^{-7}$ |
| Formation of a pool of free 40S subunits | REAC:R-MMU-... | $9.666\times10^{-7}$ |
| L13a-mediated translational silencing of Ceruloplasmin ... | REAC:R-MMU-... | $1.777\times10^{-6}$ |
| Eukaryotic Translation Initiation | REAC:R-MMU-... | $5.553\times10^{-6}$ |
| Cap-dependent Translation Initiation | REAC:R-MMU-... | $5.553\times10^{-6}$ |
| GTP hydrolysis and joining of the 60S ribosomal subunit | REAC:R-MMU-... | $8.488\times10^{-6}$ |
| Nonsense-Mediated Decay (NMD) | REAC:R-MMU-... | $1.998\times10^{-5}$ |
| Nonsense Mediated Decay (NMD) enhanced by the Exo... | REAC:R-MMU-... | $1.998\times10^{-5}$ |
| Rho GTPase cycle | REAC:R-MMU-... | $2.592\times10^{-4}$ |
| Toll Like Receptor 9 (TLR9) Cascade | REAC:R-MMU-... | $3.083\times10^{-3}$ |
| Toll-like Receptor Cascades | REAC:R-MMU-... | $9.677\times10^{-3}$ |
| MyD88 dependent cascade initiated on endosome | REAC:R-MMU-... | $1.061\times10^{-2}$ |
| Toll Like Receptor 7/8 (TLR7/8) Cascade | REAC:R-MMU-... | $1.061\times10^{-2}$ |
| Activation of the mRNA upon binding of the cap-bindin... | REAC:R-MMU-... | $1.692\times10^{-2}$ |
| TRAF6 mediated induction of NFkB and MAP kinases up... | REAC:R-MMU-... | $2.482\times10^{-2}$ |
| Translation initiation complex formation | REAC:R-MMU-... | $4.277\times10^{-2}$ |

1 to 17 of 17 Page 1 of 1

**WP**

| Term name | Term ID | $p_{adj}$ |
|---|---|---|
| Cytoplasmic Ribosomal Proteins | WP:WP163 | $1.021\times10^{-9}$ |
| TYROBP Causal Network | WP:WP3625 | $5.130\times10^{-6}$ |
| Type II interferon signaling (IFNG) | WP:WP1253 | $2.448\times10^{-4}$ |
| Microglia Pathogen Phagocytosis Pathway | WP:WP3626 | $3.635\times10^{-4}$ |
| Toll Like Receptor signaling | WP:WP88 | $1.290\times10^{-3}$ |
| Apoptosis | WP:WP1254 | $7.603\times10^{-3}$ |
| TNF-alpha NF-kB Signaling Pathway | WP:WP246 | $1.174\times10^{-2}$ |
| Macrophage markers | WP:WP2271 | $1.677\times10^{-2}$ |
| IL-6 signaling Pathway | WP:WP387 | $1.912\times10^{-2}$ |
| Oxidative Damage | WP:WP1496 | $3.075\times10^{-2}$ |
| Apoptosis Modulation by HSP70 | WP:WP166 | $4.088\times10^{-2}$ |
| Notch Signaling Pathway | WP:WP29 | $4.337\times10^{-2}$ |

1 to 12 of 12 Page 1 of 1

**Figure S3. Gutiérrez-Sanchez M.** ***et al.***

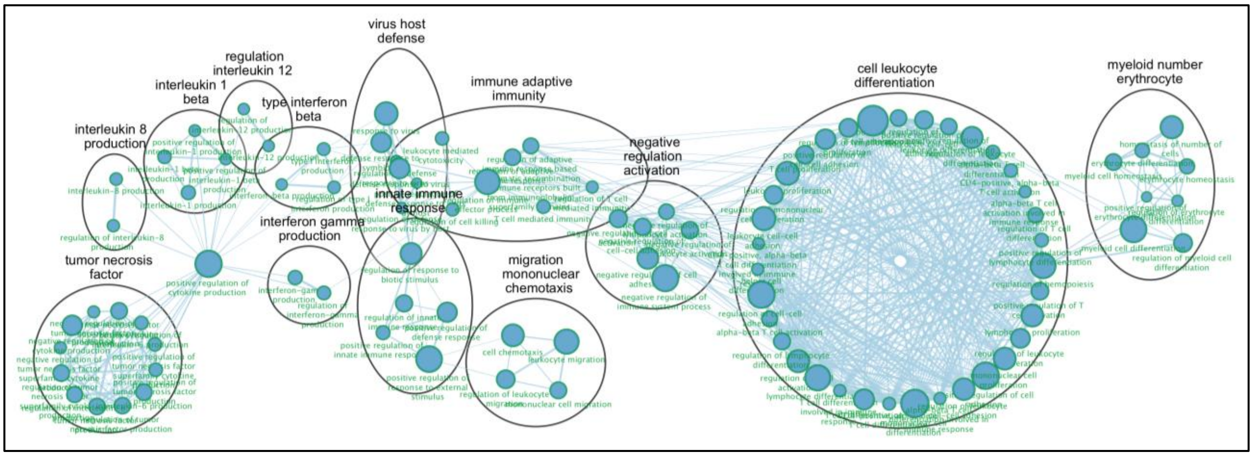


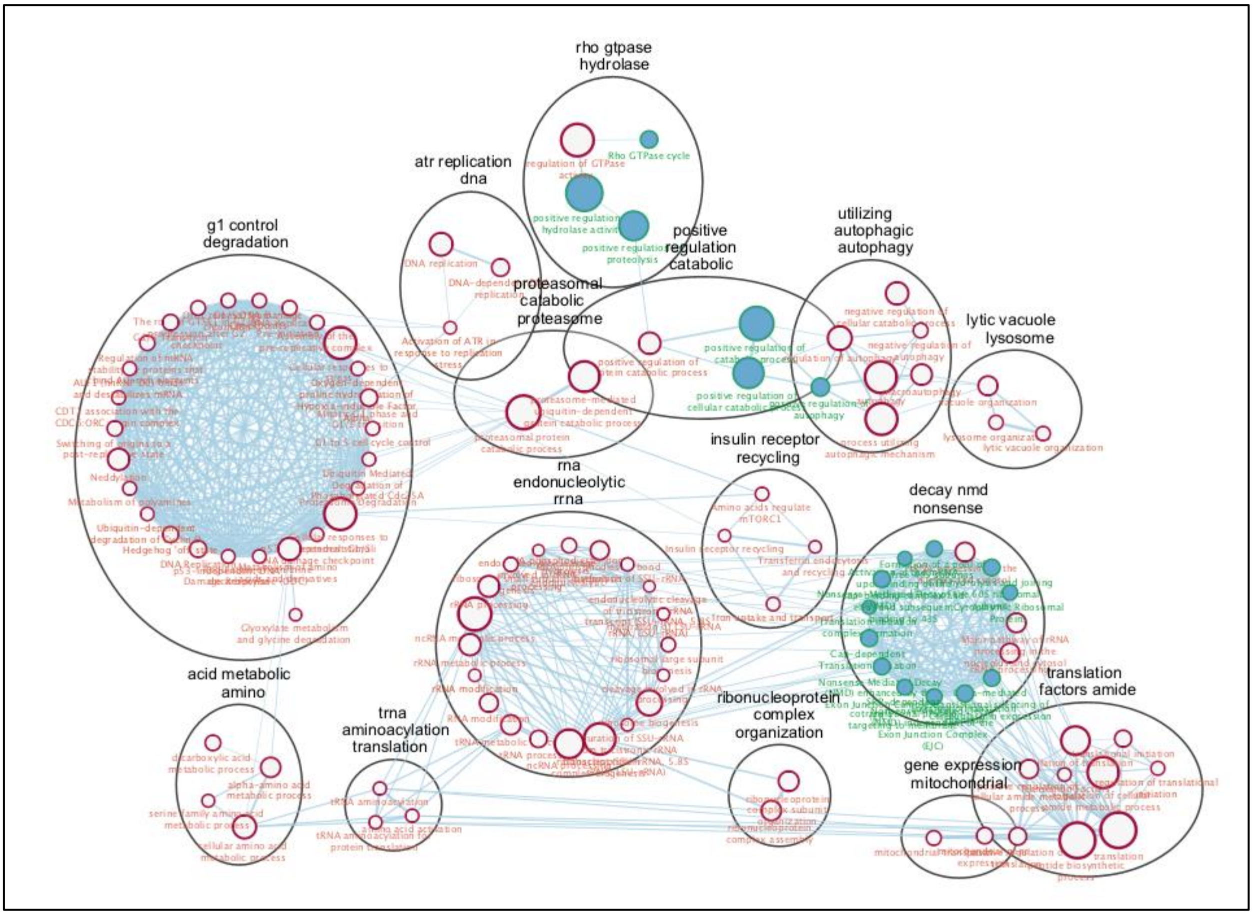


**Figure S4. Gutiérrez-Sanchez M.** ***et al.***

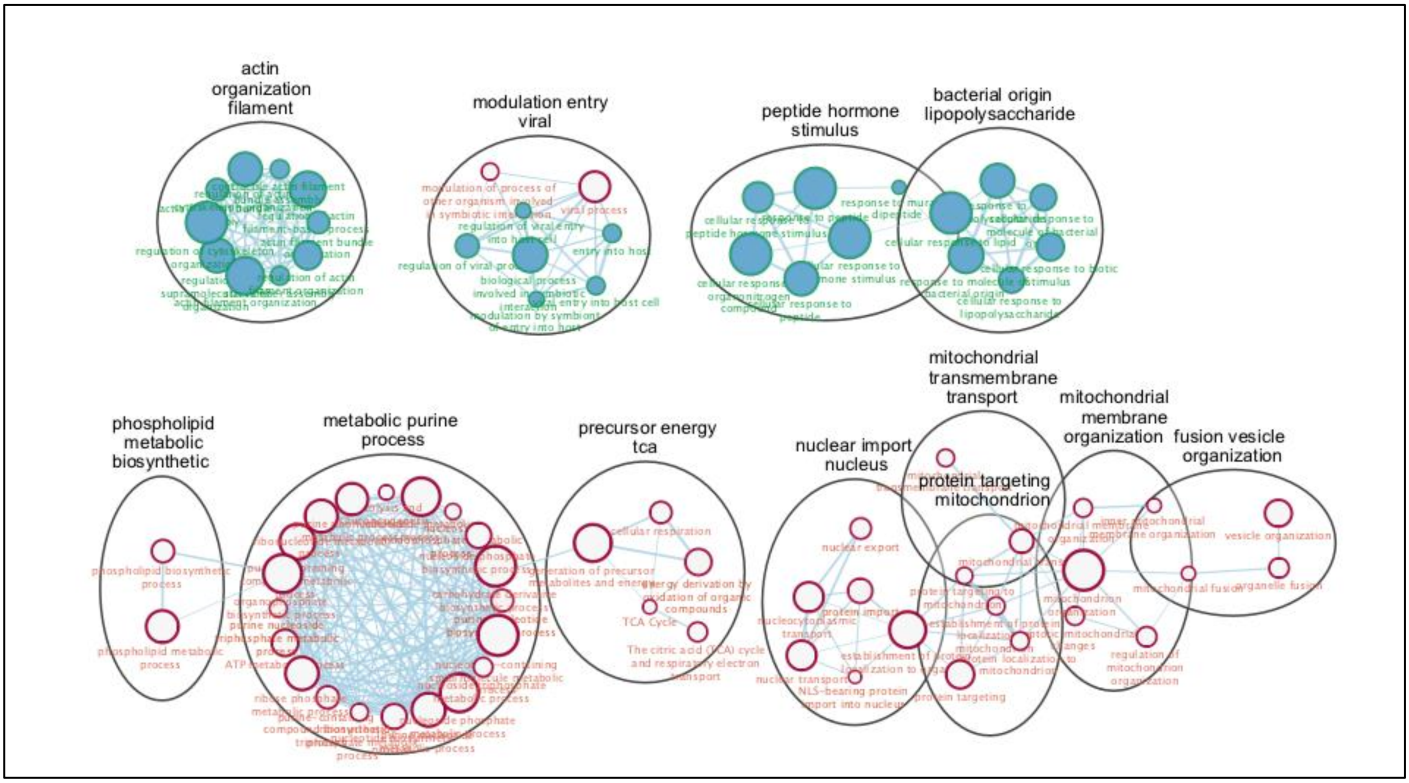


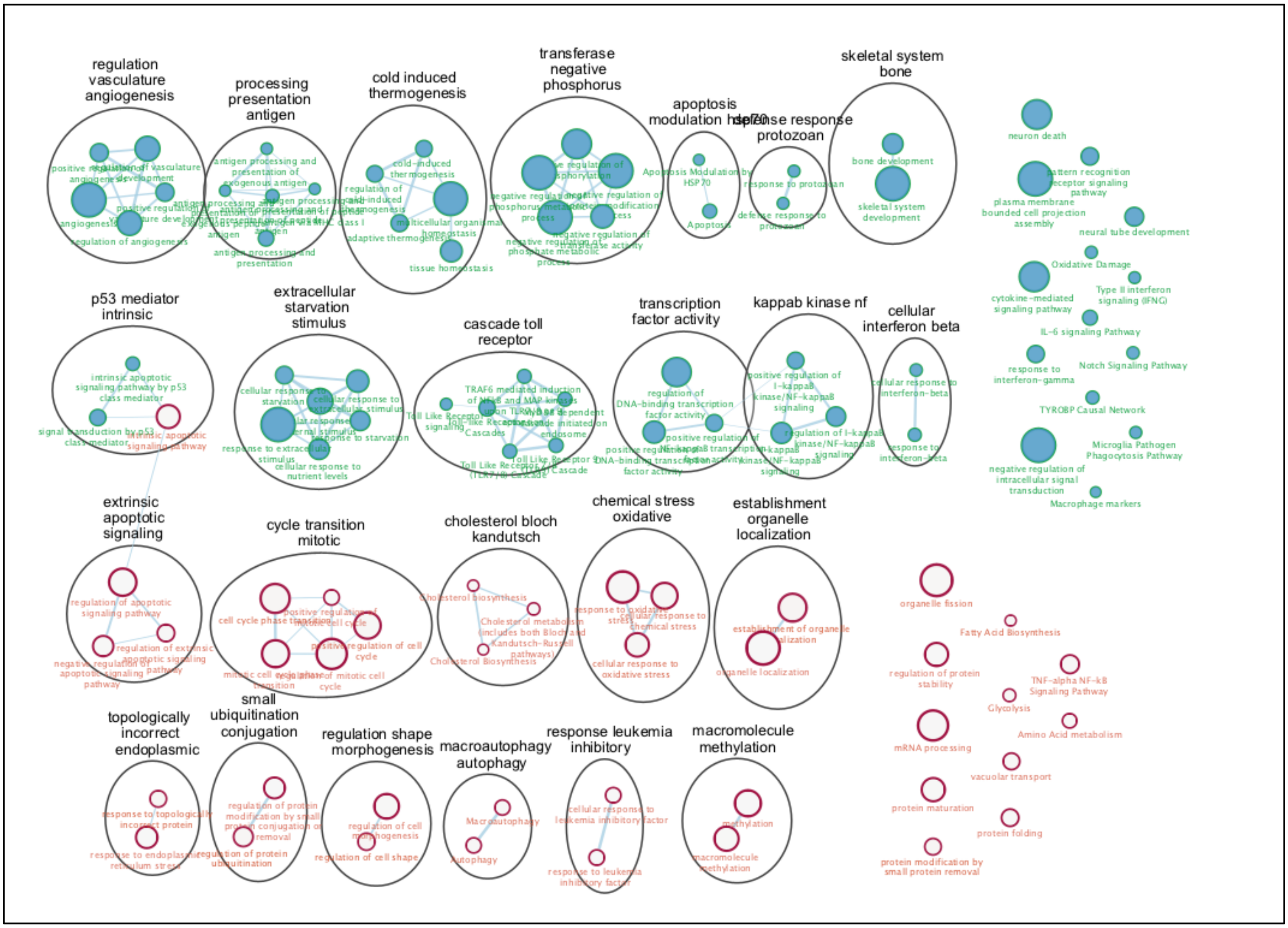


**Figure S5. Gutiérrez-Sanchez M.** ***et al.***

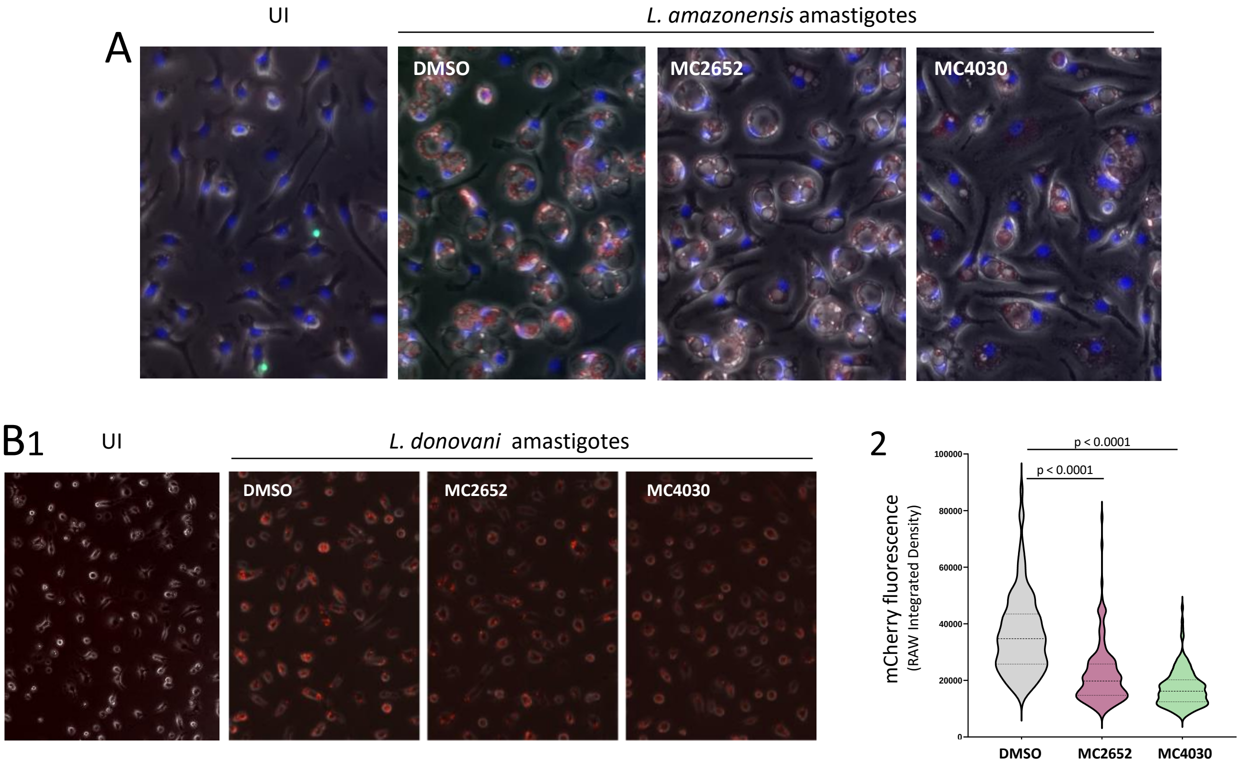


**Figure S6. Gutiérrez-Sanchez M. *et al.***

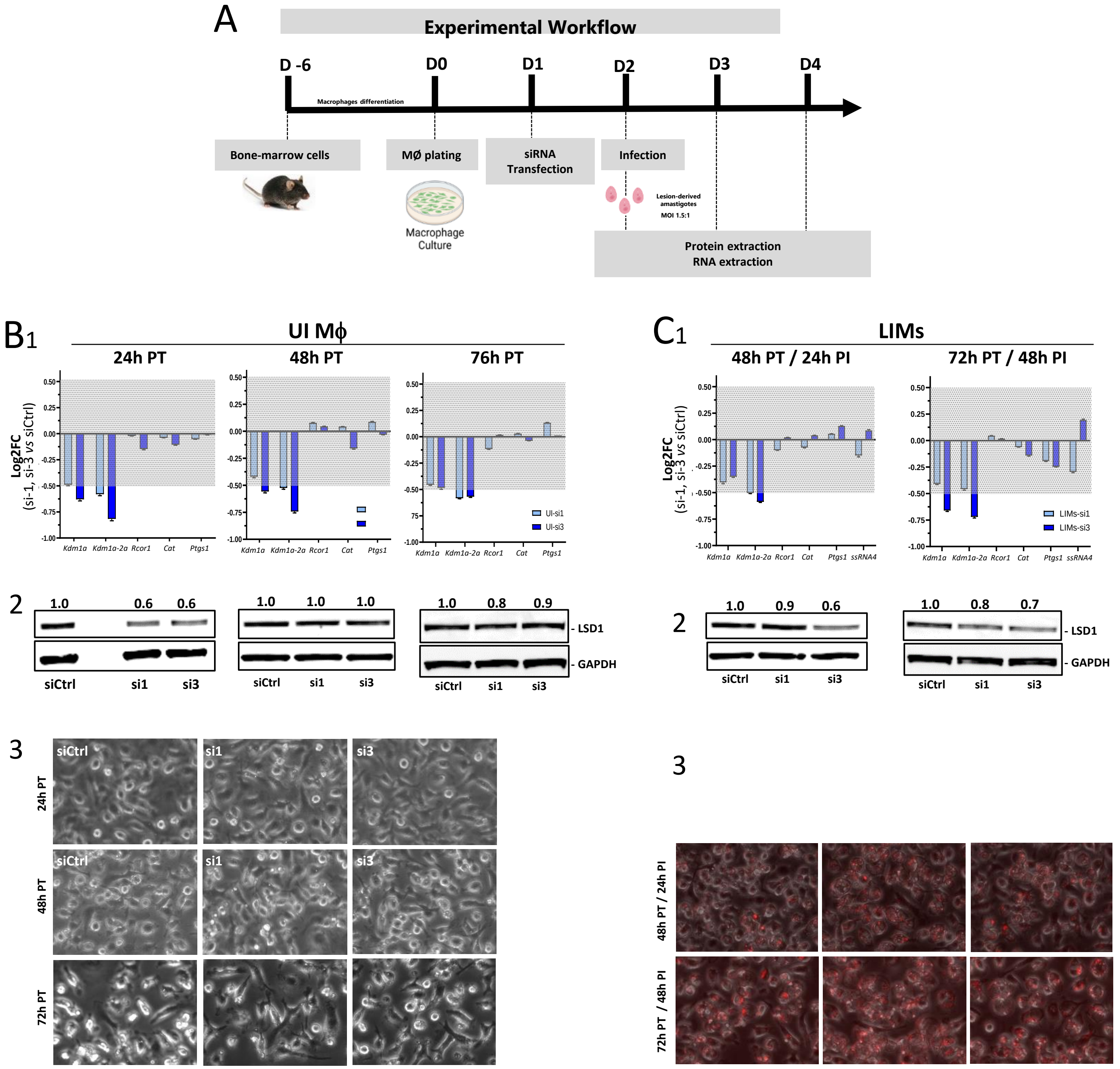


**Figure S7. Gutiérrez-Sanchez M.** ***et al.***

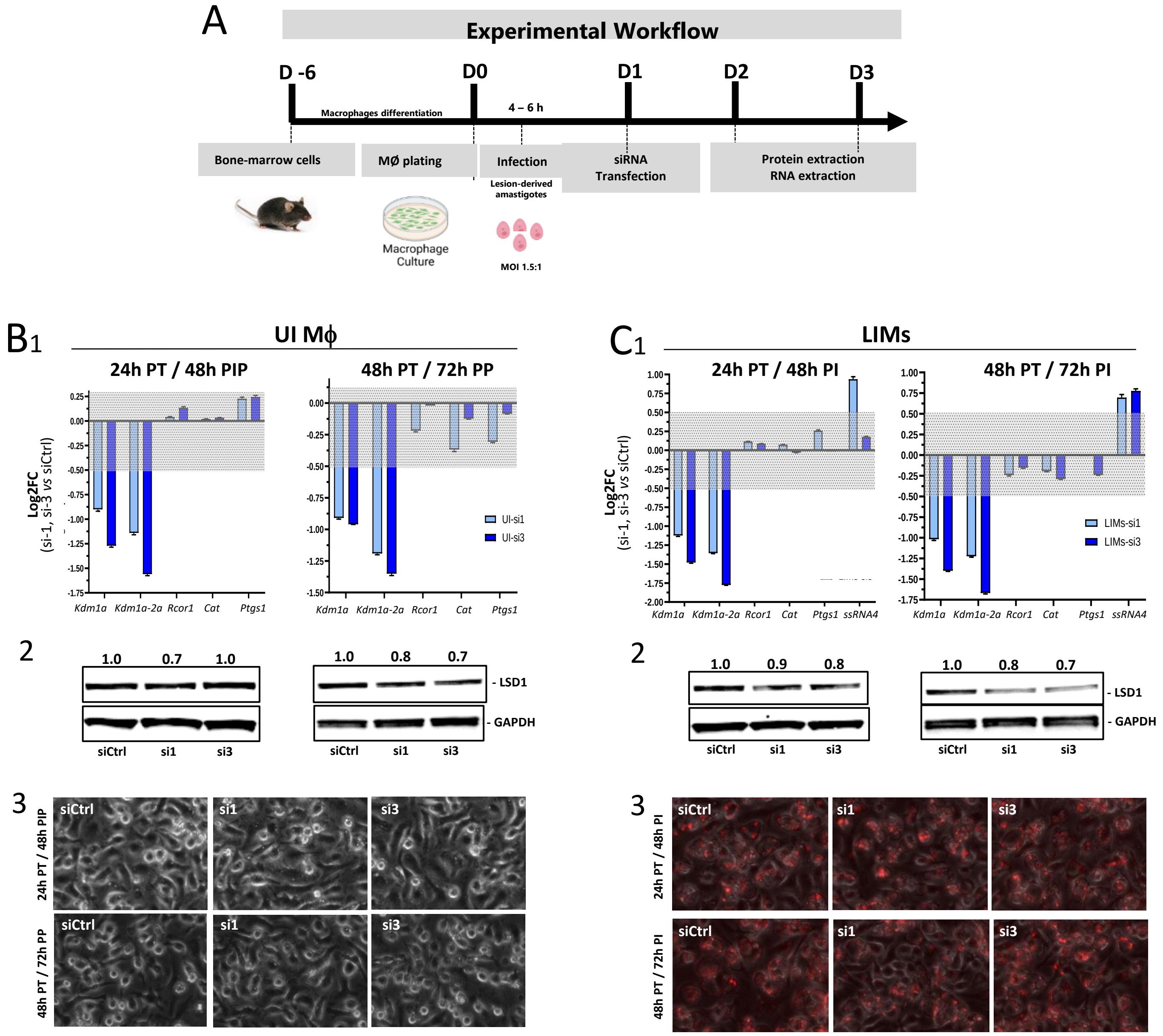


**Figure S8. Gutiérrez-Sanchez M. *et al.***

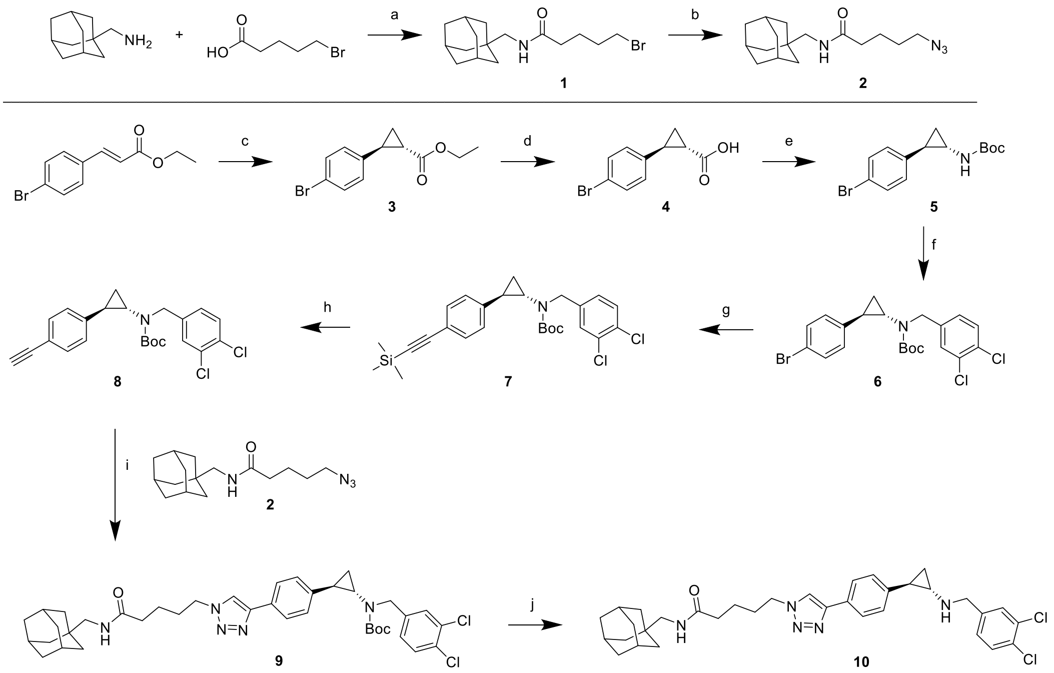


**Figure S9. Gutiérrez-Sanchez M.** ***et al.***

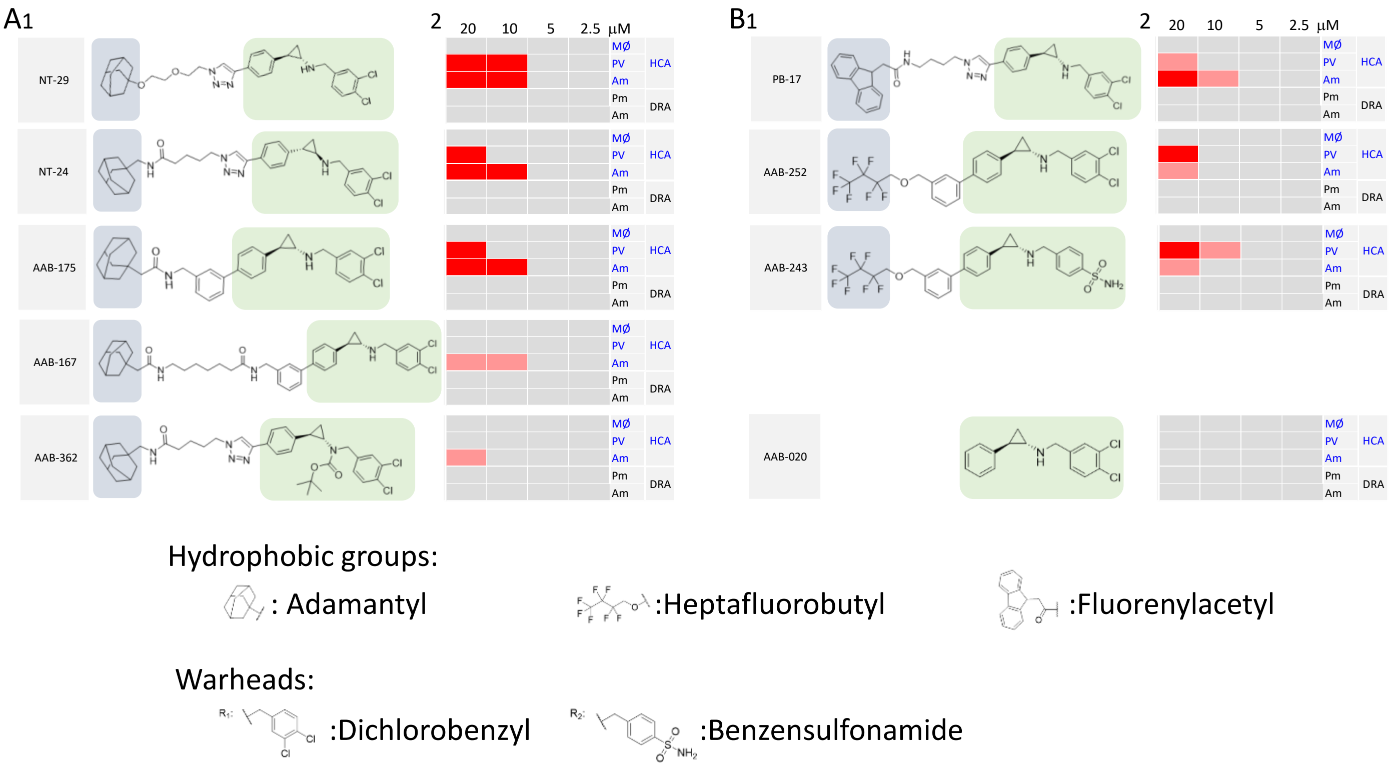


**Figure S10. Gutiérrez-Sanchez M.** ***et al.***

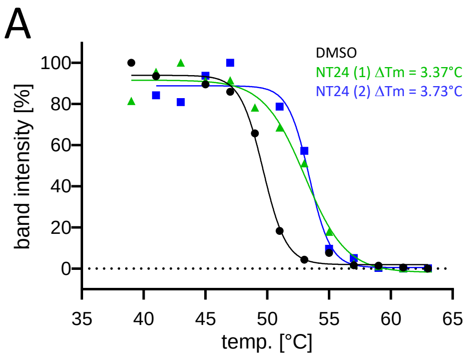


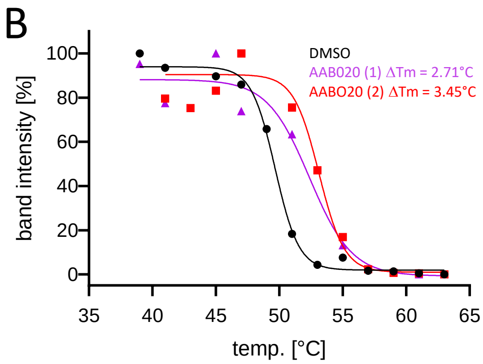




**Figure S11. Gutiérrez-Sanchez M.** ***et al.***

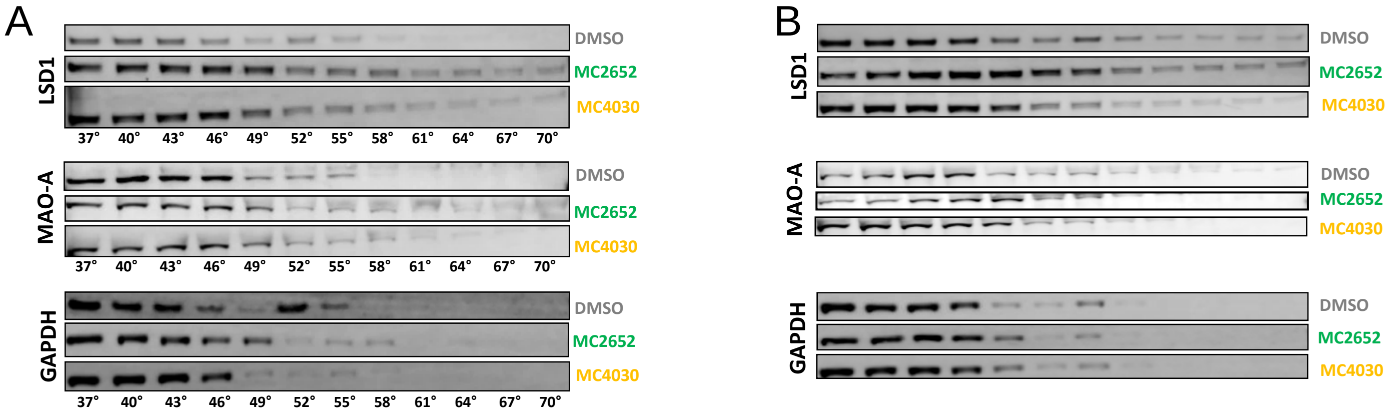


**Figure S12. Gutiérrez-Sanchez M.** ***et al.***

A

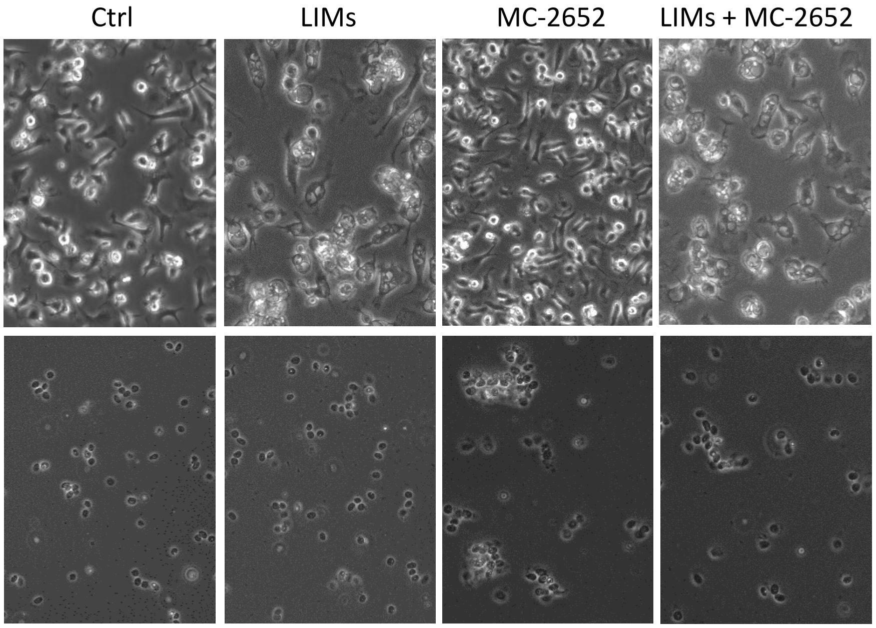


B

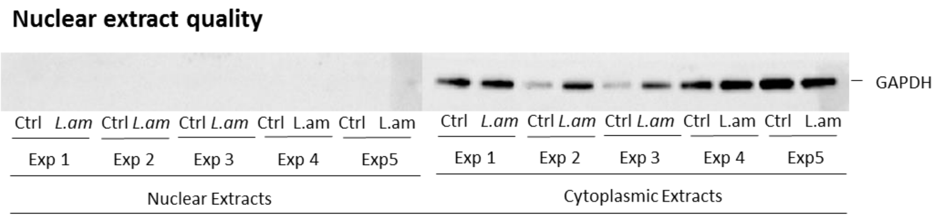


**Figure S13. Gutiérrez-Sanchez M.** ***et al.***
